# Interchain base-stacking and spermine bridges govern ssDNA condensation and reentrant transitions

**DOI:** 10.64898/2026.10.03.756434

**Authors:** Balaka Mondal, Sumit Majumder, Ankur Jain, D. Thirumalai

## Abstract

Polyvalent-cation-induced condensation and reentrant transition in double-stranded DNA are classically described by backbone electrostatics. Whether the same principles govern single-stranded DNA, whose solvent-exposed bases can interact within and between chains to form complex phases, is unknown. Here, we employ simulations and experiments to show that the tetravalent cation spermine (Spm^4+^) drives two equilibrium transitions in poly(dT) and poly(dA). The onset of phase separation, resulting in thermodynamic equilibrium between the dispersed low-density phase and high density coacervate, occurs at a low *sequence independent* Spm^4+^ concentration and is primarily driven by charge balance between phosphate groups and Spm^4+^. Strikingly, the condensed phases are stabilized by at most 1.5 (polydT) and 2.5 (polydA) bridging Spm^4+^ per chain. The ssDNA chains condense to form *fluid-like networks without chain collapse*. The fluidity of chains in the dense phases is confirmed by fluorescence recovery after photobleaching experiments. Both simulations and experiments show that the range of spermine concentration over which phase coexistence persists is *sequence dependent* and nearly threefold wider in poly(dA). At higher *sequence dependent* spermine concentrations, a reentrant phase, characterized by dissolution of the coacervates, emerges. The transition to the reentrant phase occurs when the number of bridging Spm^4+^ molecules vanishes at high sequence dependent spermine concentration. The combination of experiments and simulations show that the two phase transitions in ssDNA are driven by an interplay of stacking and electrostatic interactions.

## Introduction

Polyamines, such as putrescine, spermidine, and spermine, are small aliphatic metabolites that carry multiple amine groups. These groups are predominantly protonated at physiological pH, making them polyvalent cations that bind DNA and RNA [1, 2]. These ubiquitous metabolites occur at millimolar concentrations in cells and shape nucleic-acid folding, compaction, assembly and nucleosome condensability [3–8]. The framework for describing these interactions comes from studies of long double-stranded DNA (dsDNA). Spermine and spermidine condense dsDNA into compact toroids and liquid-crystalline aggregates [3, 9–16]. These transitions are classically explained by electrostatics alone: polyamine accumulation neutralizes phosphate charge, driving a coil-to-globule-like transition [17]; continued accumulation carries the effective charge past neutrality that results in charge inversion [18–20], restoring repulsion and redissolving the assemblies. To a first approximation, this framework treats long dsDNA as a stiff polyelectrolyte of fixed charge density, and its bases are paired and buried within the duplex.

In contrast to dsDNA, the nucleobases in single stranded nucleic acids are solvent exposed. As a result, they can engage in both intra- and intermolecular interactions, through base pairing [21] and stacking. Sticker-spacer model predicted that the interplay between these interactions can produce reentrant transition in aggregation in repeat RNA [22]. ssDNA is also substantially more flexible than dsDNA [23–25]. Its persistence length is nearly fiftyfold smaller, but varies strongly with sequence because base stacking increases chain bending rigidity [26, 27]. The behavior of polyamine-ssDNA complexes is therefore governed by both electrostatic interactions with the phosphate backbone and short-range contacts between exposed bases. These interactions are coupled: sequence-dependent conformations alter the spatial distribution of phosphate backbone charge, whereas polycation association screens electrostatic repulsion and permits closer contact between the bases and chains. Single stranded nucleic acids of identical length and total charge can therefore respond differently to the same ionic environment.

Previous studies showed that short polyamines can induce complex coacervation in single-stranded RNA sequences [28–31]. Bound polyamines screen backbone charge, and a subset simultaneously contacts phosphates on two or more chains, linking them into a connected network. Within these dense phases, nucleic-acid concentrations could exceed the overlap concentration and chains interpenetrate, making intermolecular interactions frequent. Contacts among exposed bases can reinforce that network in a sequence-dependent manner. Indeed, we previously showed that sequence-encoded intermolecular base pairing modulates molecular mobility and material properties in ssDNA and RNA condensates [28]. This interaction landscape is not captured by classical descriptions of dsDNA condensation, in which nucleobase chemistry is largely sequestered within the duplex and phase behaviour is treated primarily in electrostatic terms. Backbone electrostatics and base chemistry have largely been studied separately; how they combine to set the phase behaviour of single-stranded nucleic acids is unknown.

To isolate this coupling, we selected 50-nucleotide homopolymers of thymine and adenine (dT_50_ and dA_50_), as minimal systems. These two single-stranded homopolymers have identical lengths and backbone charge but exhibit sharply contrasting base-stacking propensities: poly(dA) forms strong helical stacks, whereas poly(dT) behaves as a flexible, disordered coil [32–36]. Neither sequence can form canonical Watson-Crick base pairs. Therefore, comparing them allows us to directly calibrate the thermodynamic contribution of base stacking without altering the baseline polymer charge or base complementarity. Using coarse-grained molecular dynamics simulations with the Three Interaction Site (TIS) model for DNA [32, 37, 38] and explicit representation of the tetravalent cation spermine (Spm^4+^), together with *in-vitro* experiments, we map the phase boundaries of these homopolymers. Both polymers undergo reentrant phase separation, forming liquid-like DNA-rich condensates at intermediate spermine concentrations and dissolving at higher concentrations. Although the onset of phase separation is sequence independent and dictated primarily by electrostatics, both the coacervate stability and its eventual reentrant dissolution at high cation concentrations are sequence-dependent: dA persists to nearly threefold higher Spm^4+^ than dT. These condensates are sustained by a sparse network of Spm^4+^ bridges, stabilized by many weak inter-chain stacking contacts. At reentrance, the number of Spm^4+^ bridging different chains drops sharply, whereas the total number of bound Spm^4+^ per chain increases only modestly. Across all three phases, individual-chain conformations remain unchanged, demonstrating a *network without chain collapse*, distinct from the classical condensation of dsDNA.

## Results

### dT_50_ and dA_50_ adopt distinct single-chain conformations that persist across spermine concentrations

Before examining multichain phase behavior, we asked whether spermine alters the conformations of isolated dT_50_ and dA_50_ chains. We performed single-chain low friction Langevin dynamics simulations at *T* = 298 K by varying the ssDNA concentration from 2 *µ*M to 10 *µ*M to calculate the angular correlation function (*C*(*n*)), helicity (*H*), radius of gyration (*R_g_*), and end-to-end (*R_ee_*) distance (Figures 1 and S1). The spermine concentration was varied from 0.2 mM to 20 mM. The structural order was quantified using, *C*(*n*) = ⟨cos *θ_n_*⟩ (Figure 1) where *θ_n_* (Figure 1C) is the angle between phosphate-sugar bond vectors separated by *n* nucleotides along the chain. The simulations show the sequence-dependent contrast between dT_50_ and dA_50_ (Figures 1). At 10 mM Spm^4+^ (purple curve, Figure 1C), *C*(*n*) for dT_50_ decays rapidly with *n*, consistent with a disordered coil structure. In contrast, *C*(*n*) for dA_50_ exhibits a pronounced oscillation with a periodicity of ≈ 10 bases (green curve, Figure 1C), characteristic of a helix stabilized by strong stacking interactions. The mean helical content [39] ⟨*H*⟩, which would be unity for a perfect helix, is ≈ 0.15 for dT_50_ and ≈ 0.7 for dA_50_ (Figure S1). Representative structures are consistent with these observations, showing that dT_50_ populates disordered, coil-like conformations whereas dA_50_ has substantial helical character (see Figure S1) at all spermine concentrations.

**Figure 1:**
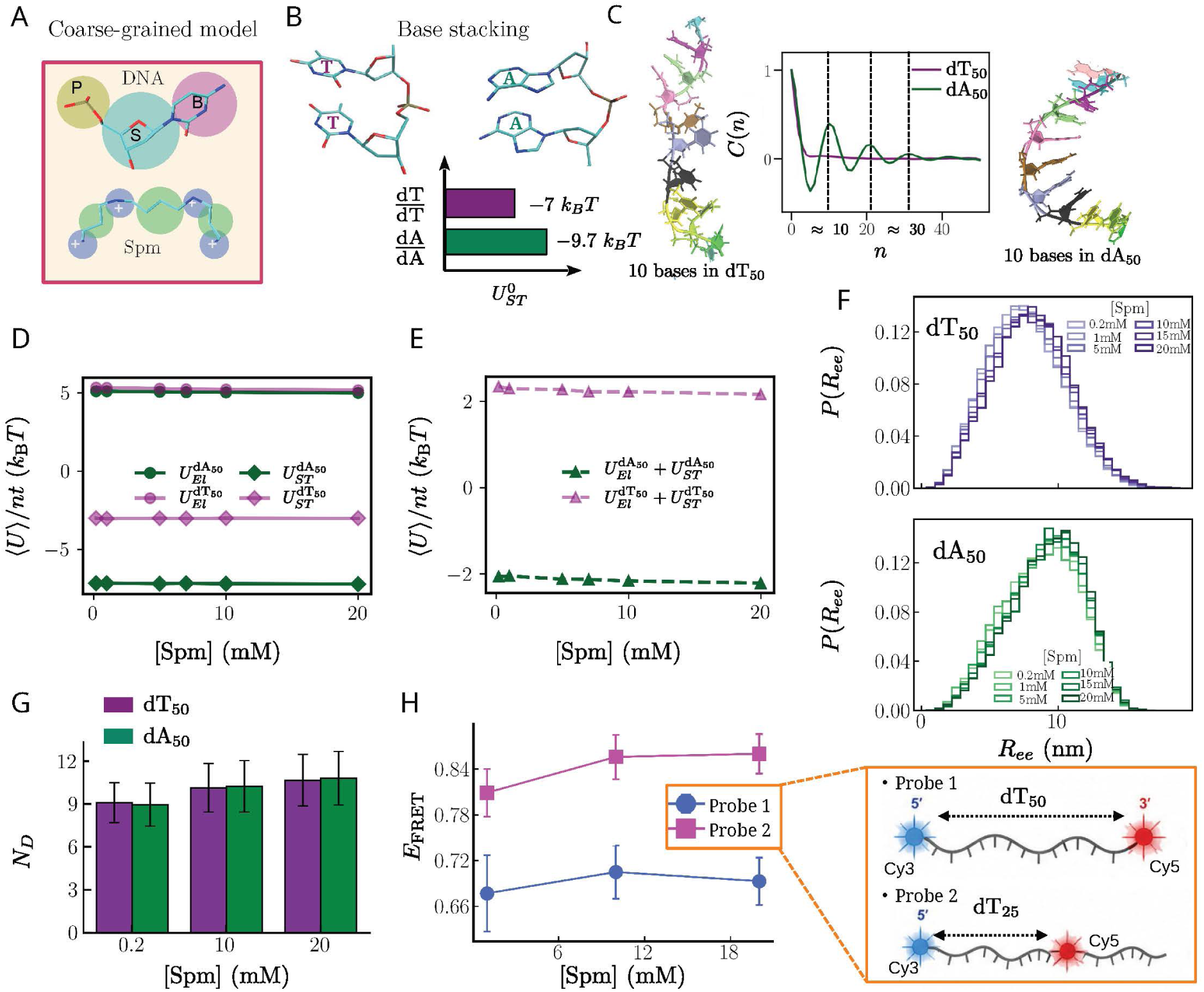
Monomer conformations are independent of spermine concentration: (A) Coarse-grained representations of a single nucleotide and Spermine (Spm^4+^). Each nucleotide is represented using three interaction sites, centered at the center of phosphate (P), sugar (S), and base (B) groups, shown in tan, cyan, and magenta, respectively. Blue beads in Spm^4+^ represent the charged amine groups and the green beads are two types of alkyl groups. (B) Base stacking structures (above) and the associated energies for thymine (dT) and adenine (dA) nucleobases. (C) Tangent correlations as a function of the sequence separation in dT_50_ (red) and dA_50_ (blue) in 10 mM Spm^4+^. The oscillations in dA_50_ are due to the helical structure. Representative conformations of 10 base-long segments from dT_50_ and dA_50_. (D) Potential energy per nucleotide due to electrostatic repulsion between phosphate groups (circles). Stabilizing base stacking interactions including both adjacent and non-adjacent bases (diamonds). (E) Total potential energy contribution to the stability of ssDNA per nucleotide. (F) End-to-end distance (*R_ee_*) distributions at varying Spm^4+^ concentrations. (G) Number of Spm^4+^ molecules, *N_D_*, in direct contact with ssDNA chains. (H) FRET efficiency, *E*_FRET_, measured for dT_50_ (= 2.5 *µ*M), using the two FRET probe constructs shown.

The origin of the contrast between the two sequences is evident from the underlying energetics. Figure 1D decomposes the potential energy per nucleotide into two competing contributions: electrostatic repulsion between backbone phosphates (circles) and stabilizing adjacent and non-adjacent base-stacking interactions (diamonds). The electrostatic contribution is nearly identical for the two sequences, as expected for chains of equal length and backbone charge. The difference between dA_50_ and dT_50_ therefore arises almost entirely from base stacking. As a result, the balance between these contributions changes sign between the sequences (Figure 1E): electrostatic repulsion dominates in dT_50_, yielding *U_EL_* + *U_ST_* ≈ +2.3 *k_B_T* per nucleotide, whereas stacking dominates in dA_50_, yielding ≈ −2 *k_B_T* per nucleotide (Figure 1E). This energetic contrast is consistent with the coil-like ensemble of dT_50_ and the persistent helical order of dA_50_. Neither contribution changes appreciably across the examined Spm^4+^ concentrations (Figure 1D). The nearly invariant electrostatic term follows from the weak concentration dependence of the number of associated cations, *N_D_*, defined as Spm^4+^ molecules with at least one amine group within the Bjerrum length (*l_B_* ≈ 0.7 nm) of a phosphate group. Over the full 100-fold range of Spm^4+^ concentration, *N_D_* rises only modestly and is nearly identical for the two sequences (Figure 1G).

Consistent with this, the single-chain conformational distributions were largely insensitive to increasing Spm^4+^ concentration. The distributions of radius of gyration (*R_g_*), end-to-end distance (*R_ee_*), and the longest contiguous stretches of stacked and unstacked bases, 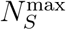 and 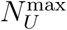 (see SI for definition), varied only weakly (see Figures 1F and S1) despite a 100-fold increase in Spm^4+^ concentration (0.2 mM to 20 mM).

To test the predicted independence of dT_50_ monomer conformation on Spm^4+^ concentration, we designed two dual-labeled FRET constructs. In one construct, Cy3 and Cy5 were positioned at the 5*^′^* and 3*^′^* ends; in the other, Cy3 was positioned at the 5*^′^* end and Cy5 at the chain midpoint (see Figure 1H and experimental methods section for details). For the Cy3-Cy5 pair, Förster radius is *R*_0_ ≈ 5 nm [40]. The end-labeled probe reports on the end-to-end distance whereas the internally labeled probe reports on the distance between one end and the chain midpoint. For both constructs, the apparent FRET efficiency, *E*_FRET_, is essentially unchanged within experimental uncertainty between 1 mM and 20 mM Spm^4+^ (2.5 *µ*M probe and 10 mM Tris pH 7.0 at *T* = 295 K; Figure (1H) . These results provide direct experimental evidence that neither the terminal nor the end-to-midpoint separation undergoes a large spermine-dependent change. Together, these results show that dT_50_ remains coil-like and dA_50_ remains helically stacked across the Spm^4+^ titration.

### Spermine drives phase separation in homopolymeric ssDNA sequences

Having established the single-chain baseline (Figure 1), we asked how spermine affects interactions between chains. A previous experimental study established that homopolymeric dT_90_ phase separates with Spm^4+^ [28]. To further probe the effects of sequence on the equilibrium phase changes in dT_50_ and dA_50_, we performed multi chain simulations (at 350 *µ*M DNA concentration) as a function of Spm^4+^ concentration. Both the sequences start phase separating at 4mM Spm^4+^ concentration (Figure 2A-D) into a DNA rich condensate phase and DNA depleted dilute phase. We ascertained that phase separation in the simulations occurs at equilibrium by verifying the equality of the chemical potentials [41] of the DNA chains and Spm^4+^ in the two phases (Figures S2 and S3). Phase separation was monitored by counting the number of ssDNA chains clustered together (see Figure S4 for cluster definition and SI for the algorithm details). The radial concentration profiles showed strong enrichment of the phosphate groups (*P*) and amine groups of Spm^4+^ within the condensate (≈ 500 fold relative to the dilute phase) whereas Cl*^−^* concentration was comparable between the two phases (Figures 2B,D). The nearly identical profiles of phosphate and amine-groups are consistent with electrostatic association between the oppositely charged components, leaving the condensate interior charge neutral. Thus, phase separation emerged directly from the molecular interactions encoded in the model rather than being imposed through the initial configuration.

**Figure 2:**
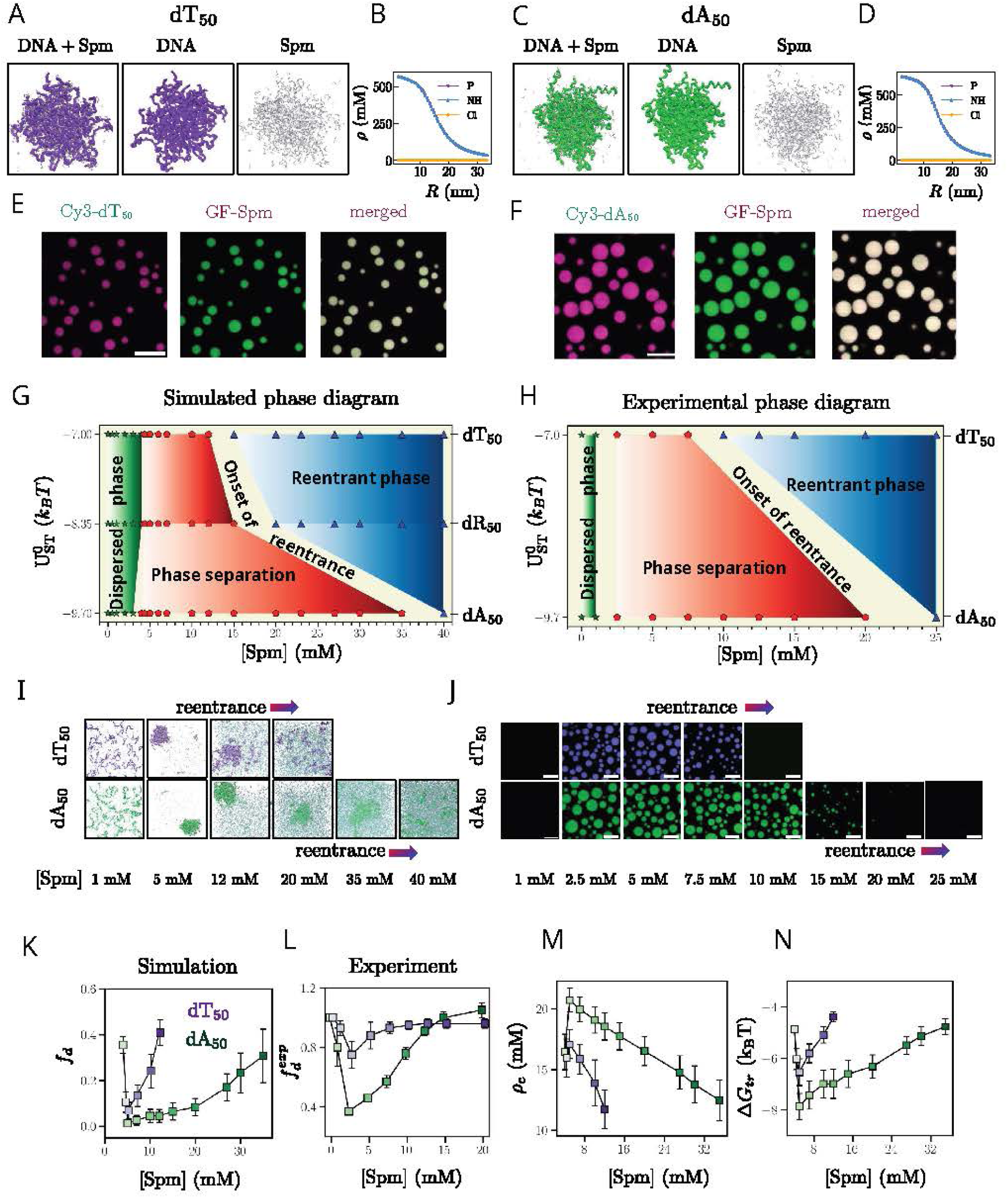
Onset of phase separation is sequence independent while reentrance is sequence dependent: (A, C) Simulation snapshots illustrating the accumulation of ssDNA and spermine in the condensate phases. dT_50_ (dA_50_) chains are in purple (green); Spm^4+^ molecules are shown in silver. (B, D) Concentration (*ρ*) profiles of phosphate (*P*), spermine (*NH*) and chloride (Cl*^−^*) ions as a function of the distance from the droplet center, *R*, for dT_50_ and dA_50_, respectively. (E, F) Fluorescent micrographs showing phase-stability of ssDNA-spermine condensates at [Spm^4+^] = 2.5mM at the DNA concentration of 50 *µ*M. Scale bars correspond 10 *µ*m. (G) Simulated phase diagram in the plane of stacking interactions and spermine concentration. At low spermine concentrations (up to 4mM), all three sequences are in the dispersed phase. At high spermine concentrations, dT_50_, dA_50_ and the mutant sequence dR_50_ undergo reentrant phase transition from a condensed to a new phase. (H) Experimental phase diagram also show that there are two transitions that are qualitatively similar to the simulation results. (I) Representative simulation snapshots illustrating the transition from a dispersed phase, to phase separated phase, and then to the re-entrant dispersed phase in of dT_50_ (top) and dA_50_ (bottom). (J) Fluorescent micrographs showing phase-stability of DNA-spermine condensates as a function of Spm^4+^ concentrations. Experiments were performed at room temperature and at a fixed DNA concentration of 75 *µ*M. (K) Fraction of DNA chains in the dilute phase, *f_d_*, as a function of spermine concentration for dT_50_ and dA_50_. (L) Dependence of the fraction of ssDNA chains, 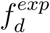, measured by nanodrop spectrophotometry at a fixed DNA concentration of ≈ 50 *µ*M. (M) Monomer concentrations in the condensate phase, *ρ_c_*. (N) Transfer free energy, 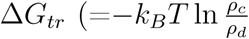, where *ρ_d_* is the monomer concentrations in the dilute phase). In (K–N), dT_50_ and dA_50_ are shown using the same color code.

To characterize the condensates formed by these two sequences experimentally, we mixed dilute solutions of dT_50_ or dA_50_ (≈75 *µ*M) with Spm^4+^ (2.5 mM, 10 mM Tris pH 7.0, at *T* = 295 K). A small fraction of fluorescently labeled Spm^4+^ (1 *µ*M) and the corresponding DNA (40 nM) were included to visualize the condensates by fluorescence microscopy. Immediately upon mixing, the system separated into a dense, macromolecule-enriched coacervate phase and a dilute supernatant phase, with DNA and Spm^4+^ strongly enriched within the condensates (Figures 2E and F). Having established that simulations and experiments both produce coacervates under comparable conditions, we next determined how sequence-specific interactions set the phase boundaries.

### Onset of phase separation is sequence independent while reentrance is sequence dependent

To elucidate sequence effects, we determined the changes in the phase behavior as a function of stacking strength, 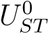, and spermine concentration, and tested the predictions experimentally. To assess the importance of 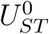, we also simulated a “synthetic” sequence, dR_50_, with an intermediate stacking strength (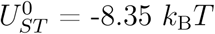, compared with -7 *k*_B_*T* for dT and -9.70 *k*_B_*T* for dA). The resulting phase diagram (Figures 2G and I) reveals three features, each of which we validated using experiments performed over a range of DNA concentrations (25 − 75 *µ*M, 10 *mM* Tris pH 7.0 at *T* = 295 K) while varying Spm^4+^ (Figures 2H and J).

First, the onset of phase separation is relatively *sequence-independent* and is primarily determined by electrostatic interactions. Simulations show that the saturation concentration, *C*_sat_, the minimum Spm^4+^ concentration at which condensation is initiated, is nearly identical (≈ 4 mM) for the three sequences. The value of *C*_sat_ coincides with the spermine concentration at which the phosphate and the charges on spermine are balanced. Experiments showed the same limited sequence dependence: the onset lay between 1 and 2.5 mM Spm^4+^ for both dT_50_ and dA_50_ at = 50 *µ*M and = 75 *µ*M DNA. Thus, both sequences enter the two-phase regime at similar spermine concentration, despite having substantially different stacking propensities, indicating that stacking has only a limited effect on the lower phase boundary.

Second, in contrast to the onset, the spermine concentration at which the condensates undergo reentrant dissolution depends strongly on the DNA sequence. The Spm^4+^ concentration range over which dT_50_ phase separation occurs is narrower than in dA_50_. Simulations predict phase separation over [Spm^4+^] ≈ 4−35 mM for dA_50_. This range narrows to ≈ 4−15 mM for dR_50_ and to ≈ 4−12 mM for dT_50_. Experiments show a similar sequence dependence. dT_50_ condensates are stable only over a narrow range (≈ 2.5−7.5 mM at 75 *µ*M DNA). In contrast, dA_50_ condensates persist over a much broader range (≈ 2.5−20 mM) (Figure 2H). The same ordering was found at the lower DNA concentrations examined, with the dissolution boundary shifting to lower spermine as the DNA concentration was reduced. Expressed as nominal amine charges per phosphate, the dissolution boundary is: ≈8 for dT_50_ and ≈ 21-24 for dA_50_ at both DNA concentrations. The ratio of 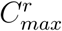 (the highest Spm^4+^ concentration up to which the two-phase regime persists) between dA_50_ and dT_50_ is ≈ 2.9 in simulations and ≈ 2.7 − 3.0 in experiments (compare Figure 2G and H), even though the DNA concentration in simulations (350 *µ*M) is higher than in experiments (50-75 *µ*M).

Third, the propensity for phase separation is maximum at 5 mM Spm^4+^ (*> C*_sat_) in both dT_50_ and dA_50_. Figure 2K shows that the fraction of ssDNA chains in the dilute phase, *f_d_*, reaches a minimum at [Spm^4+^] ≈ 5 mM in a sequence independent manner. Strikingly, the experimentally measured soluble DNA fraction 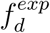 for both DNAs follows a similar non-monotonic profile, reaching a minimum at [Spm^4+^] ≈ 2.5 mM with dA_50_ partitioning more strongly than dT_50_ throughout the examined range (Figure 2L). A physically transparent measure of this trend is the transfer free energy, 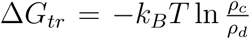, calculated from the simulations (Figure 2N), where *ρ_c_* and *ρ_d_* are the monomer densities in the condensate and dilute phases, respectively. Δ*G_tr_* is more favorable for dA_50_ than for dT_50_ over the entire Spm^4+^ range. It reaches a minimum near 5 mM in both sequences. This is consistent with the narrower distributions of condensate and dilute-phase densities found for dA_50_ relative to dT_50_ (Figures S5−S6, Table S1).

The condensate-phase densities from simulation (see Figure 2M) and experiment fall in the same regime, but the agreement is only semi-quantitative. At 2.5 mM spermine and 50*µ*M DNA, experiments give *ρ_c_* ≈ 58 ± 15 mM for dT_50_ and 74 ± 7 mM for dA_50_. Simulations ([DNA] = 350 *µ*M) at 5 mM Spm^4+^ (*N* = 160) give *ρ_c_* ≈ 21 ± 0.6 mM and 23 ± 0.7 mM, respectively. We surmise that this discrepancy reflects the coarse-grained nature of the model, finite-size effects (see Figure S7), and differences in the initial DNA concentration (Figure S8). In addition, fluorescence-based concentration estimates also carry inherent uncertainty, since they are sensitive to probe partitioning, dye quantum yield, and label-induced perturbations (see SI for details). Nevertheless, both simulations and experiments place the coacervate phase at millimolar DNA concentration with a nominally higher density for dA_50_ than for dT_50_. Taken together, the onset of phase separation is governed by electrostatics and is insensitive to sequence, whereas the width of the two-phase regime and the reentrant boundary depend on sequence-specific stacking.

### Forces driving phase separation

The phase diagrams (Figures 2G and H) show that two major competing factors control Spm^4+^ induced phase transitions. One is the favorable sequence dependent hydrophobic interactions arising from stacking of bases. The other is the chain electrostatic interaction, which remains repulsive even after condensation of Spm^4+^ due to incomplete neutralization of the charges on the phosphate groups. Unlike in rod-like polyelectrolytes [42] the effective charge, *q_eff_*, on each phosphate in the flexible ssDNA is difficult to determine. A number of factors, such as the gain in the favorable electrostatic interactions due to condensed counterions, persistence length of the ssDNA chains that depends on the strength of stacking interactions [43], fluctuations in the condensed and free counterions [44–46] contribute to *q_eff_* . An approximate estimate yields *q_eff_* ≈ -0.5*e* after spermine condensation. Because the residual repulsion is substantial, coacervates can form only if it is offset by attractive stacking interactions. It also follows that the drive to form coacervates should be greater in dA_50_ than in dT_50_, which is demonstrated in experiments (Figures 2H and L).

In order to dissect the roles of the two competing interactions, we performed a series of simulations for the two sequences at 5mM Spm^4+^ to isolate the forces that drive phase separation. (1) To assess if electrostatic interactions and charge fluctuation-induced attraction [18, 19] are sufficient, as is the case in dsDNA, we set the inter or intra (Figure 3B) stacking interactions to zero. We performed multi-chain simulations starting from an initial conformation consisting of a preformed droplet that is in equilibrium with the dispersed phase. In the absence of stacking (both neighboring and non-neighboring) interactions, the dense phases rapidly become unstable (Figures 3B and D). The angular correlation function, ⟨*cos*(*θ_ij_*)⟩ decays rapidly as |*i*−*j*| increases (Figure 3C), which differs from the result for dA_50_. Thus, to form stable droplets sequence specific stacking interactions are needed. (2) Next, we eliminated the weak intra and inter chain non-neighboring (NN) stacking interactions. Neighboring stacks and electrostatic interactions were kept intact (Figure 3D). Multi-chain simulations, starting from an equilibrated droplet, show that the dense phase is destabilized in the absence of NN stacking (Figures 3D). We conclude that the weak NN stacking, even though it contributes negligibly to monomer conformations (see Figures 3E, and S9-S11), are nonetheless required to stabilize the condensate. (3) Finally, we eliminated all the electrostatic interactions by neutralizing the charges on the DNA phosphates, Spm^4+^ and Cl*^−^* ions, while retaining only the neighboring and non-neighboring stacking interactions. Under these conditions, there is an *irreversible* “hydrophobic” collapse, in which nearly all the ssDNA chains condense into a compact structures in both dA_50_ and dT_50_ (Figures 3F-G). The droplets are hyper stable, which is different from the reversible equilibrium transition in the dT_50_ and dA_50_ condensates. The results in Figures 2G and H, and Figure 3 establish that both electrostatic and stacking interactions are crucial in driving phase separation in dT_50_ and dA_50_.

**Figure 3:**
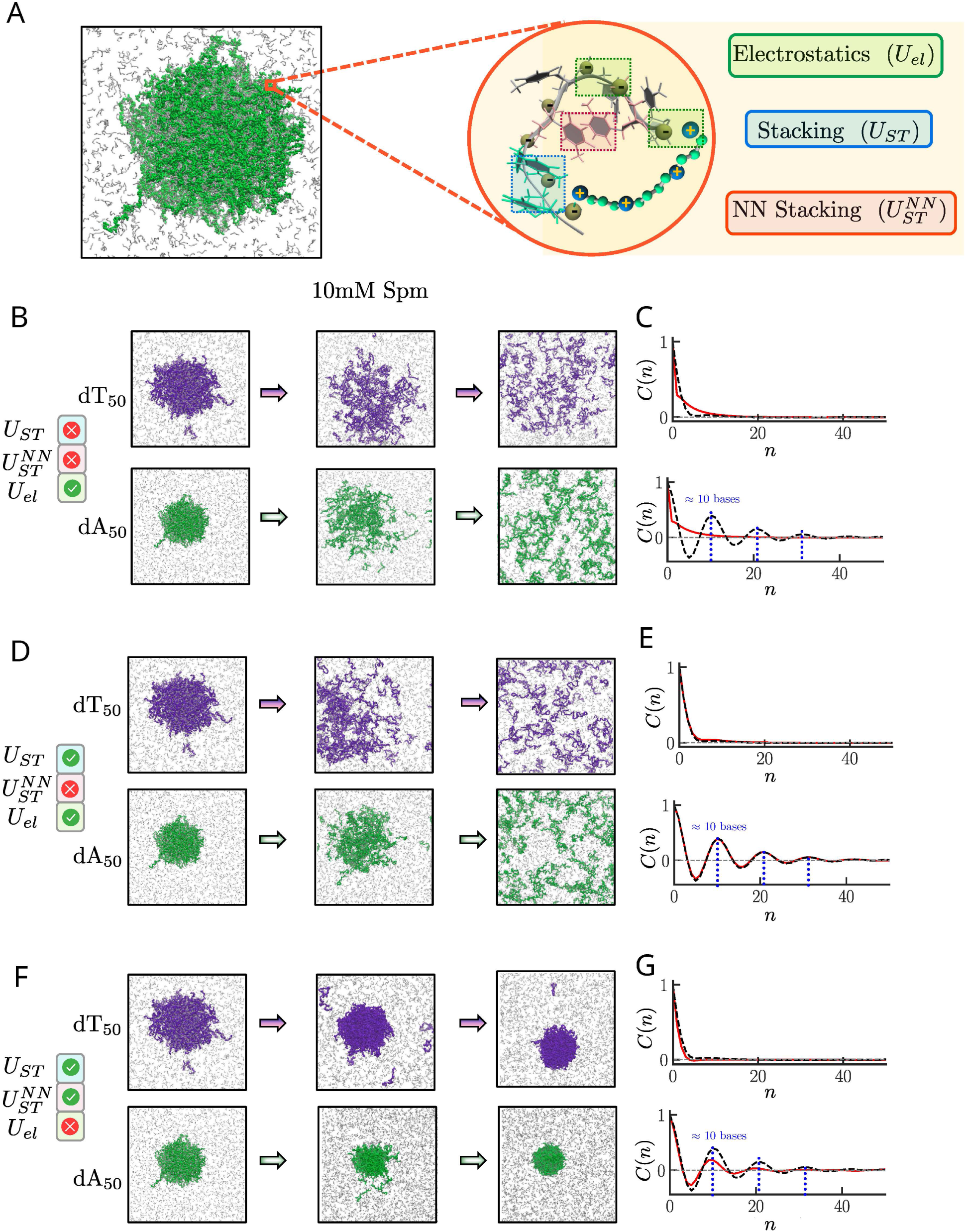
Interplay of electrostatics and base stacking in phase separation. (A) Illustration of the major contributing forces in DNA/Spm^4+^ condensate formation. (B) Stages of phase separation dynamics in multi-chain (*N* = 160) simulations, initiated from a preformed droplet with both adjacent and non-neighboring stacking interactions disabled. (C) Angular correlation function, along the sugar-phosphate backbone of ssDNA. Red line indicates the modified system with specified interactions disabled, and black dashed line corresponds to the unmodified reference system. Blue dashed lines show periodicity of ≈ 10 bases in unmodified dA_50_ constructs. Same as (D), but with only non-neighboring stacking interactions disabled. (E) Corresponding angular correlation functions. (F) Same as (B) and (D), but with only electrostatic interactions disabled. (G) Corresponding angular correlation functions. All the results are shown in 10 mM Spm^4+^ . ssDNA chains are in purple (dT_50_) and green (dA_50_). Spm^4+^ molecules are in silver.

### Loss of bridging interactions drive transition to the reentrant phase

Analysis of the chain conformations within the condensate shows that ssDNA chains interact predominantly through spermine-mediated bridging interactions (see below). In this mechanism, a single Spm^4+^ cation simultaneously associates with several DNA chains through phosphate–amine electrostatic interactions (Figure 4A). The number of bridging ions per DNA chain, which increases during the transition from the dispersed to the condensate phase, decreases precipitously with increasing spermine concentration in both sequences (Figures 4B) and C). Crucially, this decrease is not caused by spermine dissociating from the DNA: over the same range, the total number of DNA-associated spermine molecules per chain increases. At high spermine concentration, there is a sharp transition from interchain bridging to predominantly non-bridging interactions (see Figures S12), resulting in the formation of the reentrant phase.

**Figure 4:**
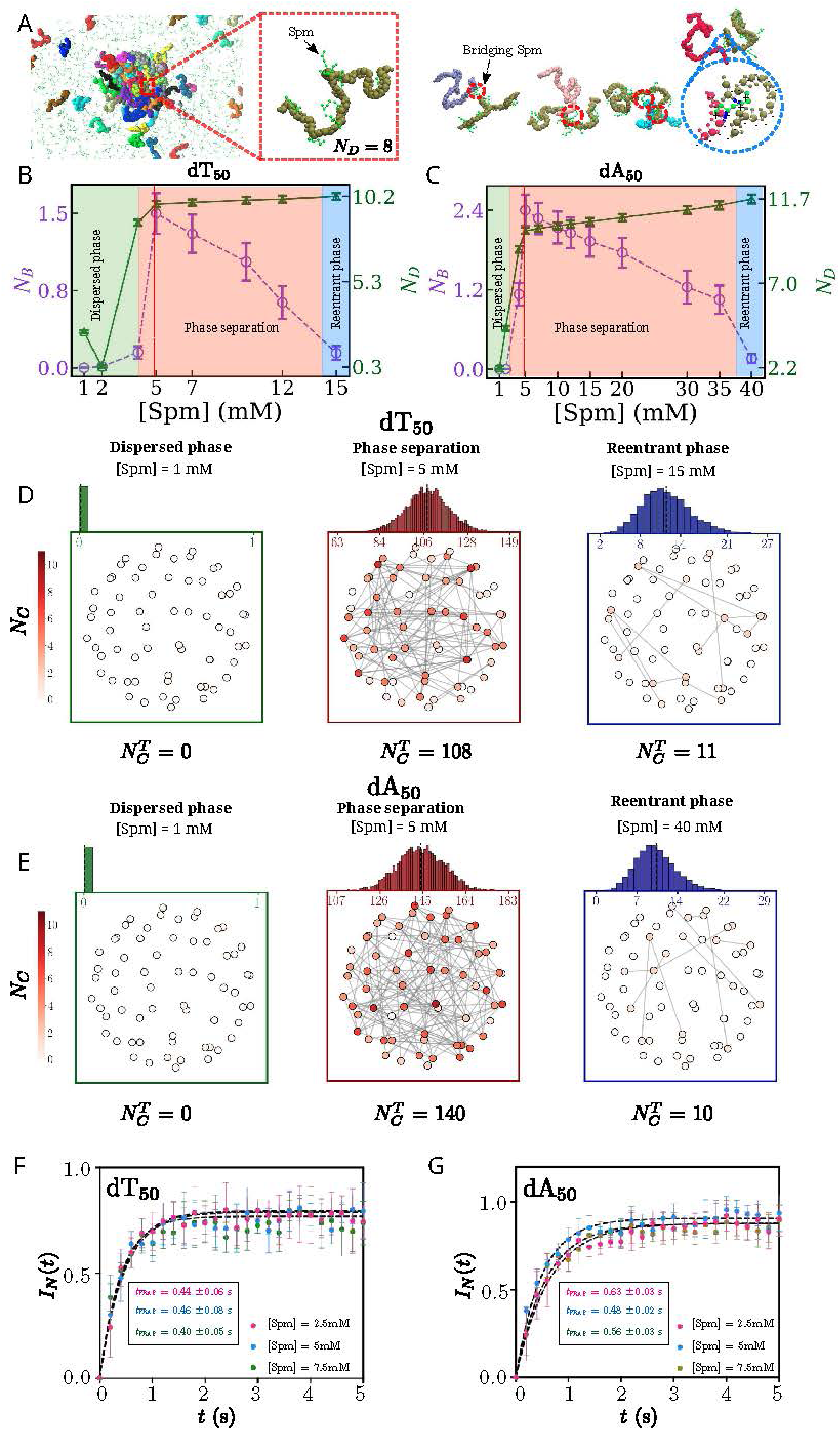
Loss of spermine-mediated bridging drives reentrant transition. (A) DNA chains in the condensates interact via Spm^4+^ -mediated bridging. A highlighted DNA chain (olive) forms bridging contacts with four others. (B) Average number of bridging spermine molecules, *N_B_* (scale is on the right), and the average number of decorating spermine molecules, *N_D_* (scale is on the right), per dT_50_ chain as a function of Spm^4+^ concentration. Phases are highlighted in green (dispersed), red (condensates) and blue (reentrant) shaded regions. Red dashed line shows the spermine concentration at which *N_B_* is a maximum. (C) Same as (B), except the results are for dA_50_. (D) Network representations of dT_50_ connectivity in the dispersed (left), condensed (middle), and reentrant (right) phases. Each node represents a single DNA chain. Gray lines are the inter-chain connections mediated by spermine bridging. Network between different DNA chains is visualized on a Fibonacci sphere [64]. Node positions do not correspond to actual spatial coordinates of the system but are arranged on the sphere solely to aid visualization of inter-chain connectivity. Color bar represents number of connections (*N_C_*) per DNA chain. Total number of connections is 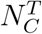. The top panel shows the distribution of 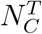 per frame, with the dashed vertical line marking the mean. For each phase, the network shown is for the frame with 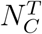 closest to the mean. (E) Same as (D), except the results are shown for dA_50_. (F) Fluorescence recovery after photobleaching (FRAP) kinetics of dT_50_ chains and spermine within the condensates. Recovery times t_FRAP_ are shown in order of increasing spermine concentration. (G) Same as (F), but for dA_50_ condensates.

The decrease in the number of bridging interactions reduces the effective connectivity of the condensate network (see Figures 4D and E). Once it falls below the connectivity threshold needed to sustain a stable network, the condensate is destabilized. Interestingly, reentrant transition in both sequences occurs when the number of bridging spermine per DNA falls below ∼ 0.5. The onset of reentrant transition at high spermine concentration identifies a *sequence-independent* connectivity criterion that governs the transition from a networked condensate to a sparsely connected state.

Figures 4B–E reveal that the condensates form a dense network. The dA_50_ network has a higher number of bridging spermine molecules (*N_B_*) and total inter-DNA connections 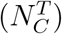 compared to dT_50_. The enhanced connectivity stems from inter-strand non-neighboring stacking interactions, 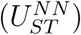. At 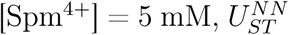 per dA_50_ chain is nearly twice compared to dT_50_ (Figures S13B and E), underscoring the role of sequence-dependent stacking in stabilizing the condensate networks.

In dA_50_, strong base stacking drives chains into a more densely packed arrangement, which enhances the probability that a single Spm^4+^ ion can bridge multiple backbones. Thus, even at similar overall spermine concentration, bridging is more pronounced in dA_50_. Notably, both sequences exhibit the same qualitative relationship between inter-chain stacking energy and the number of bridging interactions as a function of spermine concentration. This indicates that stacking and ion-mediated bridging act cooperatively to determine condensate connectivity.

Fluorescence recovery after photobleaching (FRAP) experiments (Figures 4F and G) on dT_50_ and dA_50_ condensates report nearly [Spm^4+^]-independent recovery times (*t*_FRAP_). This behavior is consistent with our finding that, within the phase-separated regime, ss-DNA chains are connected through a sparse and dynamic network of spermine-mediated bridges, which render the condensates to be fluid-like. Across the range of [Spm^4+^] that supports condensation, the average number of bridging ions per chain varies only modestly (see Figures 4B and C) and remains low. Even at the highest phase-separation propensity ([Spm^4+^] = 5 mM), each chain is associated with only ∼ 1.5 (dT_50_) to ∼ 2.5 (dA_50_) bridging Spm^4+^ molecules. This level of connectivity is sufficient to maintain condensate integrity while maintaining a fluid-like behavior. Consequently, chain mobility and hence *t*_FRAP_ are governed by transient and modest network rearrangements, and do not change significantly with [Spm^4+^]. These observations indicate that dT_50_ and dA_50_ condensates remain in a liquid-like, weakly crosslinked regime throughout the phase-separated [Spm^4+^] range.

### Condensate growth mechanism

At [Spm^4+^] = 10 mM, both dT_50_ and dA_50_ phase separate into a DNA-rich phase and a DNA-poor phase (Figures 5A and B). Despite forming spherical droplets at long times, the two sequences exhibit strikingly different growth mechanisms (see Supplementary Videos 1 and 2). Condensate growth of dT_50_ droplets proceeds predominantly through addition of a monomer at a time to a growing cluster (Figures 5A). The size of the largest cluster, *N* (*t*), increases steadily with time (Figure 5C), while the corresponding growth rate, *dN* (*t*)*/dt*, remains approximately constant. These findings show that dT_50_ droplets grow by the sequential addition of individual chains throughout the phase-separation process (Figure 5C).

**Figure 5:**
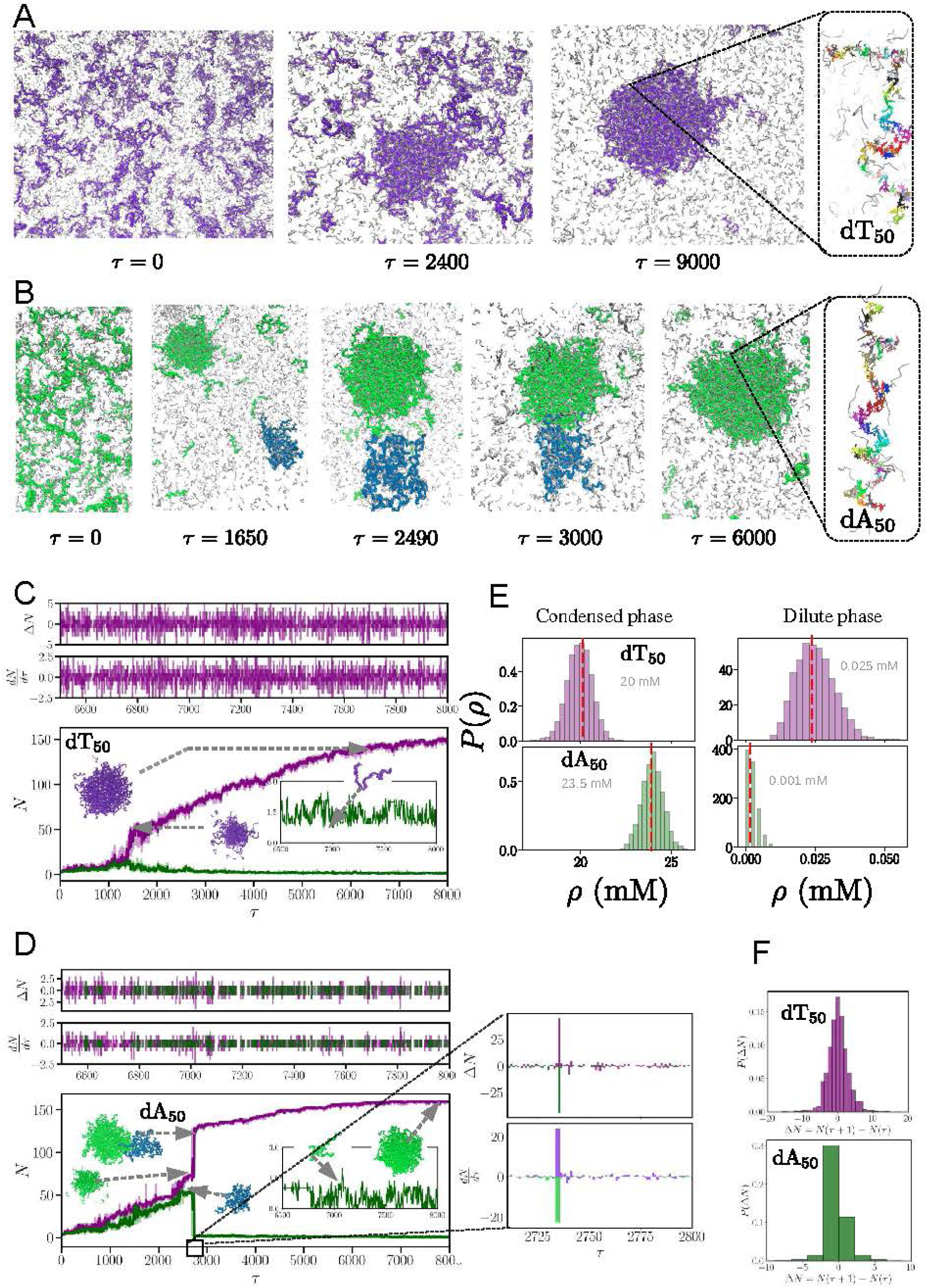
Sequence-dependent condensate growth: (A) Stages in the dynamics of phase separation dynamics in multi-chain (*N* = 160) simulations of dT_50_ in 10mM Spm^4+^ at *T* = 298 K. (B) Same as (A), except the results are for dA_50_. (C) Time evolution of the two largest clusters in dT_50_. Purple curve shows that the number of chains in the condensate increases monotonically with time. Change in the number of chains (Δ*N*) and rate of change 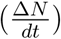 are constant (see the upper panels). (D) Growth of the dA_50_ condensate (purple) as a function of time. At times less than ≈ 3, 000*τ* two small droplets (green and blue) form. Upon fusion of the green and blue droplets at *t* ≈ 3, 000*τ* the larger (green) droplet size increases at the expense of the smaller (blue) one. Fusion event is reflected in Δ*N* and 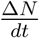 (upper panel), which show an abrupt change over a narrow time interval (2730 *τ* to 2740 *τ*). Upon fusion, a large spherical droplet emerges at long times (inset). (E) Distributions of monomer densities in the condensate and dilute phases. The distributions in both the phases are broader in dT_50_ relative to dA_50_. (F) Distributions in the change in the number of chains Δ*N* in the largest cluster between consecutive times.

In contrast, dA_50_ condensates grow through hierarchical coalescence. Small oligomers formed at early times, dissolve and are progressively absorbed into larger droplets (see also the videos in the SI), a growth mechanism that is reminiscent of Ostwald ripening [47–50]. For instance, at *τ* ≈ 2.5 × 10^3^, two oligomers fuse to generate a condensate containing approximately 130 chains, corresponding to nearly 80% of the system (Figure 5D). The fusion event produces a discrete, step-like increase in *N* (*t*) and a sharp transient increase in *dN* (*t*)*/dt*, in marked contrast to the nearly linear growth observed of dT_50_ droplet (Figure 5C). After the major fusion event, further growth occurs primarily through monomer addition to the pre-existing condensate (Fig. 5F).

Thus, sequence determines not only the stability of the condensed phase but also the growth mechanism. The assembly of dT_50_ occurs through gradual addition of one chain-by-chain at a time, whereas dA_50_ forms through the merger of metastable oligomers into a condensate. The newly formed dA_50_ condensates are initially highly nonspherical but relax to nearly spherical droplets at long times, to minimize the interfacial free energy (see Figure S14). Similar findings are found in other trajectories (see Figure S15 in the SI). Because multiple independent nuclei can form only when the simulation box contains enough chains, oligomer fusion is observed in larger systems (*N* = 160). At present, the early stages of growth are difficult to resolve experimentally. Moreover, possible shear stresses generated during mixing and sample loading may enhance droplet encounters and coalescence relative to purely diffusion-driven assembly. The well calibrated simulations, therefore, provide a molecular-level view of sequence-dependent assembly that is currently not readily accessible in experiments.

### Monomer conformations are unchanged in the three phases

Strikingly, the monomer conformations are virtually unchanged in the dispersed, condensed and reentrant phases in both the sequences (Figures 6 and Figure S16). For instance, the distributions of *R_g_*, *R_ee_*, as well as the dependence of *C_n_* on |*i* − *j*|, are virtually identical (Figures 6 and S16). FRET measurements on dT_50_ support this conclusion experimentally. Using both the terminal (term-dT_50_) and internal (int-dT_50_) probe constructs, we find that *E*_FRET_ is unchanged within experimental uncertainty between the dispersed (1 mM Spm^4+^) and reentrant (20 mM Spm) phases: 0.69 ± 0.03 vs. 0.72 ± 0.1 for term-dT_50_, and 0.86 ± 0.1 vs. 0.86 ± 0.1 for int-dT_50_. Thus, neither the end-to-end nor the end to chain midpoint separation changes measurably across a twenty-fold change in spermine concentration.

**Figure 6:**
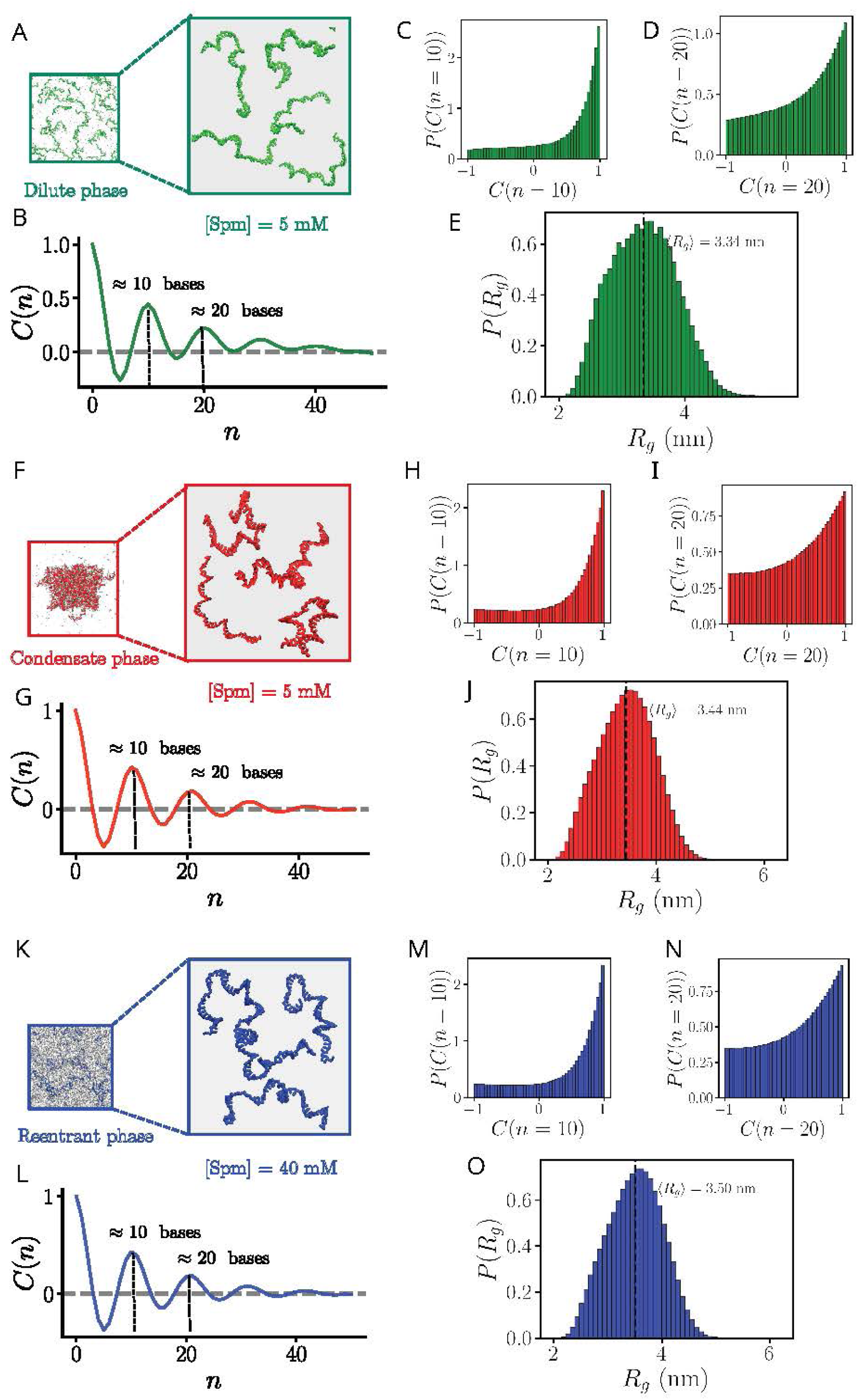
dA_50_ monomer conformations in the three phases are similar: (A) A snapshot showing the conformation in the dilute phase of dA_50_ in 5mM Spm^4+^. Few representative conformations are highlighted (in green). Spm^4+^ molecules are in silver. (B) Angular correlation function, *C*(*n*), as a function of inter-residue separation, *n*, averaged over the conformations in the dilute phase. (C, D) Distributions of *C*(*n*) at *n* = 10 and *n* = 20, corresponding to the first and second peaks in Figure 6B. (E) Distributions *P* (*R_g_*)s of radius of gyration *R_g_* of monomers in the dilute phase. The dashed lines are average values of *R_g_* obtained from the distributions. (F-J) Same as (A-E), except the results correspond to condensate phase ([Spm^4+^] = 5mM). (K-O) same as (A-E), except the results are for the reentrant phase ([Spm^4+^] = 40mM). The Jensen-Shannon divergence (JSD) between pairs of *C*(*n*) distributions from the three phases is approximately zero.

The FRET measurements could not be extended to the condensed phase. The high local chain density in the coacervate mixes intra- and inter-chain donor-acceptor signals, precluding a well-defined *R*_Cy3_*_−_*_Cy5_. Simulations and the available experiments point to the same conclusion that chain conformation is decoupled from the phase state.

The invariance of monomer conformations in three phases is surprising because there are substantial substantial changes in the interactions between two or more chains mediated by spermine during the dispersed → condensed → reentrant phase transitions (Fig. 5). Furthermore, Fig. 4 shows that the stability of the condensate phase depends on the inter-chain non-neighboring (NN) stacking interactions, which also changes during the phase transition (Figure S13). Nevertheless, the conformations of the individual chains are virtually unaltered during the phase changes.

There are a few factors that determine the fate of the monomers in the three phases. (1) The spermine dependent bridging network that undergoes large changes during the transition between the three phases (Fig. 4). At the spermine concentration of 5 mM, *N_B_* (number of Spm^4+^-mediated bridges per DNA chain) reaches a maximum of approximately 1.5 for dT_50_ and 2.5 for dA_50_ (see Figures 4B and C). The Spm^4+^-mediated inter-chain bridge involves two phosphate (*P*) groups from different chains. Thus, these values correspond to 1.5 phosphate groups per dT_50_ chain and 2.5 phosphate groups per dA_50_ chain, which represent only ∼3% of the total phosphates in dT_50_ and ∼5% in dA_50_. (2) At neutral pH, spermine is a short tetravalent cation with an average end-to-end distance (⟨*R_ee_*⟩) of ∼1.5 nm (see Figure S4A). In contrast, ⟨*R_ee_*⟩ of the dT_50_ and dA_50_ (Figures S17) are a few times larger than the range of a single Spm^4+^ bridge. The inter-chain bridge formation requires only local fluctuations that bring two nearby phosphate groups within ∼1.5 nm of one another. The free energy cost to bend of the DNA chains is small because the persistence length of ssDNA is only ∼1 nm, comparable to the span of spermine. Consequently, spermine-mediated bridging can occur without substantial bending or large-scale rearrangement of the chain conformations. (3) It is possible that structural change in the monomer could arise due to favorable interchain NN stacking, which stabilizes the condensate. However, the simulations show that the intra-chain stacking energy, 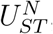, is overwhelmingly greater than inter-chain contribution in the three phases (Figures 7C-D, and S13). In the dispersed regime, 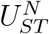 is approximately 200-fold greater than 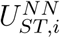 for dT_50_ and nearly 500-fold stronger for dA_50_. In the condensed phase, where intermolecular contacts are most abundant, intra-chain stacking is 40–80 times stronger in dT_50_ and approximately 100 times stronger in dA_50_. In the reentrant phase, the disparity increases to roughly 200-fold and 350-fold, respectively for the two sequences. We surmise that intermolecular stacking is too weak to disrupt the pre-existing single-chain organization.

**Figure 7:**
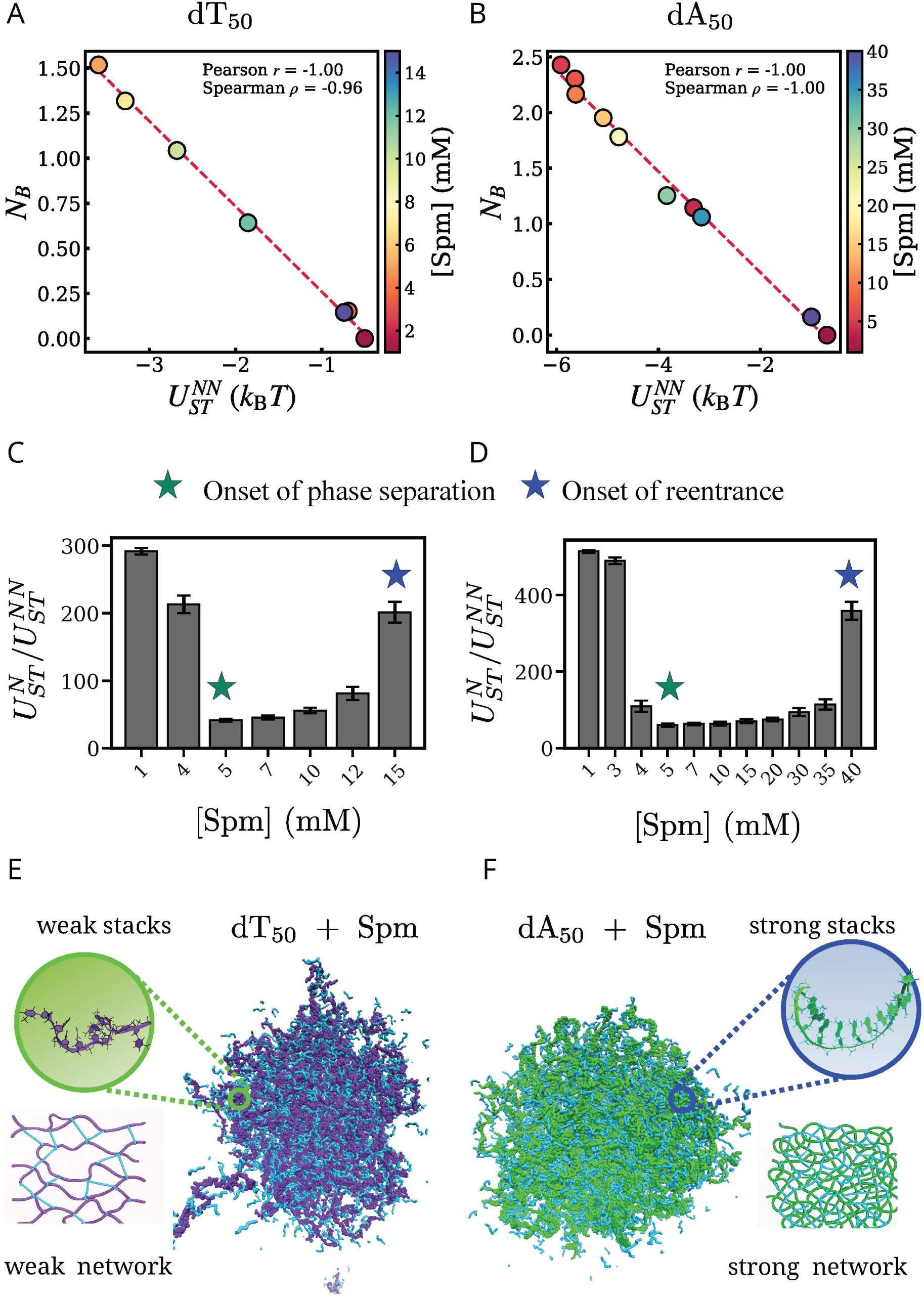
Bridging correlates with favorable inter-strand stacking: Correlation between the mean number of spermine-mediated bridging contacts, *N_B_*, and the inter-strand stacking energy, 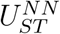, for (A) dT_50_ and (B) dA_50_. Color circles denote the spermine concentration. For both sequences, *N_B_* increases as 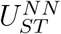 becomes more favorable. The red dashed line is a least-squares linear regression of *N_B_* versus 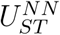, included. Pearson correlation coefficient (*r*) and Spearman rank correlation coefficient (*ρ*) are shown in each panel. (C) Ratio of 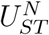 and 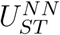 per dT_50_ chain as a function of Spm^4+^ concentration. Green and blue stars indicate the onset of phase separation, and reentrance, respectively. (D) Same as (C), except the results are for dA_50_. (E) Schematic of a dT_50_ condensate (purple) in 10 mM Spm^4+^ (cyan). A segment of a dT_50_ chain within the condensate is highlighted to illustrate weak base stacking. A representative spermine-mediated network between dT_50_ chains is shown. (F) Same as (E), except the results for dA_50_. Although both are fluid-like, the network density is greater in dA_50_ than in dT_50_.

The condensate phase is stabilized by the cumulative effect of many weak inter-chain contacts. Figure S13 shows the distributions of the cumulative inter-strand stacking energy, 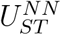, experienced by individual chains in the three phases. Because 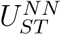 arises from the sum of many weak inter-chain stacking contacts, it is better viewed as a collective, mean-field-like manifestation of base hydrophobicity rather than as a set of discrete pairwise interactions. Even in the condensed phase, where intermolecular contacts are most abundant, the average cumulative inter-chain stacking energy per chain remains weaker than the stabilization provided by a single ideal intra-chain stack. Nevertheless, the simultaneous formation of many such weak contacts across multiple chains drive separation without altering the conformational ensemble of the individual polymers.

## Conclusion and Discussion

Let us summarize the major findings from the joint simulation and experimental study of spermine-driven phase transitions in homopolymeric single stranded DNA sequences. (1) We discovered that dT_50_ and dA_50_ undergo two transitions as a function of Spm^4+^ concentration. The first, from the dispersed to the condensate phase at low spermine concentration, is *sequence independent*. However, the stability of the condensate phase *depends on the sequence*. Because the droplet stability is principally determined by stacking interactions, the dA_50_ condensate is more stable over a broader spermine concentration range than the dT_50_ droplets. (2) At a higher, *sequence dependent* Spm^4+^, the sequences undergo a reentrant transition, at which the condensate phase becomes unstable due to near complete loss of Spm^4+^-mediated inter chain interactions. (3) The two sequences spontaneously form spherical condensates at long times by different mechanisms. Upon formation of small clusters, dT_50_ grows by the addition of one monomer at a time until the spherical droplet forms. In contrast, dA_50_ forms small droplets, which fuse leading to larger droplets - a coalescence-driven mechanism distinct from sequential chain addition. (4) The DNA chains in the condensates are held together by only a few bridging spermine molecules per chain. The number of spermine bound per chain increases modestly with spermine concentration in the two phase region and in the reentrant phase. However, the small number of spermine molecules that bridge two or more chains decreases precipitously as reentrance approaches. Despite this substantial change in intermolecular connectivity, the conformation of the ssDNA chains in the dispersed, condensed, and reentrant phases are almost identical.

### Condensed phases form liquid-like network without chain collapse

Besides the stacking and electrostatic interactions, it is possible that charge fluctuations [20, 51–54] could lead to an effective inter-chain attraction, thus contributing to the observed phase changes. However, in ssDNA the magnitude of attraction is insufficient to drive phase separation because in the absence of inter-chain stacking, phase separation does not occur (Figure 3). Remarkably, the formation of the droplet phase and subsequent dissolution requires the weak spermine dependent non-neighboring stacking interactions, which explains (Figure S17) the dynamic fluid-like network in the dense phase without chain collapse.

### On the invariance of the ssDNA conformations in the three phases

It is surprising that the chain conformations in the three phases are virtually identical, even though there are visible changes in the ssDNA-spermine interactions during the phase transitions. This finding may be understood by noting that the condensate stability is principally determined by the number of spermine molecules that bridge two or more ssDNA chains (see Figures 4B and C). The maximum value of *N_B_* per dT_50_ (dA_50_) is only 1.5 (2.5). On the other hand, *N_D_*, the average number of Spm^4+^ molecules that decorate each ssDNA chain is ∼ (11–12). In the three phases, the ssDNA chains may be pictured as being almost isolated, which explains the preservation of the ssDNA conformations. These findings are in accord with theoretical predictions [55], which showed that flexible low-complexity monomers, such as dT_50_ and dA_50_ should form liquid-like condensates without significant conformational changes.

The results in Figure 6 imply that phase separation and single-chain structures are only partially decoupled. Thus, condensate formation and dissolution are governed primarily by changes in the network connectivity. The network stability is determined by sparse ion-mediated bridges, stabilized by many weak inter-chain stacking contacts (Figure 7), without involving large-scale restructuring of the individual polymers. Remarkably, the number of bridging interactions decreases linearly as the strength of the sequence dependent non-neighboring staking interactions decreases (Figure 7 (A - B)). The weakness of the non-neighboring interactions is evident in also evident in (Figure 7 (C - D)), which shows the ratio 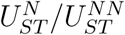 is far greater than unity. This mechanism should be contrasted with condensation of dsDNA that is accompanied by pronounced chain compaction. Homopolymeric ssDNA condensates form by a fluid-like “*network-without-chain collapse*”.

### Condensate and reentrant transition mechanisms

Condensation at a critical concentration of Spm^4+^ followed by reentrant transition at a higher spermine concentration in long double stranded DNA (dsDNA) was predicted in a prescient theoretical study [56]. The first transition is a consequence of substantial decrease in the negative charge on dsDNA (nearly vanishing in long dsDNA) with increasing Spm^4+^, as predicted by the Oosawa-Manning counterion condensation theory [42, 51, 57]. As a result, the dsDNA molecules experience a short range attraction resulting in chain condensation. As the Spm^4+^ concentration increases and exceeds a critical value, the net charge becomes *positive* by the charge inversion mechanism [20, 56]. Subsequently, the dsDNA undergoes reentrant transition, resulting in the dissolution of the chain. Thus, the occurrence of the two transitions could be explained based solely on the electrostatic interactions that are modulated by the Spm^4+^ [56].

The transition between the dispersed and the condensate phase in ssDNA is similar to condensation in dsDNA. In both cases, it is initiated when the net charge on DNA is reduced by Spm^4+^. The extent of charge neutralization in the short ssDNA (≈ −0.5*e* per phosphate group - see SI for details) is much less than in the long dsDNA where it nearly vanishes. The reentrant mechanism in ssDNA is driven by both the loss in the bridging interactions (Figures 4A and B) and the decrease in the weak, but important, non-neighboring stacking (Figure 7 and Figure S13). These weak interactions that render the ssDNA droplets stable are both sequence dependent. Clearly, this mechanism is distinct from a purely electrostatic charge-inversion model of reentrance.

## Methods

### Coarse-grained (CG) models for DNA-Polyamines

The total energy of a system consisting of DNA and polyamines is,

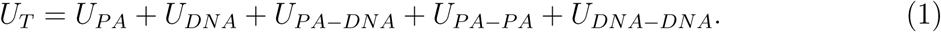

The first two terms in Eq. 1 represent the energy functions for PA and DNA, respectively. The last three terms account for interactions between PA and DNA, PA-PA, and DNA chains, respectively.

### Spermine

We modeled spermine using seven coarse-grained interaction sites. Four represent the positively charged amine groups and the others correspond to the alkyl groups (Figure 1A). The energy of a PA molecule is, *U_PA_* = *U_B_* + *U_EV_* + *U_El_*. The bond stretch term, *U_B_*), is given by a harmonic potential, *U_B_* = *k_b_*(*r* − *r*_0_)^2^. The equilibrium bond length, *r*_0_, is the sum of the radii of the bonded beads *i* and *j*, and *k_b_* = 20.0 kcal/mol/Å^2^ is the spring constant. Excluded volume repulsion, *U_EV_*, between any two non-bonded sites *i* and *j*, separated by a distance, *r* (in nm), is,

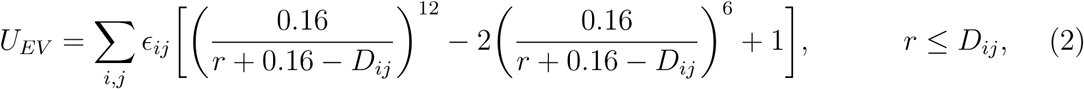

where *D_ij_* = *R_i_*+*R_j_* and 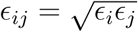; *R_i_* and *R_j_* are the radii of *i^th^* and *j^th^* beads, respectively, and *ɛ_i_* and *ɛ_j_* are the respective interaction strengths. The values of *R_i_* and *ɛ* for the amine beads are taken from the OPLS-AA [58] force field. We used the same *ɛ* values for the amine and the alkyl groups because the short-range nature of the repulsive excluded volume interaction should be insensitive to the *ɛ* values. The radii for the alkyl groups, (CH_2_)_3_ and (CH_2_)_4_, are taken from the Self-Organized Polymer model with side chains (SOP-SC) [59, 60], corresponding to the side chain radii of valine and leucine, respectively.

### Electrostatic interactions

We model the electrostatic interactions between the amine groups with charges *Q_i_* and *Q_j_* that are separated by a distance *r_ij_* using the Coulomb potential,

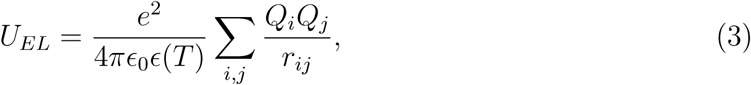

where *ɛ*_0_ is the vacuum permittivity. The temperature-dependent dielectric constant is taken to be[61],

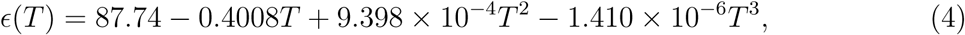

where the temperature, *T*, is in Celsius. The charge on the amine group is *Q_N_* = 1e (unit of electron charge). The alkyl groups are neutral (*Q_C_* = 0e). The PAs interact with other PAs, DNA and the ions, through the excluded volume and electrostatic potentials.

### Three Site Interaction with ions (TIS-ION) model for DNA

The general energy function in the coarse-grained TIS-ION model for DNA [32] is *U_T IS−ION_* = *U_B_* + *U_Ang_* + *U_EV_* + *U_EL_* + *U_HB_* + *U_ST_* . The potentials for bond stretch (*U_B_*) and bond angle (*U_Ang_*) are, *U_B_* = *k_b_*(*r*−*r*_0_)^2^ and *U_Ang_* = *k_θ_*(*θ* −*θ*_0_)^2^, where *r*_0_ and *θ*_0_ are the equilibrium bond length and bond angle, respectively, taken from an ideal B-form DNA helix. Excluded volume repulsion between sites *i* and *j* (DNA sites or ions) has the same form as eq. 2. The interaction between the phosphate beads is given by the Coulomb potential. The charge on the phosphate group is *Q_P_* = −1e. Because the ssDNA investigated here are homopolymers, *U_HB_* is not relevant. *U_ST_* : Stacking interactions between bases are written as 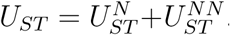. The contribution arising from adjacent bases 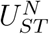 is,

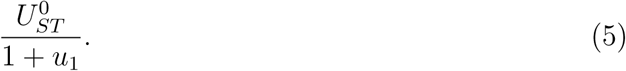

The potential *u*_1_ given by,

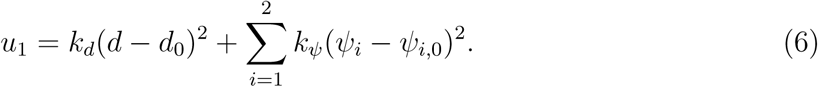

In the equation above, *d* is the distance between the bases B_i_ and B_i+1_, and *ψ*_1_ (P_i_, S_i_, P_i+1_, S_i+1_) and *ψ*_2_ (P_i+2_, S_i+1_, P_i+1_, S_i_) are the dihedral angles around the sugar-phosphate bonds (Figure S9). The equilibrium stacking distance, *d*_0_, and the backbone dihedral angles *ψ*_1,0_ and *ψ*_2,0_ as well as 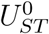, the strength of the stacking energy between two adjacent nucleotides are calculated from the coarse-grained representation of an ideal B-form DNA helix.

Non-neighboring bases within a single chain or between multiple chains could engage in base stacking interactions. The contribution to *U_ST_* due to stacking between non-neighboring (NN) bases that are separated by at least one base (Figure S9), 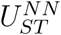, is,

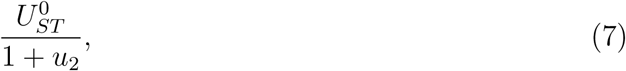

where

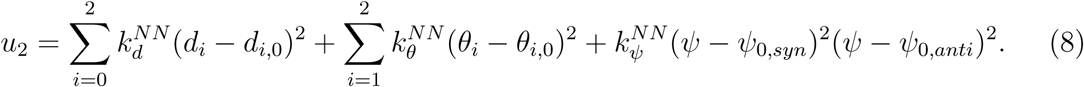

The dihedral angle, *ψ*(S_i_, B_i_, B_j_, S_j_), around any two non-neighboring bases *B_i_* and *B_j_* specifies the glycosidic orientation of the stacked bases. In NN stacks, both syn (*ψ*_0*,syn*_) and anti (*ψ*_0*,anti*_) orientations of the glycosidic bonds are allowed as found in RNA structures [62] and are also observed in the structures of the ssDNA-protein complexes [63]. To account for both these possibilities, we used a double well potential for the dihedral angle (Eq. 8 and Figure S9).

We calculated *ψ*_0*,anti*_ from the Protein Data Bank (PDB) structure of a 16 nucleotide ssDNA in complex with the Human Transcription Cofactor PC4 containing two non-neighboring bases (dT) that form stacks (PDB ID: 2C62). The rest of the parameters, such as, *d*_0_, *d*_1_, *d*_2_, *θ*_1,0_, *θ*_2,0_, *ψ*_0*,syn*_ are calculated from the adjacent stacks of ideal-B DNA helix (Figure S9). The values of 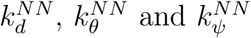 are optimized to reproduce the structural properties of the homopolymeric ssDNA chains dA_30_ and dT_30_ [32] (see Figure S10). The force-field parameters are listed in Tables S2-4.

Comparison of the distributions of the radius of gyration *R_g_*, *P* (*R_g_*), and the effective Flory exponents *ν*, for the monomeric sequences dT_50_ and dA_50_ in 10 mM Spm^4+^, at *T* = 298 K, with and without 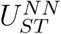, are essentially the same (Figures S11). Additional comparison with experiments are provided in the SI (Figure S10), for dT_30_ and dA_30_ chains, in solution of Mg^2+^ and NaCl [33]. Inclusion of the non-neighboring stacks, which is inconsequential in short monomers, is important in stabilizing the condensate phase.

### Simulations

We performed low friction Langevin dynamics simulations of monomeric dT_50_ and dA_50_ constructs in spermine (details are in the SI). To probe PA-driven DNA coacervation and reentrant behavior, we performed simulations by randomly placing multiple ssDNA chains in a cubic box and varied the PA concentration. In the simulations, a fixed DNA concentration of 350 *µ*M was used.

The multi chain simulations were unbiased in the sense that condensate formation occurs when the polymer chains coalesce spontaneously as the spermine concentration is increased. We measured concentration of monomers in the condensed phase (*ρ_c_*), as well as transfer free energy, Δ*G_tr_*, as a function of *N*, the number of chains in the simulation box. Both quantities show negligible variations when *N* ≥ 90. Because the observed finite-size effects are modest even at *N* = 64, we constructed phase diagram with *N* = 64 to maintain computational efficiency (Figure S7).

### Experiments

To validate the computational predictions, we performed experiments to monitor Spm^4+^-induced phase transitions. Lyophilized DNA, purchased from IDT, USA, was resuspended in IDTE pH 7.5 (IDT, USA) buffer at 250 *µ*M concentration and subsequently stored at -20 °C. Spermine-tetra-hydrochloride and green-fluorescent spermine (Cat # SCT250) were obtained from Millipore Sigma, USA. Stock solutions of spermine (100 mM in nuclease free water) and green-fluorescent spermine (5 mM in DMSO) were stored at – 20 °C and - 80 °C, respectively.

#### Coacervate formation

Complex coacervates were formed by mixing DNA and spermine at *T* = 295 K at the following solution conditions: DNA = (25 - 75) *µ*M, spermine = (1-20) mM in 10 mM Tris pH 7.0 (Thermo Fisher, USA). Freshly prepared samples were deposited onto a passivated [28] 384 glass bottom well plate (Brooks, Matri Plate MGB101-1-2-LG-L) and subsequently characterized using an Andor Spinning Disc setup (Oxford Instruments, USA).

#### Soluble DNA fraction

Soluble DNA fraction was measured using a Nanodrop 8000 spectrophotometer (Thermo Fisher, USA). Fresh DNA/spermine complex coacervates, prepared at room temperature, were transferred to 1.5 ml Eppendorf tubes. Samples were then centrifuged at 16,000g for 20 minutes. Next, few microliters of supernatant was collected and DNA concentration analyzed using the spectrophotometer at room temperature. See SI for additional details.

#### Fluorescence Recover After Photobleaching (FRAP)

To assess the fluid-like nature of the condensates, we measured the mean fluorescence intensity of the circular bleached region, *B*(*t*) (1 *µ*m diameter), and the mean intensity at the center of an unbleached coacervate, *U* (*t*) (1 *µ*m diameter), before and after photobleaching. The background intensity, *N* (*t*), was measured in a circular area of the surrounding solution phase outside the droplet. The corrected intensity, *I*(*t*), was calculated as,

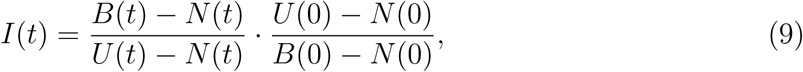

where *U* (0), *B*(0), and *N* (0) are the mean pre-bleach intensities. The recovery data were corrected for signal loss due to the bleaching pulse and for photobleaching during imaging. The normalized recovery, *I_N_* (*t*), was fit to,

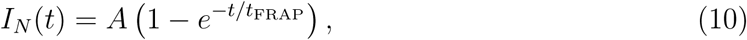

where *A* is the recovery amplitude and *t*_FRAP_ is the characteristic FRAP recovery time.

#### FRET

To measure FRET efficiency of Cy3-Cy5 labeled dT_50_ chains, we recorded fluorescence intensities in three channels: DD, DA, and AA. The DD channel corresponds to donor emission (600/50 nm filter) after excitation at 561 nm. The DA channel corresponds to acceptor emission (700/50 nm filter) after excitation at 561 nm. The AA channel corresponds to acceptor emission (700/75 nm filter) after acceptor excitation at 637 nm. For the Cy3-Cy5 pair, the Förster radius is *R*_0_ = 6 nm [40]. To account for cross-channel bleed-through, we determined the leakage coefficients *α* and *β* from singly labeled controls:

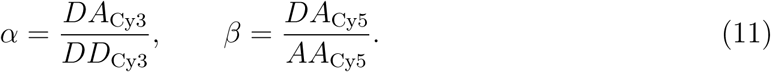

Using *α* and *β*, the corrected FRET signal was calculated as

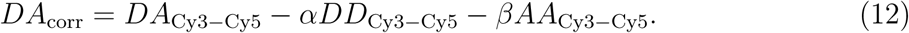

The corrected FRET efficiency was calculated using,

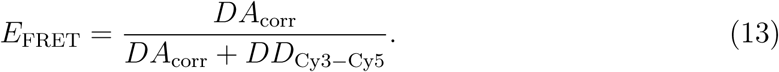

To probe dT_50_ conformation, we used two dual-labeled constructs. In term-dT_50_, Cy3 was attached to the 5’ end and Cy5 to the 3’ end. In int-dT_50_, Cy3 was attached to the 5’ end and Cy5 was embedded between the 25th and 26th nucleotides from either end. For FRET measurements in the dispersed phase, we prepared a mixture containing 1 mM spermine, 47.5 *µ*M dT_50_, and 2.5 *µ*M term-dT_50_ or int-dT_50_ in Tris buffer at pH 7.0. Under these conditions, phase separation did not occur. For measurements in the reentrant phase, the DNA concentration was kept the same and the spermine concentration was increased to 20 mM.

## Author Contributions

B.M, A.J, and D.T. designed the research. B.M. performed the simulations. S.M. performed the experiments. B.M., S.M., A.J., and D.T. interpreted the results and wrote the manuscript. All authors reviewed and approved the final manuscript.

## Acknowledgments

This work was supported by the NSF (grants CHE 2320256 to D.T.), NIH (grant R35GM151111 to A.J.) and the Welch Foundation (grant F-0019 to D.T. administered through the Collie-Welch Regents Chair). We thank the Texas Advanced Computing Center and Pittsburgh Supercomputing Center for providing generous computational resources.

## Data Availability

Simulations were performed using an in-house code built on the OpenMM platform. The code is available at https://github.com/balaka92/TIS-DNA-Ex-Ion and videos are available at https://zenodo.org/records/22212158.

## References

[1] T Thomas and TJ Thomas. Polyamines in cell growth and cell death: molecular mechanisms and therapeutic applications. Cell. Mol. Life Sci., 58(2):244–258, 2001.

[2] Frank Madeo, Tobias Eisenberg, Federico Pietrocola, and Guido Kroemer. Spermidine in health and disease. Science, 359(6374):eaan2788, 2018.

[3] Leonard C Gosule and John A Schellman. Compact form of DNA induced by spermidine. Nature, 259(5541):333–335, 1976.

[4] Leonard C Gosule and John A Schellman. DNA condensation with polyamines: I. spectroscopic studies. Journal of Molecular Biology, 121(3):311–326, 1978.

[5] Dhruba K Chattoraj, Leonard C Gosule, and John A Schellman. Dna condensation with polyamines: II. Electron microscopic studies. Journal of molecular biology, 121(3):327–337, 1978.

[6] Phillip Wulfridge and Kavitha Sarma. Intertwining roles of R-loops and G-quadruplexes in DNA repair, transcription and genome organization. Nature cell biology, 26(7):1025–1036, 2024.

[7] Eda Koculi, Nam-Kyung Lee, D Thirumalai, and Sarah A Woodson. Folding of the Tetrahymena ribozyme by polyamines: Importance of counterion valence and size. J. Mol. Biol., 341(1):27–36, 2004.

[8] Sangwoo Park, Raquel Merino-Urteaga, Violetta Karwacki-Neisius, Gustavo Ezequiel Carrizo, Advait Athreya, Alberto Marin-Gonzalez, Nils A Benning, Jonghan Park, Michelle M Mitchener, Natarajan V Bhanu, et al. Native nucleosomes intrinsically encode genome organization principles. Nature, 643(8071):572–581, 2025.

[9] Victor A. Bloomfield. DNA condensation by multivalent cations. Biopolymers, 44(3):269–282, 1997.

[10] Richard W. Wilson, Victor A. Bloomfield, and Donald C. Rau. Polyelectrolyte effects in DNA condensation by polyamines. Biophys. Chem, 11(1):51–62, 1980.

[11] Eric Raspaud, Dominique Durand, and Françoise Livolant. Interhelical spacing in liquid crystalline Spermine and Spermidine-DNA precipitates. Biophys. J., 88(1):392–403, 2005.

[12] Eric Raspaud, M Olvera De La Cruz, J-L Sikorav, and Francoise Livolant. Precipitation of DNA by polyamines: a polyelectrolyte behavior. Biophys. J., 74(1):381–393, 1998.

[13] Victor A. Bloomfield. Condensation of dna by multivalent cations: Considerations on mechanism. Biopolymers, 31(13):1471–1481, 1991.

[14] J. Pelta, F. Livolant, and J.-L. Sikorav. DNA aggregation induced by polyamines and cobalthexamine. J. Biol. Chem., 271(10):5656–5662, 1996.

[15] M. Saminathan, T. Antony, A. Shirahata, L. H. Sigal, T. Thomas, and T. J. Thomas. Polyamine structural effects on the induction and stabilization of liquid crystalline DNA: Potential applications to DNA packaging, gene therapy and polyamine therapeutics. Nucleic Acids Res., 30(17):3722–3731, 2002.

[16] Bart van den Broek, Marc C. Noom, and Gijs J. L. Wuite. Visualizing the formation and collapse of DNA toroids. Biophys. J., 98(9):1902–1910, 2010.

[17] IM Lifshitz, A Yu Grosberg, and AR Khokhlov. Some problems of the statistical physics of polymer chains with volume interaction. Reviews of Modern Physics, 50(3):683, 1978.

[18] Toan T. Nguyen, Ioulia Rouzina, and Boris I. Shklovskii. Reentrant condensation of DNA induced by multivalent counterions. J. Chem. Phys., 112(5):2562–2568, 2000.

[19] Viet Duc Nguyen, Toan T. Nguyen, and Paolo Carloni. DNA like-charge attraction and overcharging by divalent counterions in the presence of divalent co-ions. J. Biol. Phys., 43:257– 272, 2017.

[20] A Yu Grosberg, TT Nguyen, and BI Shklovskii. Colloquium: The physics of charge inversion in chemical and biological systems. Rev. Mod. Phys., 74(2):329, 2002.

[21] Hung T Nguyen, Naoto Hori, and D Thirumalai. Condensates in RNA repeat sequences are heterogeneously organized and exhibit reptation dynamics. Nature Chem., 14(7):775–785, 2022.

[22] Ofer Kimchi, Ella M King, and Michael P Brenner. Uncovering the mechanism for aggregation in repeat expanded RNA reveals a reentrant transition. Nat. Commun., 14(1):332, 2023.

[23] Yiqing Shen, Serguei V Kuznetsov, and Anjum Ansari. Loop dependence of the dynamics of DNA hairpins. J. Phys. Chem. B, 105(48):12202–12211, 2001.

[24] MC Murphy, Ivan Rasnik, Wei Cheng, Timothy M Lohman, and Taekjip Ha. Probing single-stranded DNA conformational flexibility using fluorescence spectroscopy. Biophys. J., 86(4):2530–2537, 2004.

[25] Bernard Tinland, Alain Pluen, Jean Sturm, and Gilbert Weill. Persistence length of single-stranded DNA. Macromolecules, 30(19):5763–5765, 1997.

[26] Anisha Shakya and John T King. DNA local-flexibility-dependent assembly of phase-separated liquid droplets. Biophys. J., 115(10):1840–1847, 2018.

[27] Alain AM André and Evan Spruijt. Rigidity rules in DNA droplets: Nucleic acid flexibility affects model membraneless organelles. Biophys. J., 115(10):1837–1839, 2018.

[28] Sumit Majumder, Sebastian Coupe, Nikta Fakhri, and Ankur Jain. Sequence-encoded intermolecular base pairing modulates fluidity in dna and rna condensates. Nat. Commun., 16(1):1–12, 2025.

[29] William M Aumiller Jr, Fatma Pir Cakmak, Bradley W Davis, and Christine D Keating. RNA-based coacervates as a model for membraneless organelles: formation, properties, and interfacial liposome assembly. Langmuir, 32(39):10042–10053, 2016.

[30] Paul Pullara, Ibraheem Alshareedah, and Priya R Banerjee. Temperature-dependent reentrant phase transition of RNA–polycation mixtures. Soft matter, 18(7):1342–1349, 2022.

[31] Ankur Jain and Ronald D Vale. RNA phase transitions in repeat expansion disorders. Nature, 546(7657):243–247, 2017.

[32] Balaka Mondal, Debayan Chakraborty, Naoto Hori, Hung T Nguyen, and Dave Thirumalai. Competition between stacking and divalent cation-mediated electrostatic interactions determines the conformations of short DNA sequences. J. Chem. Theory Comput., 20(7):2934–2946, 2024.

[33] Alex Plumridge, Steve P Meisburger, Kurt Andresen, and Lois Pollack. The impact of base stacking on the conformations and electrostatics of single-stranded dna. Nucleic Acids Res., 45(7):3932–3943, 2017.

[34] Dustin B. McIntosh, Gina Duggan, Quentin Gouil, and Omar A. Saleh. Sequence-dependent elasticity and electrostatics of single-stranded DNA: Signatures of base-stacking. Biophys. J., 106:659–666, 2014.

[35] Alex Plumridge, Steve P Meisburger, and Lois Pollack. Visualizing single-stranded nucleic acids in solution. Nucleic Acids Res., 45(9):e66–e66, 2017.

[36] Xavier Viader-Godoy, Maria Manosas, and Felix Ritort. Stacking correlation length in single-stranded DNA. Nucleic Acids Res., 52(21):13243–13254, 2024.

[37] Debayan Chakraborty, Balaka Mondal, and D Thirumalai. Brewing COFFEE: a sequence-specific coarse-grained energy function for simulations of DNA-protein complexes. J. Chem. Theory Comput., 20(3):1398–1413, 2024.

[38] Debayan Chakraborty, Naoto Hori, and Dave Thirumalai. Sequence-dependent three interaction site model for single- and double-stranded DNA. J. Chem. Theory Comput., 14(7):3763– 3779, 2018.

[39] Mahmoud Moradi, Volodymyr Babin, Christopher Roland, Thomas A Darden, and Celeste Sagui. Conformations and free energy landscapes of polyproline peptides. Proc. Natl. Acad. Sci. USA, 106(49):20746–20751, 2009.

[40] Asif Iqbal, Sinan Arslan, Burak Okumus, Timothy J Wilson, Gerard Giraud, David G Norman, Taekjip Ha, and David MJ Lilley. Orientation dependence in fluorescent energy transfer between Cy3 and Cy5 terminally attached to double-stranded nucleic acids. Proc. Natl. Acad. Sci. USA, 105(32):11176–11181, 2008.

[41] Herbert B Callen. Thermodynamics and an introduction to thermostatistics. John Wiley& Sons, 2, 1993.

[42] Fumio Oosawa. Polyelectrolytes. Marcel Dekker, Inc., 1971.

[43] Balaka Mondal and D Thirumalai. Persistence length of short homopolymeric single-stranded DNA sequences in polyvalent cations. *bioRxiv*, 2026. Preprint.

[44] Bae-Yeun Ha and D Thirumalai. Bending rigidity of stiff polyelectrolyte chains: A single chain and a bundle of multichains. Macromolecules, 36(25):9658–9666, 2003.

[45] Murugappan Muthukumar. Theory of counter-ion condensation on flexible polyelectrolytes: Adsorption mechanism. J. Chem. Phys., 120(19):9343–9350, 2004.

[46] Naoto Hori, Natalia A Denesyuk, and D Thirumalai. Ion condensation onto ribozyme is site specific and fold dependent. Biophysical Journal, 116(12):2400–2410, 2019.

[47] Ilya M Lifshitz and Vitaly V Slyozov. The kinetics of precipitation from supersaturated solid solutions. J. Phys. Chem. Solids, 19(1-2):35–50, 1961.

[48] Carl Wagner. Theory of precipitate change by redissolution. Z. Elektrochem, 65:581–591, 1961.

[49] Peter W Voorhees. The theory of Ostwald ripening. J. Stat. Phys., 38(1):231–252, 1985.

[50] Farkhad Maksudov, Mauro L Mugnai, Laura Dominguez, Dmitrii E Makarov, and D Thirumalai. Droplet growth, Ostwald’s rule, and emergence of order in Fused in Sarcoma. Proc. Natl. Acad. Sci. USA, 123(4):e2519135123, 2026.

[51] Fumio Oosawa. Interaction between parallel rodlike macroions. Biopolymers: Original Research on Biomolecules, 6(11):1633–1647, 1968.

[52] Amy E Larsen and David G Grier. Like-charge attractions in metastable colloidal crystallites. Nature, 385(6613):230–233, 1997.

[53] Niels Grønbech-Jensen, Robert J Mashl, Robijn F Bruinsma, and William M Gelbart. Counterion-induced attraction between rigid polyelectrolytes. Phys. Rev. Lett., 78(12):2477, 1997.

[54] William M Gelbart, Robijn F Bruinsma, Philip A Pincus, and V Adrian Parsegian. Dna-inspired electrostatics. Physics today, 53(9):38–45, 2000.

[55] Ryota Takaki and Dave Thirumalai. Sequence complexity and monomer rigidity control the morphologies and aging dynamics of protein aggregates. Proc. Natl. Acad. Sci. USA, 121(50):e2409973121, 2024.

[56] Toan T Nguyen, I Rouzina, and Boris I Shklovskii. Reentrant condensation of DNA induced by multivalent counterions. J. Chem. Phys., 112(5):2562–2568, 2000.

[57] G. S. Manning. Limiting Laws and Counterion Condensation in Polyelectrolyte Solutions I. Colligative Properties. J. Chem. Phys., 51:924–933, 1969.

[58] Robert C Rizzo and William L Jorgensen. OPLS all-atom model for amines: Resolution of the amine hydration problem. J. Am. Chem. Soc., 121(20):4827–4836, 1999.

[59] Zhenxing Liu, Govardhan Reddy, Edward P OâBrien, and D Thirumalai. Collapse kinetics and chevron plots from simulations of denaturant-dependent folding of globular proteins. Proc. Natl. Acad. Sci. USA, 108(19):7787–7792, 2011.

[60] Changbong Hyeon, Ruxandra I Dima, and D Thirumalai. Pathways and kinetic barriers in mechanical unfolding and refolding of RNA and proteins. Structure, 14(11):1633–1645, 2006.

[61] J. B. Hasted. Water: A comprehensive treatise. The Physics and Physical Chemistry of Water, 1:255–305, 1972.

[62] Ayush Jhunjhunwala, Zakir Ali, Sohini Bhattacharya, Antarip Halder, Abhijit Mitra, and Purshotam Sharma. On the nature of nucleobase stacking in RNA: a comprehensive survey of its structural variability and a systematic classification of associated interactions. J. Chem. Inf. Model., 61(3):1470–1480, 2021.

[63] Raktim Mitra, Ari S Cohen, Jared M Sagendorf, Helen M Berman, and Remo Rohs. Dnaprodb: an updated database for the automated and interactive analysis of protein–DNA complexes. Nucleic Acids Res., 53(D1):D396–D402, 2025.

[64] Aric A. Hagberg, Daniel A. Schult, and Pieter J. Swart. Exploring network structure, dynamics, and function using NetworkX. In Gaël Varoquaux, Travis Vaught, and Jarrod Millman, editors, Proceedings of the 7th Python in Science Conference, pages 11–15, 2008.

