## Supplementary material for "Interchain base-stacking and spermine bridges govern ssDNA condensation and reentrant transitions": SI

### Computational Details:

**Non-neighboring (NN) stacking interactions:** If the length of the ssDNA is long, it is conceivable that the chain could fold onto itself, thus making it necessary to include NN stacking interactions (see Figure 1B in the main text). The contribution due to NN stacking of bases that are separated by at least one nucleobase, $U_{ST}^{NN}$ is taken to be $\frac{U_{ST}^{0}}{1+u_2}$, where

$$u_2 = \sum_{i=0}^{2} k_d^{NN}(d_i - d_{i,0})^2 + \sum_{i=1}^{2} k_\theta^{NN}(\theta_i - \theta_{i,0})^2 + k_\psi^{NN}(\psi - \psi_{0,syn})^2(\psi - \psi_{0,anti})^2. \qquad \text{(S1)}$$

In Eq. S1, $\psi(\mathrm{S_i}, \mathrm{B_i}, \mathrm{B_j}, \mathrm{S_j})$ is the dihedral angle (Figures S9 (A) and (C)) around any two non-neighboring bases $B_i$ and $B_j$, specifying the glycosidic orientation of the stacked bases. In NN stacks, both syn ($\psi_{0,syn}$) and anti ($\psi_{0,anti}$) (Figure S9) orientations of the glycosidic bonds are allowed. Their relevance has been demonstrated in RNA [1] and in ssDNA-protein complexes.[2] To account for $\psi_{0,syn}$ and $\psi_{0,anti}$, we used a double well potential for the dihedral angle (Eq. S1 and Figure S9). The strength of the stacking interactions between two bases, $U_{ST}^{0}$ (Eq. 7 in main text), is the same for both adjacent and non-neighboring pairs, because the underlying chemical nature of stacking is unchanged regardless of their relative positions (see Table S4).

We obtained the value of $\psi_{0,anti}$ from the Protein Data Bank (PDB) structure of a 16 nucleotide ssDNA in complex with the Human Transcription Cofactor PC4 containing two non-neighboring bases (dT) form stacks (PDB ID: 2C62). An estimate of $\psi_{0,syn}$ was made from adjacent stacks in the ideal B-form DNA helix, corresponding to the *syn* glycosidic conformation. Other parameters, such as $d_0$, $d_1$, $d_2$, $\theta_{1,0}$, and $\theta_{2,0}$, were also calculated from adjacent stacks of the ideal B-DNA helix (see Figure S9B and Table S3). To determine the spring constants ($k_d^{NN}$, $k_\theta^{NN}$, $k_\psi^{NN}$ in Eq. S1), we performed simulations in $Mg^{2+}$ by including NN stacking interactions. The adjacent stacking parameters were kept fixed (see Eqs. 5 and 6 in the main text). The parameters $k_d^{NN} = 5\,\text{Å}^{-2}$, $k_\theta^{NN} = 5\,\text{radians}^{-2}$, and $k_\psi^{NN} = 5\,\text{radians}^{-2}$ reproduce the structural properties, such as the radius of gyration ($R_g$)

and the effective Flory exponent ($\nu$), of the homopolymeric ssDNA chains $dA_{30}$ and $dT_{30}$ in the divalent cation, $Mg^{2+}$.[3] The agreement of $R_g$ with and without NN interactions show that non-neighboring stacking has only a minimal impact on single-chain properties (see Figure S10). However, the weak NN stacking plays an indispensable role in stabilizing the $Spm^{4+}$ -induced condensate phase (see the main text).

**Simulations:** We used OpenMM,[4] a high-performance toolkit for molecular simulations, to simulate single and multiple ssDNA homopolymers. The simulations were performed on a Graphics Processing Unit (GPU). To account for the long-range nature of the Coulomb interactions we used Particle-Mesh-Ewald (PME) algorithm.[5] We integrated the Langevin equation in the low friction limit[6] to enhance conformational sampling. The equation of motion for a site with coordinate $\vec{r_i}$, is,

$$m_i\ddot{\vec{r_i}} = -m_i\zeta\dot{\vec{r_i}} + \vec{F_c} + \vec{\Gamma}, \tag{S2}$$

where $m_i$ is the mass of the bead, $\zeta$ is the friction coefficient, $\vec{F_c} = -\frac{\partial U_{TIS-ION}(\{\mathbf{r}\})}{\partial \vec{r_i}}$ is the systematic force, and $\vec{\Gamma}$ is the random force with a white noise spectrum. The autocorrelation function of the random force in the discretized form is $\langle \Gamma(t)\ \Gamma(t+nh)\rangle = \frac{2\zeta m_i k_B T}{h}\delta_{0,n}$, where $n = 0, 1, ...$ and $\delta_{0,n}$ is the Kronecker delta function. The equations of motion were integrated using the Langevin leap-frog method[7] using $\zeta = 0.1$ ps$^{-1}$. We used an integration time step of 2 femtoseconds (fs) to advance the simulations. Time is measured in the unit of $\tau$. In real time it is given by, $\tau$ = (time step) $\times$ (data output frequency) = 2 fs $\times$ 5000 = 10 ps. All the simulations were carried out at $T = 298$ K.

**Classification of chains based on clustering:** We monitored phase separation by classifying each DNA chain as a monomer, a oligomer, or a condensate, based on a clustering algorithm (Figure S4). The algorithm is initiated by assigning each chain a unique cluster ID. Clusters were then iteratively merged based on 1.5 nm cutoff: if any coarse-grained beads from two chains were within this distance, they were assigned to the same cluster.

Furthermore, if a third chain came within 1.5 nm of any chain already in the cluster, it was also incorporated. The iterative merging process was continued until all the chains in the system were assigned to non-overlapping clusters. The distance cut-off of 1.5 nm is the average end-to-end distance of a $Spm^{4+}$ molecule, which can form bridging interactions across distinct DNA chains.

**Radius of gyration, $R_g$, and end-to-end distance, $R_{ee}$:** The radius of gyration, $R_g$ is calculated using,

$$R_g = \sqrt{\frac{1}{2{N_S}^2} \sum_{i=1}^{N_S} \sum_{j=1}^{N_S} r_{ij}^2}, \tag{S3}$$

where $N_S$ is the total number of coarse-grained sites in the ssDNA chain, and $r_{ij}$ is the distance between beads $i$ and $j$. The end-to-end distance $R_{ee}$ is the distance between the terminal beads (Figure S4A).

**Persistence length, $l_p$:** We estimated $l_p$ of ssDNA chains by fitting the calculated end-to-end distance ($r_{ee}$) distribution function, $P(r_{ee}, L)$, to an analytical expression for a worm-like chain.[8,9] The analytic expression for $P(r_{ee}, L)$ is,

$$P(r_{ee}, L) = \frac{4\pi A r_{ee}^2}{(1 - r_{ee}^2)^{\frac{9}{2}}} \exp\left(\frac{-3t}{4(1 - r_{ee}^2)}\right), \tag{S4}$$

where $A$, the normalization constant is $A = \frac{4(\frac{3t}{4})^{\frac{3}{2}} \exp(\frac{3t}{4})}{\pi^{\frac{3}{2}}(4 + \frac{12}{(3t/4)} + \frac{15}{(3t/4)^2})}$ and $t = \frac{L}{l_p}$.

**Flory exponent, $\nu$:** To obtain $\nu$, we fit the calculated mean distance $R_{ij}$ between the phosphate groups $i$ and $j$ using,

$$R_{ij} = R_0 |i - j|^{\nu}, \tag{S5}$$

where $R_0$ is the prefactor, and $\nu$ is the effective Flory exponent. The equation above can only be used for homopolymers. For heteropolymers the assumption that the ssDNA polymers are self-similar at all scales (two segments with identical $|i - j|$ values would have the same statistical properties) would be violated. Due to the finite value of $N_S$ the range of $|i - j|$ is limited which likely affects that the extracted values of $\nu$ and $R_0$ might not be accurate.

**Persistent stacks:** To describe the effects of stacking in ssDNA, we calculated $p_S$, which is related to the helix content.[10] We assume that each nucleotide is either in a stacked ($S$) or in a random coil state ($C$). To measure the stacking propensity of the $i^{th}$ nucleotide, we calculated the sugar-phosphate dihedral angles, $\psi(P_i, S_i, P_{i+1}, S_{i+1})$ and $\phi(S_i, P_{i+1}, S_{i+1}, P_{i+2})$, and compared them to $\psi_B$ and $\phi_B$ values in an ideal B-form helix. If the deviations $\delta\psi(= |\psi - \psi_B|)$ and $\delta\phi(= |\phi - \phi_B|)$ are within 25% of the ideal helix parameters, we consider the nucleotide to be in the $S$ state. For a given ssDNA conformation, $p_S$ is defined as,

$$p_S = \frac{\sum_n^{n_S} N_{st}\Theta(N_{st} - 3)}{N_S^I}, \tag{S6}$$

where $n_S$ is the number of consecutive stacks along the ssDNA contour, $N_{st}$ is the length of the $n^{th}$ stack and $N_S^I$ is the length of the persistent stacks in completely stacked ideal B helix, and $\Theta(N_n - 3)$ is unity only if $N_n \geq 3$. A minimum of three consecutive nucleotides ($N_{st} \geq 3$) should be in the $S$ state to obtain a meaningful value for $p_S$. For instance, in the oligomer $CCSCC\overline{SSSS}CCC\overline{SSS}CCSSC$, there are two persistent stacks ($n_S = 2$) of length 4 and 3, ($N_1 = 4$, $N_2 = 3$) respectively, yielding $p_S = \frac{4+3}{20}$, which means that there are 7 nucleotides that are persistently stacked. In the example, the lengths of the longest stretch of stacked and unstacked bases are, $N_S^{max} = 4$ and $N_U^{max} = 3$, respectively.

**Helicity, $H$:** We computed ($H$) the helical content of the [11] by considering a set of four consecutive backbone points, denoted as $A, B, C, D$, which define the vectors $\vec{AB}$ and $\vec{CD}$. The midpoints of the vectors, $E$ and $F$, determine the vector $\vec{EF}$. The handedness of each segment is calculated as the normalized scalar triple product,

$$H_{ABCD} = \frac{\vec{EF} \cdot (\vec{CD} \times \vec{AB})}{|\vec{EF}|\,|\vec{CD}|\,|\vec{AB}|}. \tag{S7}$$

Segments exhibiting positive (negative) values of $H_{ABCD}$ correspond to right-handed (left-handed) helix. The overall helicity $H$ of the ssDNA strand is computed by summing these contributions along the backbone:

$$H = \sum_{i=1}^{n-3} H_{i,i+1,i+2,i+3}, \tag{S8}$$

where $n$ is the total number of nucleotides. For normalization, the calculated value of $H$ is reported relative to the total helicity of an ideal B-DNA helix of the same length.

**Angular correlation function,** $C(n)$**:** The orientational order along the contour of the chain is defined as,

$$C(n) = \frac{1}{N_n} \sum_{\substack{i,j \\ |j-i|=n}} \langle \cos(\theta_{i,j}) \rangle, \tag{S9}$$

where $n = |j-i|$ is the contour separation, $N_n$ is the number of pairs with $|j-i| = n$ and $\theta_{i,j}$ is the angle between $i^{th}$ and $j^{th}$ bond vectors. The symbol $\langle\rangle$ an average over the is ensemble of conformations sampled in the simulations. In calculating $C(n)$, we used sugar-phosphate bond vectors along the backbone.

**Shape parameters:** The gyration tensor $\mathbf{S}$ for a given conformation is defined as the $3 \times 3$ matrix,

$$\mathbf{S} = \begin{pmatrix} S_{xx} & S_{xy} & S_{xz} \\ S_{yx} & S_{yy} & S_{yz} \\ S_{zx} & S_{zy} & S_{zz} \end{pmatrix}, \tag{S10}$$

where each component $S_{ij}$ is computed using,

$$S_{ij} = \frac{1}{N_s} \sum_{k=1}^{N_s} (r_{ki} - \overline{r}_i)(r_{kj} - \overline{r}_j). \tag{S11}$$

In the above equation, $N_s$ is the total number of particles, $r_{ki}$ and $r_{kj}$ are the $i^{\text{th}}$ and $j^{\text{th}}$ Cartesian coordinates of particle $k$, respectively, and $\overline{r}_i$, $\overline{r}_j$ are the coordinates of the center of mass along the respective axes. Diagonalization of $\mathbf{S}$ yields the eigenvalues $\lambda_1 \leq \lambda_2 \leq \lambda_3$, quantifying the spatial extent of the conformation along the principal axes.

The radius of gyration $R_g$, characterizing the overall size of the structure, is given by

$$R_g = \sqrt{\lambda_1 + \lambda_2 + \lambda_3}. \tag{S12}$$

Using the gyration tensors, we calculated the following shape parameters.[12,13] The asphericity $b$, measuring the deviation from spherical symmetry, is,

$$b = \frac{\lambda_3 - \frac{1}{2}(\lambda_2 + \lambda_1)}{{R_g}^2}; \tag{S13}$$

The relative shape anisotropy $\kappa^2$, that is bounded between 0 (spherical) and 1 (rod-like). is a measure of the reflecting the overall anisotropy. It is given by,

$$\kappa^2 = 1 - \frac{3(\lambda_1\lambda_2 + \lambda_2\lambda_3 + \lambda_3\lambda_1)}{(\lambda_1 + \lambda_2 + \lambda_3)^2}. \tag{S14}$$

The shape of the condensates or individual monomers can be determined using $S$, where

$$S = 27\frac{\prod_{i=1}^{3}(\lambda_i - \bar{\lambda})}{R_g^6}. \tag{S15}$$

$S$ satisfies the bound $-0.25 \leq S \leq 1$. Negative values of $S$ correspond to oblate ellipsoids and positive values indicate prolate ellipsoids.

**Calculation of concentrations in the two phases:** Each condensate is treated as an effective ellipsoid[14] whose principal radii are determined from the eigenvalues of its gyration tensor. The volume of the $i^{th}$ condensate is:

$$V_i = 4\pi\sqrt{3}\lambda_1\lambda_2\lambda_3. \tag{S16}$$

Its concentration is $\rho_c^i = N_i/V_i$, with $N_i$ the number of chains in the droplet. The average condensate concentration is,

$$\langle\rho_c\rangle = \frac{1}{N_{\text{drop}}}\sum_{i=1}^{N_{\text{drop}}}\rho_c^i. \tag{S17}$$

The dilute-phase concentration is

$$\rho_d = \frac{N_{\text{tot}} - \sum_{i=1}^{N_{\text{drop}}} N_i}{V_{\text{box}} - \sum_{i=1}^{N_{\text{drop}}} V_i}, \tag{S18}$$

where $N_{\text{tot}}$ is the total number of chains, $N_{\text{drop}}$ is the total number of condensates, and $V_{\text{box}}$ is the simulation box volume.

**Effective phosphate charge, $q_i^{eff}$:** For reasons explained elsewhere[15–17] the computation of the renormalized charge on the phosphate groups in irregular shaped polymers, such as ssDNA, is non-trivial. We made an approximate estimate of the effective charge ($q_i^{eff}$) on each phosphate using the following prescription. Let the ion-specific cutoff for a phosphate–ion pair $(i, j)$ be,

$$r_c^{ij} = R_{\text{P}} + R_j + \Delta, \tag{S19}$$

where $R_{\text{P}}$ is the phosphate radius, $R_j$ is the radius of ion $j$ and $\Delta$ is the diameter of the ion. We define,

$$q_i^{eff} = q_i^{(0)} + \sum_{j \in \mathcal{N}(i)} q_j, \tag{S20}$$

where $q_i^{(0)} = -e$ is the bare phosphate charge and $\mathcal{N}(i)$ the set of ions satisfying the cutoff criterion.

### Experimental details:

**Chemicals:** Unlabeled and labeled DNA were purchased from IDT, USA. Lyophilized DNA was resuspended in IDTE pH 7.5 (IDT, USA) buffer at 250 µM concentration and stored at -20 ° C. Spermine-tetra-hydrochloride and green-fluorescent spermine (BODIPY fluorophore Cat # SCT250) were obtained from Millipore Sigma, USA. Stock solutions of spermine (100 mM in nuclease free water) and green-fluorescent spermine (5 mM in DMSO) stored at -20 °C and -80 °C, respectively.

**Complex coacervates:** Complex coacervates were formed by mixing DNA and spermine at room temperature at the following solution condition: DNA = 25-150 $\mu$M, spermine = 1-20 mM in 10 mM Tris pH 7.0 (Thermo Fisher, USA). Freshly prepared samples were deposited onto a passivated 384 glass bottom well plate (Brooks, Matri Plate MGB101-1-2-LG-L) and subsequently characterized using an Andor Spinning disk microscope (Oxford Instruments, USA) as described previously.[18]

**Soluble DNA fraction:** We measured the Soluble DNA fraction using a Nanodrop 8000 spectrophotometer (Thermo Fisher, USA). Briefly, fresh DNA /spermine complex coacervates were prepared at room temperature and subsequently transferred to 1.5 ml Eppendorf tubes. Samples were then centrifuged at 16,000g for 20 min. Next, few microliters of supernatant were collected and DNA concentration analyzed using the spectrophotometer at room temperature.

**DNA density in complex coacervates:** We quantified DNA concentration inside $dT_{50}$/spermine complex coacervates using fluorescence calibration. First, we calibrated the fluorescence intensity of Cy3-$dT_{50}$ as a function of probe concentration in IDTE pH 7.5 buffer. Next, we generated $dT_{50}$/spermine complex coacervates in which a small fraction of the DNA was Cy3-labeled, maintaining an overall $dT_{50}$:Cy3-$dT_{50}$ ratio of 1:X, where X is varied between 1000 and 2500. We measured intensity of Cy3-$dT_{90}$ probes within the resulting complex coacervates under the same imaging conditions used for the calibration. Using the calibration curve, we estimated the concentration of Cy3-$dT_{50}$. By accounting for the labeling ratio, we obtained the total $dT_{50}$ concentration in the coacervate phase.

**Limitations:** Fluorescent probes are widely used[19,20] to estimate concentrations of biomolecules within condensates but such measurements have several notable caveats. Apparent concentrations can be skewed by altered probe partitioning between the dilute and dense phases.[21] Furthermore, the quantum yield of the dye can depend on ionic environment within the condensate[22,23] which may differ from the conditions used for the calibration (here, IDTE pH 7.5). In addition, the dye label itself may slightly perturb the phase behavior or DNA conformation. We did not correct for these effects here. Thus, our measurements likely have a systematic uncertainty in the estimate of the absolute DNA concentration. Nevertheless, they robustly establish that DNA concentrations within the coacervate phase reaches high millimolar levels.

### Additional Results

**NN-stacking does not affect monomer conformations:** The distributions of the radius of gyration, $P(R_g)$, normalized end-to-end distance, $P(R_{ee}/L)$, and the effective Flory exponents $\nu$, for both $dT_{50}$ and $dA_{50}$ with and without $U_{ST}^{NN}$ in 10 mM $Spm^{4+}$ are essentially the same (Figures S11). Using 0.62 nm as the distance between adjacent phosphate groups in ssDNA, the contour length becomes $L = 0.62 \times (N_m - 1)$ nm, where $N_m$ is the number of nucleotides in the homopolymer. With $L$ at hand the persistence length can be calculated using Eq. S4. The $l_p$ values (Eq. S4) for $dT_{50}$ (Figure S11C) and $dA_{50}$ (Figure S11C) are also unaffected by the inclusion on NN stacking. Thus, for the short ssDNA chains, NN stacking interactions are negligible.

**Strength of stacking determines Monomer properties:** Visual inspection of the conformations of $dT_{50}$ at low and high $Spm^{4+}$ show that $dT_{50}$ behaves like a random coil (Figures S1(A)) whereas $dA_{50}$ has helical features (Figures S1(B). The angular correlation $C(n)$ decays rapidly as $|i-j|$ increases in $dT_{50}$, which implies absence of orientational order along the chain contour. Due to strong base stacking interactions, $C(n)$ is periodic in $dA_{50}$, characteristic of helical conformations. The periodicity corresponds to 10 bases (Figure 1C in the main text), which is equivalent to one helical turn. Additional properties at various $Spm^{4+}$ concentrations in Figures S1 confirm that $dT_{50}$ is a random coil whereas $dA_{50}$ has persistent helical character. The average helical content [11] for $dT_{50}$ is $\langle H \rangle \approx 0.15$ whereas it is $\approx 0.7$ for $dA_{50}$ (Figure S1). Like other properties, $P(H)$ the distributions are insensitive to $Spm^{4+}$ concentration.

Figure S1g shows that the average length of the longest stacked (unstacked) in $dT_{50}$

($dA_{50}$) is $\langle N_S^{\max}\rangle \approx 3$ nucleotides (16 nucleotides). The helical segments in $dT_{50}$ are short and are interspersed with longer unstacked stretches. The distribution of stacked length in $dA_{50}$ peaks at $\langle N_S^{\max}\rangle \approx 8$ nucleotides, with a tail that extends up to 16 nt. The distribution of unstacked segments in $dA_{50}$ is narrower, with $\langle N_U^{\max}\rangle \approx 3$ nt.

The differences between the two sequences are due to the interplay between the unfavorable charge phosphate-phosphate electrostatic repulsion ($U_{EL}$) and the favorable neighboring stacking interactions ($U_{ST}$). Because $U_{EL}$ for both the sequences are independent of $Spm^{4+}$ , we expect that the monomer conformations are determined by $U_{ST}$. Indeed, $U_{EL}$ exceeds $U_{ST}$ by approximately $8k_BT$ per nucleotide (Figure S1(H)) in $dT_{50}$. The sum of $U_{EL}$ and $U_{ST}$ per nucleotide is greater than zero (blue symbols in Figure S1(I)). As a result, random coil conformations dominate in the ensemble. In $dA_{50}$, favorable stacking interactions outweigh the repulsive electrostatic interactions by $2k_BT$ ((Figure S1(H)) with $U_{EL} + U_{ST} \sim -2k_BT$ per nucleotide Figure S1(I)) resulting in helix formation at all $Spm^{4+}$ concentrations.

### Condensate formation

To investigate condensate formation, we performed simulations by placing $N$ DNA chains in a cubic box and varied the $Spm^{4+}$ concentration. Let us first describe a few technical details that are important in understanding the simulation results. Using the trajectories we performed a number of tests to ensure that the ssDNA-polyamine systems are at equilibrium.
**Chemical Potentials and phase equilibrium:** Equilibrium between two phases requires that the chemical potentials of every chemical species in the distinct phases should be equal. For coexistence between the dilute and condensed phases the equality, $\mu_i^{\mathrm{d}} = \mu_i^{\mathrm{c}}$ for each species should be satisfied. Here, $\mu_i^{\mathrm{d}}$ and $\mu_i^{\mathrm{c}}$ are the chemical potentials ($\mu$) of species $i$ in the two phases, respectively. Here, the chemical potentials of both the ssDNA chains and $Spm^{4+}$ ions should to be equal at coexistence. The chemical potential is estimated using the Widom insertion method,[24] deletion methods,[25] or a combination of the two.[25,26] In the

Widom method, a virtual test particle is introduced into a frozen snapshot of the system. The interaction energy ($U$) with existing particles is used to evaluate, $\mu = -\frac{1}{\beta}\ln\left\langle\exp\left(-\beta U\right)\right\rangle_N$ where $\beta = (k_B T)^{-1}$ and the average is over the $N$ particles. Although the Widom insertion method is readily implemented for simple fluids, this approach is challenging for chain molecules due to steric hindrance, resulting in low insertion probability. Alternative methods, such as segmental chemical potential[27] and configurational bias Monte Carlo,[28] address these issues but are computationally demanding for dense polyelectrolyte systems, where maintaining charge neutrality is nontrivial.[29]

The chemical potential $\mu$ is given by,

$$\mu = \left(\frac{\partial F}{\partial N}\right)_{T,V}. \tag{S21}$$

where $F$ is the Helmholtz free energy. To keep calculations of $\mu$ tractable in the ssDNA-PA systems, we estimated the free energy change $\Delta F$ by tracking the number of chains in the dense (dilute) phase, and measured the energy differences when $\Delta N = 1$, using the equilibrated conformations. Although this method favors low-energy states, applying it systematically to both the phases and averaging over chain conformations should reduce the bias. The resulting $\mu$ is the effective free energy cost of transferring a chain from one phase to another. In the dense phase, the interaction energies were computed for particles within the droplet. In the dilute phase, only particles beyond the droplet boundary and a buffer region (taken to be 10 nm) were considered to minimize the interfacial effects. Figures S2 and S3 present the distributions of the chemical potential ($\mu$) for the condensate and dilute phases in the $dT_{50}$ and $dA_{50}$ systems a function of $Spm^{4+}$ concentrations. The differences between the chemical potentials ($\Delta\mu$s) for the DNA chains sequences are small ($\approx 0$) in the range of $Spm^{4+}$ where phase separation occurs (Figures S2 and S3). The differences $\Delta\mu$ between the condensate and dilute phases is nearly zero (Figures S2 and S3), indicating that chemical equilibrium has been achieved. Because of larger fluctuations in $Spm^{4+}$ in both the phases the differences in their chemical potentials are greater.

The chemical potential distributions, especially for the ssDNA chains, overlap closely, as indicated by the Jensen-Shannon divergence (JSD). The symmetric JSD, a measure of difference between two probability distributions $P$ and $Q$, is defined as,

$$\text{JSD}(P \parallel Q) = \frac{1}{2} D_{\text{KL}}(P \parallel M) + \frac{1}{2} D_{\text{KL}}(Q \parallel M), \quad \text{where} \quad M = \frac{P+Q}{2}. \tag{S22}$$

The Kullback-Leibler divergence is given by,

$$D_{\text{KL}}(P \parallel M) = \sum_i P(i) \log \frac{P(i)}{M(i)}. \tag{S23}$$

The JSD values are bounded between 0 and 1, being 0 when P = Q and 1 when the distributions are maximally different. The values of the fluctuations in $\mu$ ( $\sigma_\mu = \sqrt{\langle \mu^2 \rangle - (\langle \mu \rangle)^2}$) in $\mu$ (Figures S2 and S3 decreases as the $\text{Spm}^{4+}$ concentration increases.

**Dynamic equilibrium between the phases:** Another measure of equilibrium is to ensure there are frequent chain exchange between the two phases. Let the distance between the center of mass (COM) of a chain and the COM of the condensate be,

$$r_i(t) = \left| \mathbf{R}^{\text{c}}_{\text{COM}}(t) - \mathbf{R}^{i}_{\text{COM}}(t) \right|, \tag{S24}$$

where $\mathbf{R}^{\text{c}}_{\text{COM}}(t)$ is the position of the condensate center at time $t$, and $\mathbf{R}^{i}_{\text{COM}}(t)$ is the COM of the $i^{\text{th}}$ chain. If $r_i(t)$ is less than $R_D$, the radius of the droplet, it implies that the $\mathbf{R}^{i}_{\text{COM}}(t)$ is within the condensate. Otherwise the chain is in the dispersed (dilute) phase. By calculating $r_i(t)$ as a function of time for representative chains, one can observe multiple transitions between the condensed and dilute states (Figure S18). The time dependent variations in $r_i(t)$ gives an estimate of the residence time of a chain in the condensate. Upon averaging $r_i(t)$ over all the chains in the droplet the distribution of lifetimes can be calculated. In the condensates the mean residence time is between (15 - 30)$\tau$ (Figure S18 D and H). Thus, there are frequent exchanges of the polymers between the two phases on the time scales in which the droplet forms. The equilibrated droplets exhibit fluctuations in the number of chains, with $N_c$ sampling values around a stable mean (Figures S18A and E). Cluster-size analysis further shows that individual chains frequently switch between monomeric, small cluster, and large cluster states, indicating continuous exchange between these conformational states (Figures S18B and F).

**Finite size effects:** We assessed finite-size effects by varying the number ($N$) of $dA_{50}$ chains at 350 $\mu$M DNA concentration in 10 mM $Spm^{4+}$. For computational reasons, we varied $N$ only over a modest range ($N = 32$ to $N = 160$). Although there are variations in the monomer density distribution of $P(\rho_c)$ in the dense phase at small $N$ (Figures S7A-E), at the two highest $N$ it seems that $P(\rho_c)$ is converged. The fluctuation in $\rho_c$ (denoted by $\sigma_c = \sqrt{\langle \rho_c^2 \rangle - (\langle \rho_c \rangle)^2}$) decreases as $N$ increases (Figure S7F), in accord with statistical mechanics. Similarly, Figures S7G-K show the system-size dependent changes in the distributions of monomer concentrations in the dilute phase, $P(\rho_d)$. With increase in $N$, the fluctuations (denoted by standard deviation $\sigma_d = \sqrt{\langle \rho_d^2 \rangle - (\langle \rho_d \rangle)^2}$) decrease. Figure S7M shows transfer free energies $\Delta G_{tr}$ as a function of $N$. Both concentrations and $\delta G_{tr}$ show negligible variations when $N \geq 90$. Because the observed finite-size effects are modest even at $N = 64$, we constructed phase diagram with $N = 64$ to preserve computational expediency.

$Spm^{4+}$ **dependent changes in the two phases:** Figures S5 shows that the average values of $\rho_c$ in the condensate phase decrease sharply as $Spm^{4+}$ increases. At comparable $Spm^{4+}$ levels, $dA_{50}$ droplets contain a larger fraction of chains in the dense phase than $dT_{50}$ (see Figures S20). The density distribution in the dilute phase are broad with the mean that increases with increasing $Spm^{4+}$ concentration in (Figure S6 and Table S1). The average values of $\rho_d$ increase only modestly as the $Spm^{4+}$ increases (Table S1). Using Eqs. S12, we calculated the radius of gyration ($R_g$), as well as the number of ssDNA chains $N_c$ in the $dT_{50}$ and $dA_{50}$ condensates as function of $Spm^{4+}$ concentration (Figures S20, S21). For both the sequences, with increase in $Spm^{4+}$ concentrations, the number of chains in the condensate phase $N_c$ decreases, which is harbinger of the reentrant phase (see Figure 2K in the main text).

**Role of ions in the droplets:** To determine the role of ions in rendering the droplets neutral, we calculated the concentration profiles of $P^-$, $N^+$, and $Cl^-$ using,

$$\rho_X(r) = \frac{1}{N_A V(r)} \int_0^r n_X(r)\, 4\pi r^2 dr. \tag{S25}$$

where $r$ is the distance from the condensate COM. In Eq. S25 $n_X(r)$ is the number density of ion $X$ at a distance $r$ and $V(r) = \frac{4}{3}\pi r^3$, and $N_A$ is Avogadro's number. Figures 2B and D in main text show that at $Spm^{4+}$ concentrations ($\approx$ 10 mM), condensates are neutralized primarily by $Spm^{4+}$ amine groups. The $Cl^-$ ions neutralize presence of excess $Spm^{4+}$ ions. Needless to say, the condensates are charge neutral across at all $Spm^{4+}$ concentrations.

**Condensate growth mechanism:** The main text shows $dT_{50}$ droplets grow predominantly by the addition of diffusing monomers. In contrast, droplet growth in $dA_{50}$ occurs by merging of oligomers. The small oligomers that form at the early stages are devoured by larger droplets, which is reminiscent of Ostwald ripening. Similar mechanisms are noted in Figure S15 in four additional independent trajectories (Figure S15). The number of chains in the largest cluster, $N(t)$ (shown in purple) in $dT_{50}$ increases continuously with time (Figures S15A,C,D, and E). The change, $\frac{dN(t)}{dt}$ (upper panel) is roughly a constant, fluctuating around unity, which implies that the chain growth must occur by addition of one or few monomers at a time to a growing droplet. Figures S15B,D,F and G show fusion of two oligomers in $dA_{50}$ condensate growth. The quantized increase in the droplet size is reflected in the abrupt change in $\frac{dN(t)}{dt}$, showing a jump in the slope over a short time interval. Notably, fusion events in $dA_{50}$ are prominently observed in simulations in larger system sizes ($N = 160$), indicating that such coalescence dynamics are sensitive to finite-size effects.

**Condensate shape dynamics:** We expect that at long times the droplets have to be spherical in order to minimize the surface tension, regardless of the sequence. In accord with this expectation, the shape anisotropy $\kappa^2$ ( Eq. S14) of the largest ssDNA cluster (Figures S14A and B) decreases as function of time as the condensate matures and reaches steady state. As phase separation progresses, the size of the largest cluster grows. At times $\tau < 4,000\tau$ in $dT_{50}$ and $\tau < 2,000\tau$ in $dA_{50}$ there are substantial fluctuations in $\kappa^2$. At longer times, the droplets are stable and become spherical ($\kappa^2 \approx 0$). The time to reach sphericity as well as the steady state values of $\kappa^2$ are sequence dependent, reflecting the different growth mechanisms (see also Figure S17). The dependence of $\kappa^2$ on the cluster in Figures S14C,D illustrate that larger clusters are more spherical. The distributions of shape anisotropy, $P(\kappa^2)$ in Figures S14E,F show that as $Spm^{4+}$ concentrations increase the widths of $P(\kappa^2)$ increases.

**Impact of Weak NN Stacking Interactions:** The contribution of non-neighboring (NN) inter-chain stacking interactions to the droplet stability is only on the order of a few $k_BT$ each (see Figure S13). Nevertheless, several such contacts form spontaneously in the condensate, contributing to stabilize the dense phase. The average NN stacking energy per ssDNA chain decreases with increasing $Spm^{4+}$ concentration beyond 5mM. Interestingly, at the same $Spm^{4+}$ concentration, inter chain stacking in $dA_{50}$ condensates are almost twice as large in the $dT_{50}$ condensates. The enhanced cooperative contribution of NN stacking explains the increased $Spm^{4+}$ reentrant phase forms $dA_{50}$. In the dispersed phase, $U_{ST}^{NN}$ per $dT_{50}$ ($dA_{50}$) chain is ≈ -0.5 (-0.7) $k_BT$, which is 7.2 (9) times weaker compared to condensate phase. The $U_{ST}^{NN}$ values between dispersed and reentrant phases are comparable.

### Additional experimental results:

**DNA concentration dependent phase transition of ssDNA in spermine complex coacervates:** We tested how phase-stability of ssDNA/spermine complex coacervates varies with DNA concentration. To this end, we prepared mixtures with a fixed concentration of $Spm^{4+}$ (2.5 mM) and titrated DNA from 25 to 150 $\mu$M. $dT_{50}$/spermine mixtures again displayed a reentrant phase-behavior, now as a function of DNA concentration. Complex coacervates formed only at intermediate DNA concentrations (50- 100 $\mu$M DNA concentration 1:40 to 1:1.33 charge ratio, Figure S22A). Outside of this range $dT_{50}$ chains did not phase separate and remained in solution (Figure S22B). In contrast, $dA_{50}$ chains formed complex coacervates at all tested DNA concentrations (1:8 to 1:1.33 charge ratio). Under these conditions, the soluble $dA_{50}$ fraction was consistently lower than that of $dT_{50}$, again indicating that a larger fraction of $dA_{50}$ chains partitioned into the complex coacervate phase across the entire DNA concentration range.

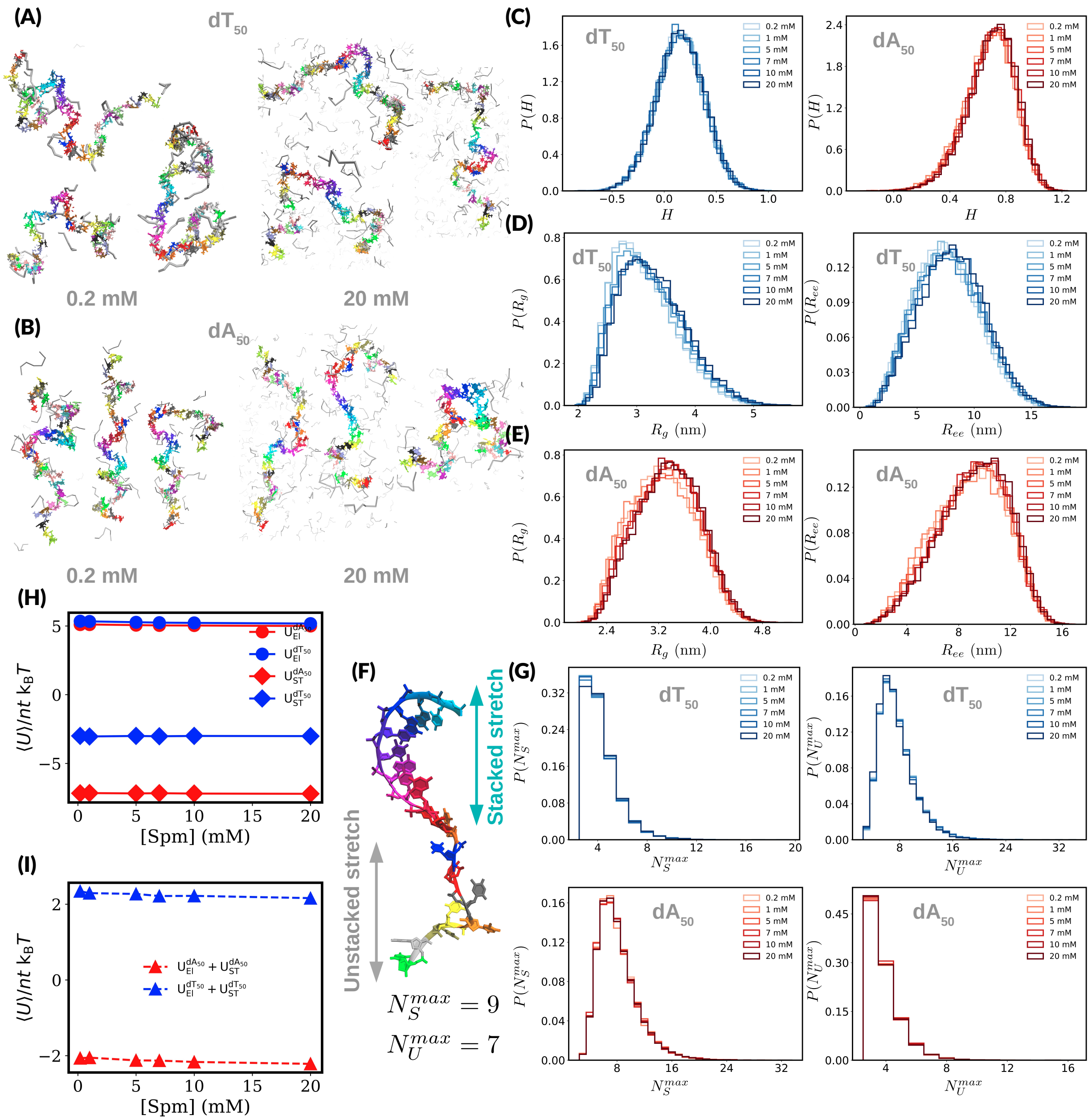


Figure S1: [Spm$^{4+}$ ] **dependent features of monomers:** Representative conformations of (A) dT$_{50}$ and (B) dA$_{50}$ in 0.2mM and 20 mM of Spm$^{4+}$ . Individual residues are color-coded for clarity. (C) Distributions of helix content $H$, (D) $R_g$, and (E) $R_{ee}$ as a function of Spm$^{4+}$ concentrations.(F) Schematic of the longest stretches of persistent stacked, and unstacked nucleobases. Distributions of longest stretches of (G) stacked, and unstacked nucleobases in dT$_{50}$ and dA$_{50}$ as function of [Spm$^{4+}$ ].(H) Potential energy per nucleotide due to electrostatic repulsion between phosphate groups (circles). Stabilizing base stacking interactions including both adjacent and non-adjacent bases (diamonds). (I) Total potential energy contribution to the stability of ssDNA per nucleotide.

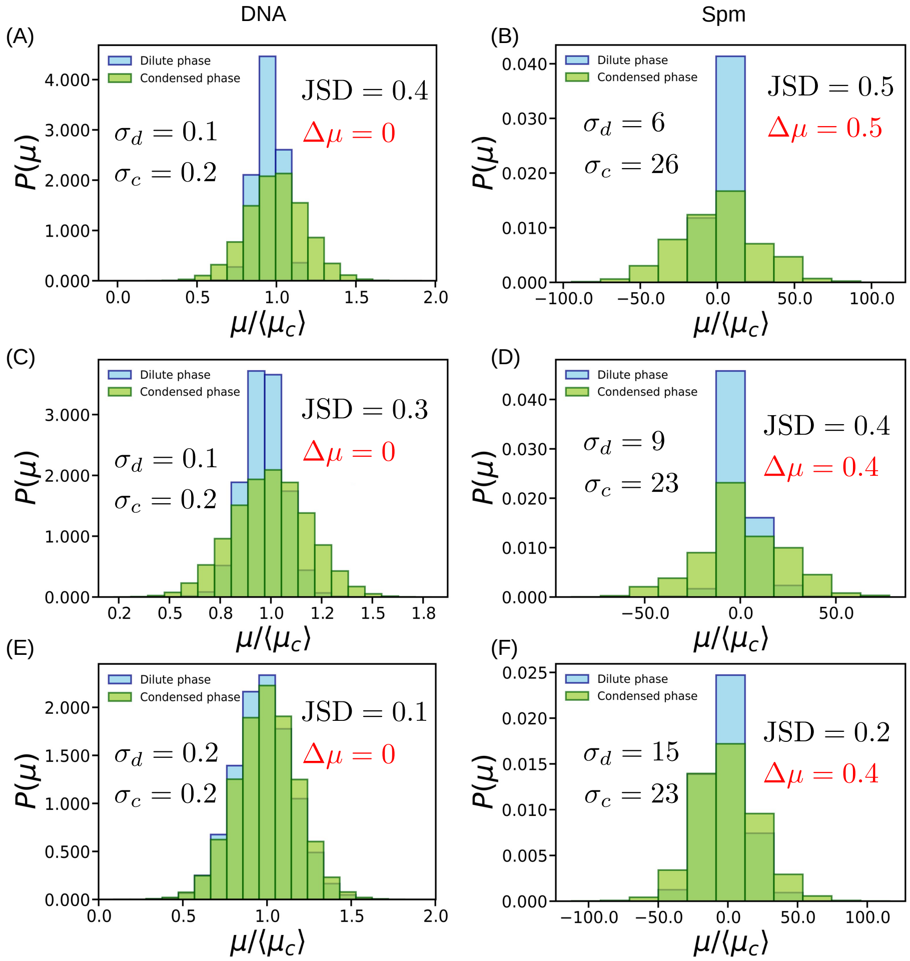


Figure S2: **Phase equilibrium in** $dT_{50}$ **:** Distributions of the chemical potentials ($\mu$s) for $dT_{50}$ and $Spm^{4+}$ in condensed and dilute phases. The $\Delta\mu$ values are within the standard deviation of the $\mu$ distributions for both the species, ensuring that chemical equilibrium is reached. The difference $\Delta\mu$ between the two phases for DNA and $Spm^{4+}$ is in red. Results are for (A, B) [Spm] = 4 mM, (C, D) [Spm] = 7mM and (E, F) [Spm] = 10 mM. The Jensen-Shannon divergence (JSD) values and the fluctuations in concentrations in dilute ($\sigma_d$) and condensate ($\sigma_c$) phases are listed.

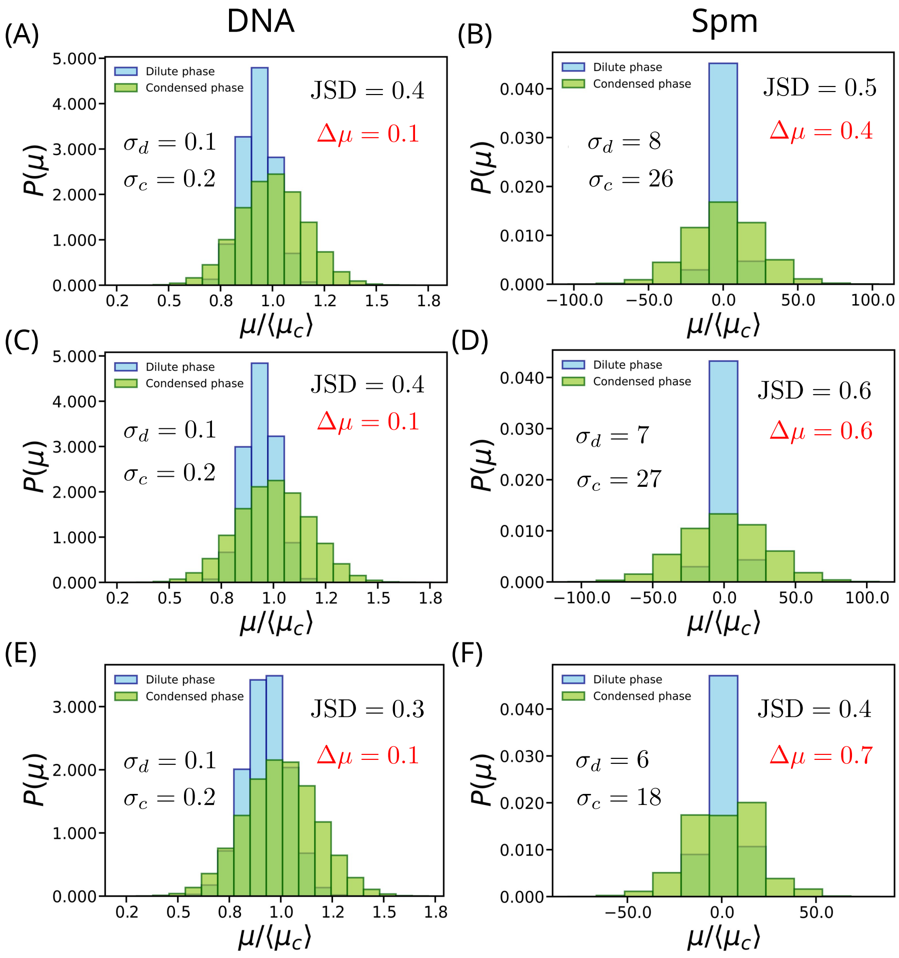


Figure S3: **Phase equilibrium in** $dA_{50}$**:** Establishing chemical equilibrium in the phase separation of $dA_{50}$. See the caption in Figure S2 for an explanation. Results correspond to (A, B) [Spm] = 4 mM, (C, D) [Spm] = 7 Mm, and (E, F) [Spm] = 10mM.

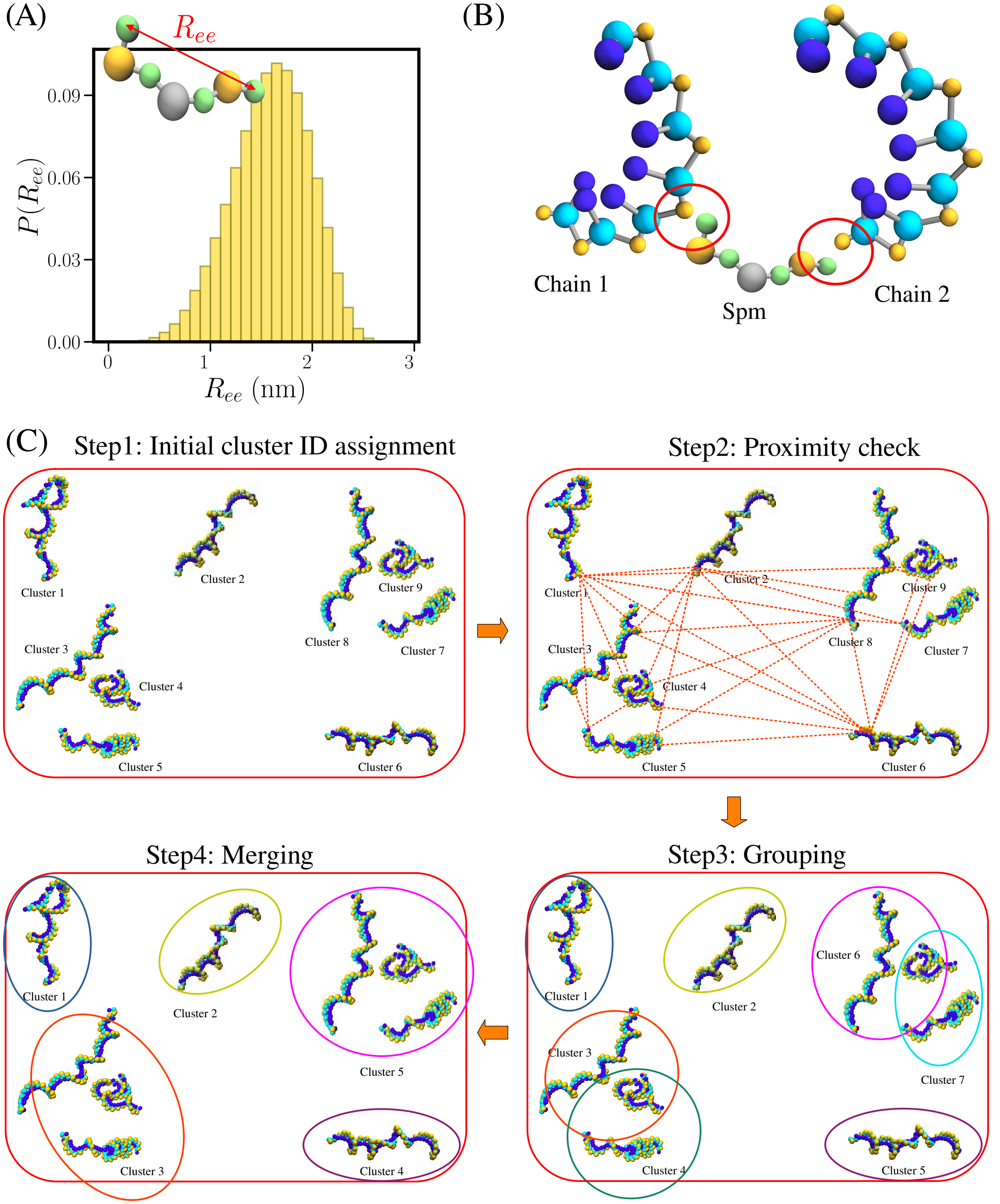


Figure S4: **Clustering algorithm:** (A) End-to-end distance distribution, $P(R_{ee})$, for $Spm^{4+}$. (B) A schematic of a Spm molecule bridging two ssDNA chains. (C) Flowchart for the clustering algorithm. Each chain is initially assigned a unique cluster ID. The closest distances between all the pairs of chains are computed to assess proximity. Clusters are iteratively merged if coarse-grained beads from different chains are within a specified cutoff distance. This process continues until all chains are grouped into non-overlapping clusters.

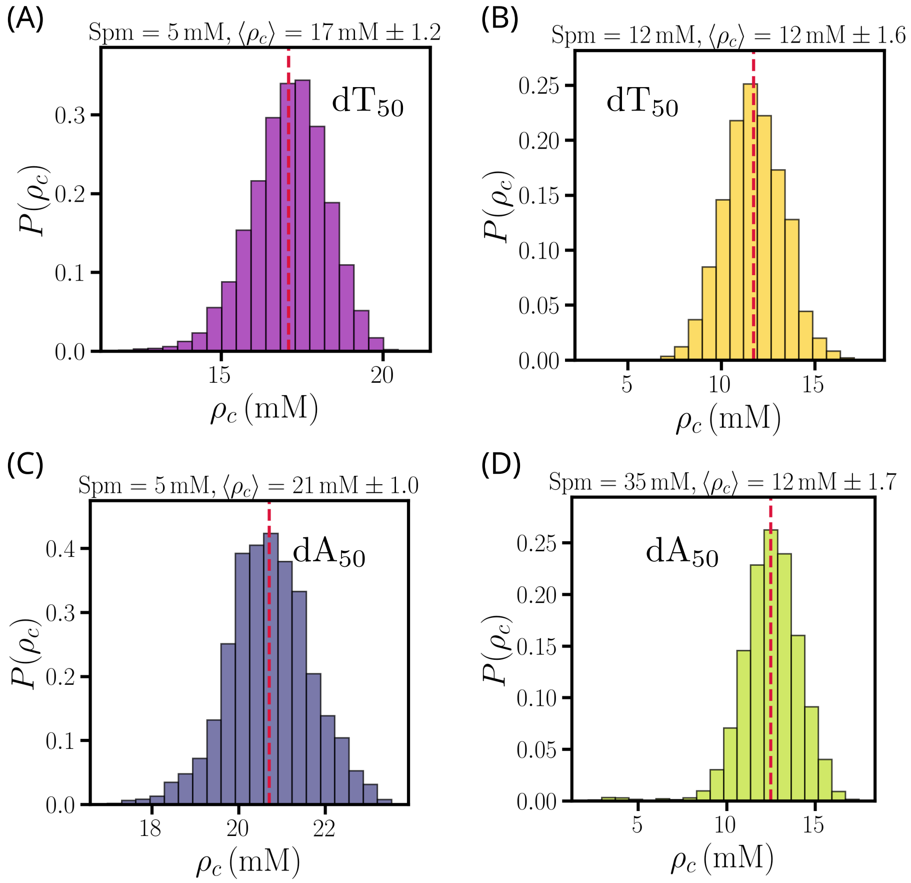


Figure S5: **Density distribution in the condensed phase:** (A-D) Distribution $P(\rho_c)$ of monomer concentrations, $\rho_c$, as a function of Spm concentrations. Vertical dashed lines are the average values (Table. S1 lists numerical values).

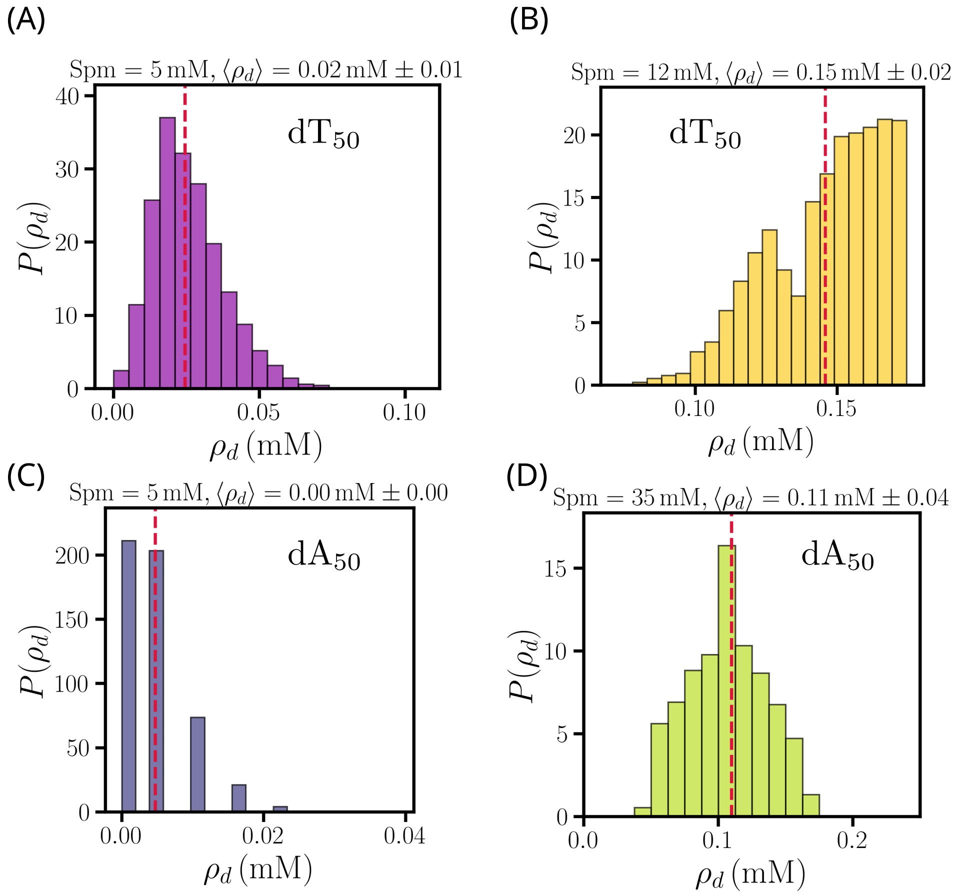


Figure S6: **Density distribution in the dilute phase:** (A-D) Distribution $P(\rho_d)$ of monomer concentration in the dilute phase , $\rho_d$, in different Spm concentrations. Vertical dashed lines specify the average values ( Table. S1 for numerical values).

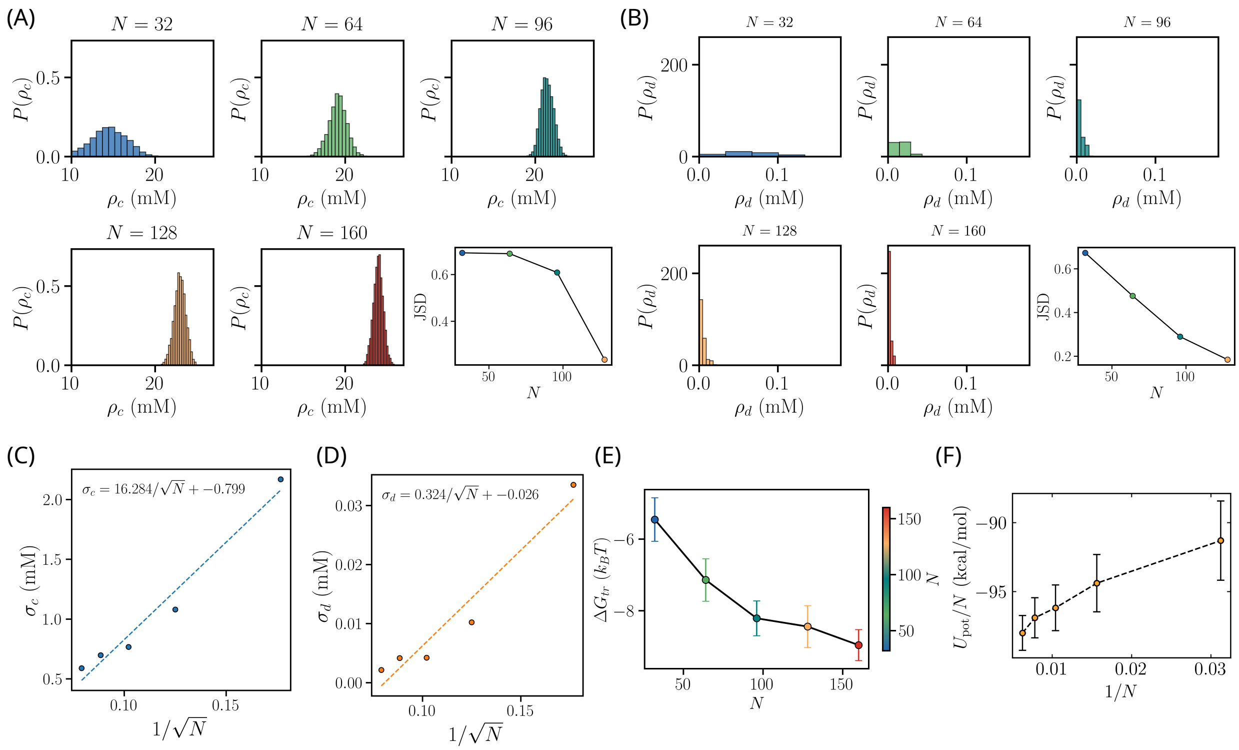


Figure S7: **Finite size effects in** $dA_{50}$ **at 10 mM** $Spm^{4+}$**:** (A) Distributions of concentration of the ssDNA chains in the condensed phase for different number of chains $N$. Jensen-Shannon divergence (JSD) values are plotted with respect to $N = 160$. (B) Same as (A), except the distributions are plotted for concentration of the chains in the dilute phase. (C, D) Standard deviation in the dense ($\sigma_c$) and dilute phase concentrations ($\sigma_d$) versus $1/\sqrt{N}$. Fits to $\sigma_c = a_c/\sqrt{N} + b_c$ and $\sigma_d = a_d/\sqrt{N} + b_d$ (coefficients shown in the plot) show approximately linear dependence and small intercepts, confirming the expected $\sigma \propto 1/\sqrt{N}$. (E) Transfer free energies, $\Delta G_{tr}$ as a function of $N$. (F) Total potential energy per DNA chain, as a function of $1/N$.

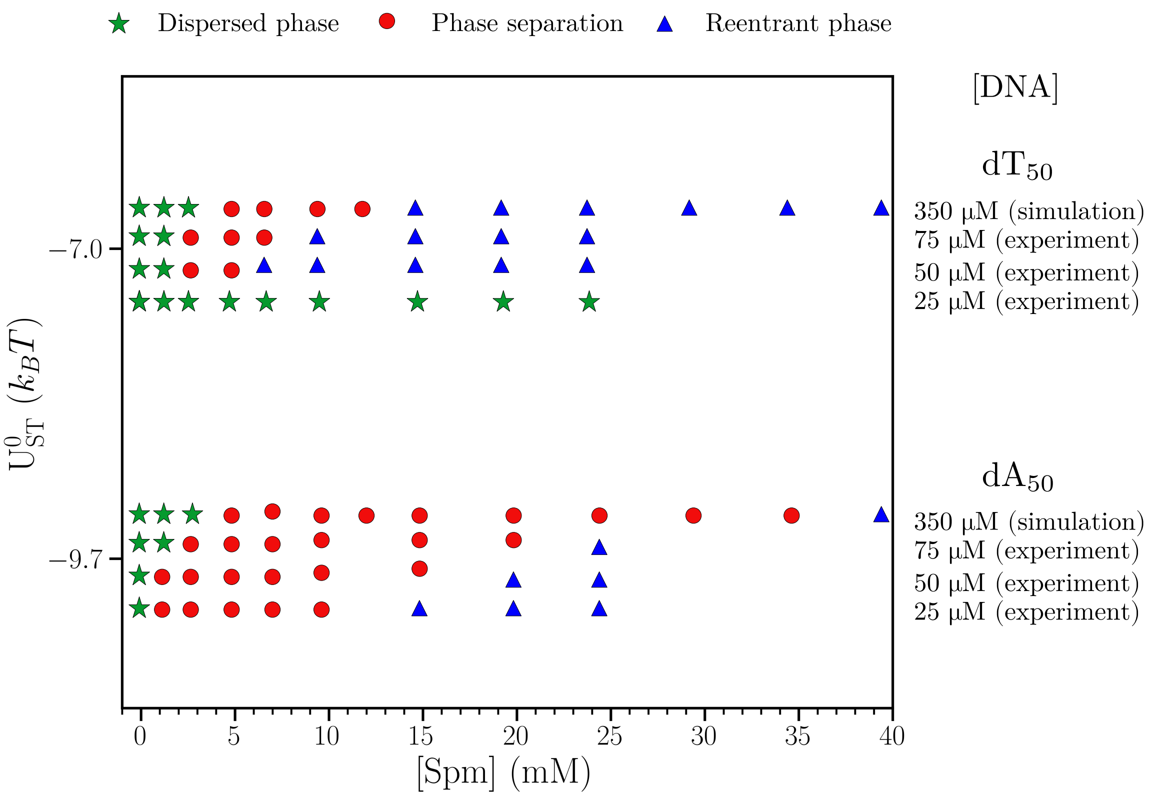


Figure S8: Phase diagram of $dT_{50}$ and $dA_{50}$ in different DNA concentrations.

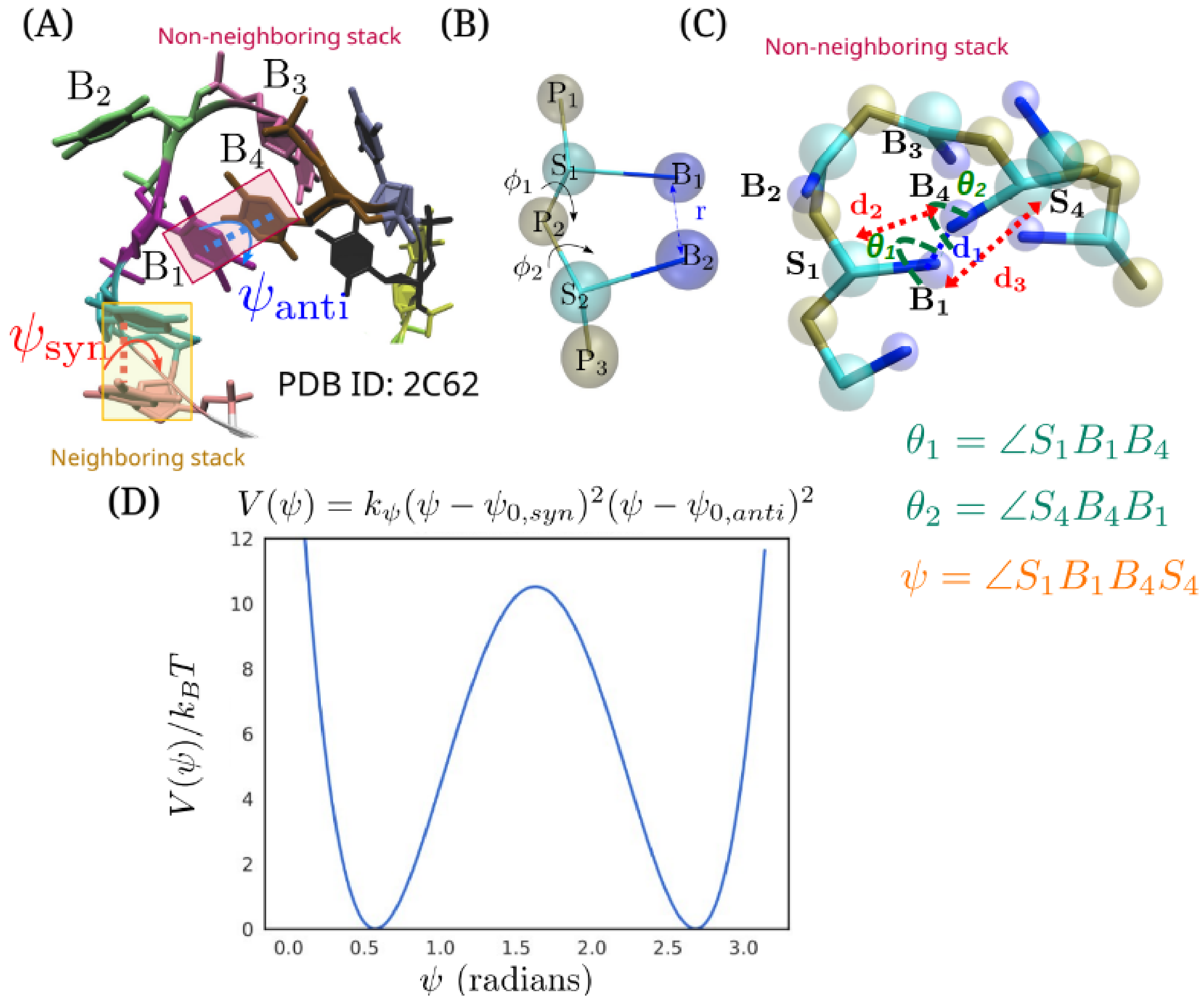


Figure S9: **Non-neighboring (NN) stacking interactions:** (A) *Syn* and *anti* conformations of the dihedral angle $\psi(S_iB_iB_jS_j)$ formed by two stacked bases $B_i$ and $B_j$ illustrated using a fragment of ssDNA from PDB ID: 2C62. $B_1$ and $B_4$, separated by two other bases $B_2$ and $B_3$, form non-neighboring (NN) stacks. (B) Pictorial representation of near neighbor base stacking interactions. The centers of phosphate (P), sugar (S), and base (B) groups, are in tan, cyan, and blue colors, respectively. (C) Coarse-grained representation of the geometry of NN base stacking in the structure in (A). (D) Double-well potential $V(\psi)$ (the last term in Eq S1) for the NN stacking interaction. The values of $\psi_{0,syn}$, $\psi_{0,anti}$, and $k_\psi^{NN}$ are listed in Table S3.

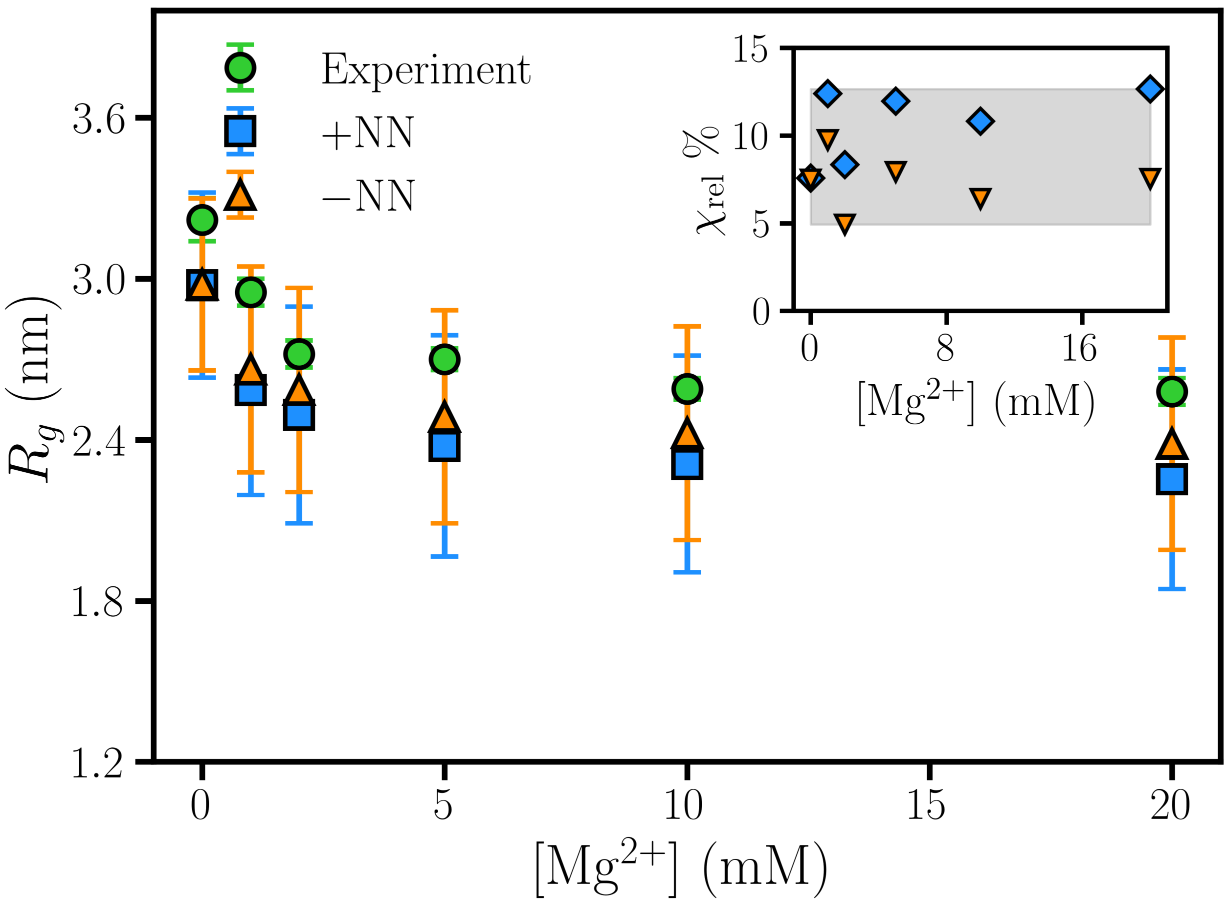


Figure S10: Mg$^{2+}$ **dependent radius of gyration ($R_g$)** dT$_{30}$ **monomer:** Experimental[30] (green circles) and simulated $R_g$ values (blue squares with NN and orange triangles without NN stacks) in 20 mM NaCl and varying Mg$^{2+}$ concentrations. Inset shows relative error, $\chi_{rel}, = \left(\frac{|R_g^{\rm sim} - R_g^{\rm exp}|}{R_g^{\rm exp}}\right) \times 100\%$.

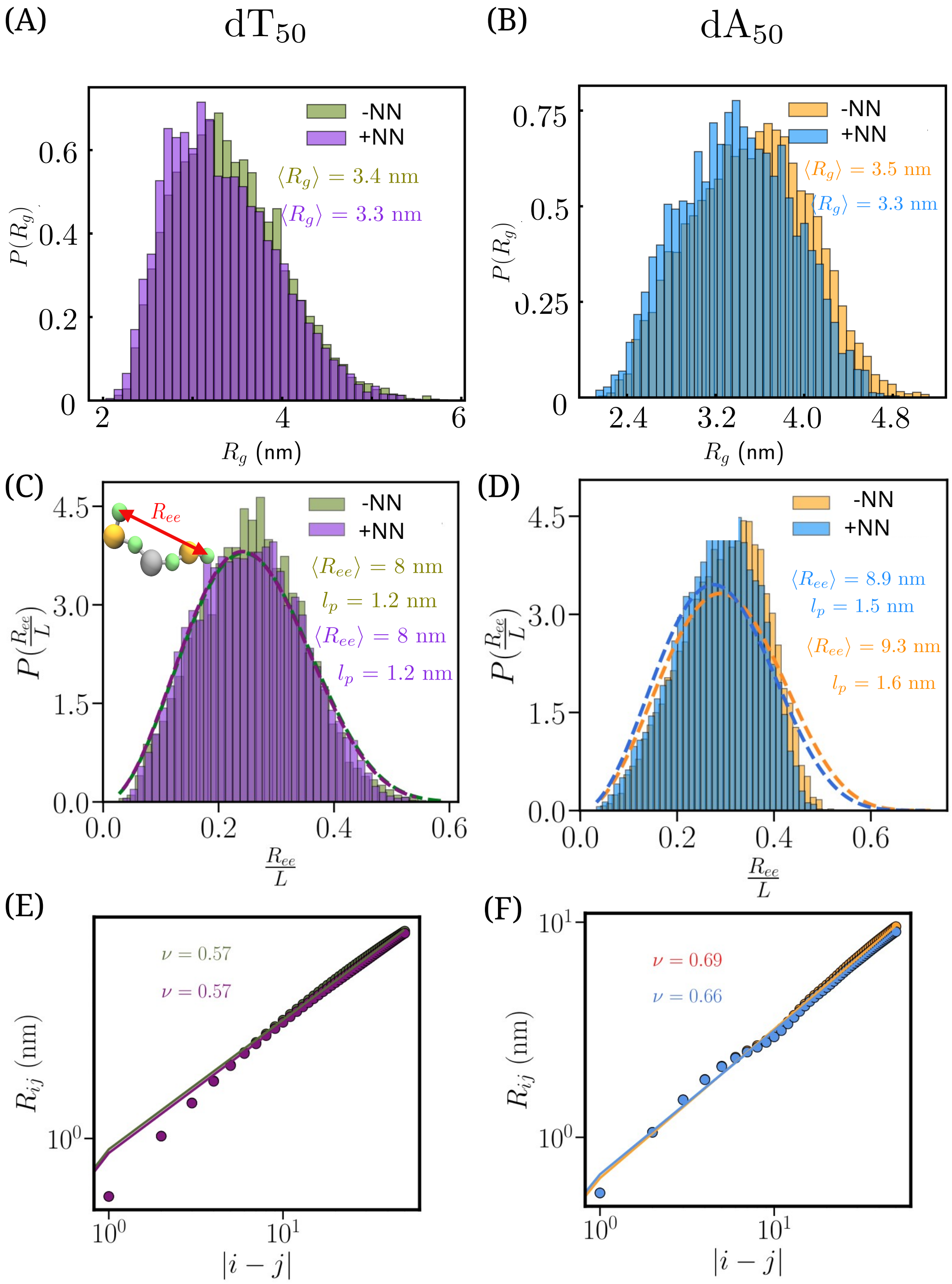


Figure S11: **Monomer characteristics:** (A) and (B): Distribution of $P(R_g)$s of $R_g$ for $dT_{50}$ (left) and $dA_{50}$ (right) in 10 mM $Spm^{4+}$ with and without non-adjacent stacking interactions. (C) and (D): Normalized end-to-end distance, $R_{ee}/L$ distributions ($P(R_{ee}/L)$s) where $L$ is the contour length. The dashed lines are fits to Eq. S4. Average values of $R_g$, $R_{ee}$, and $l_p$ are shown. (E) and (F): Effective Flory exponent $\nu$ calculated using Eq. S5 for $dT_{50}$ (left panel) and (F) $dA_{50}$ (right panel).

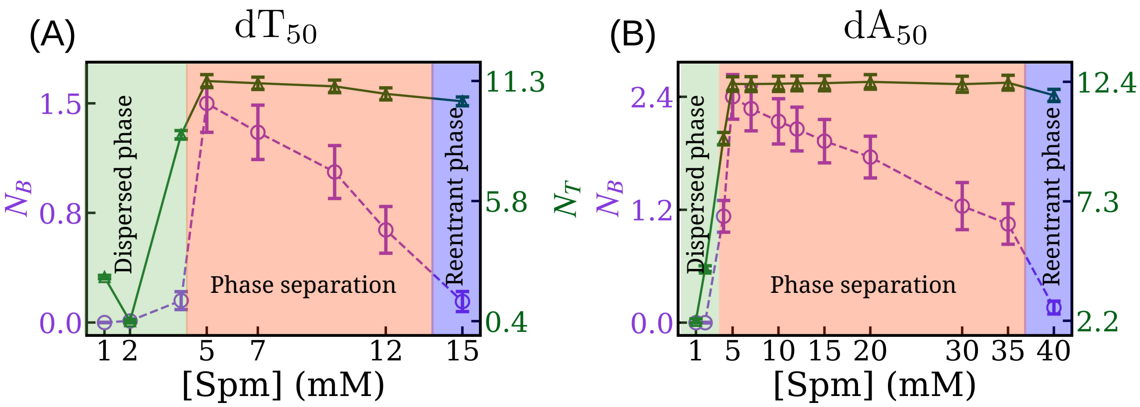


Figure S12: **Dependence of bridging and total spermine per DNA chain on spermine concentration:** Average number of bridging Spm ($N_B$) and average number of total Spm ($N_T$) per ssDNA chain, plotted as a function of [Spm].

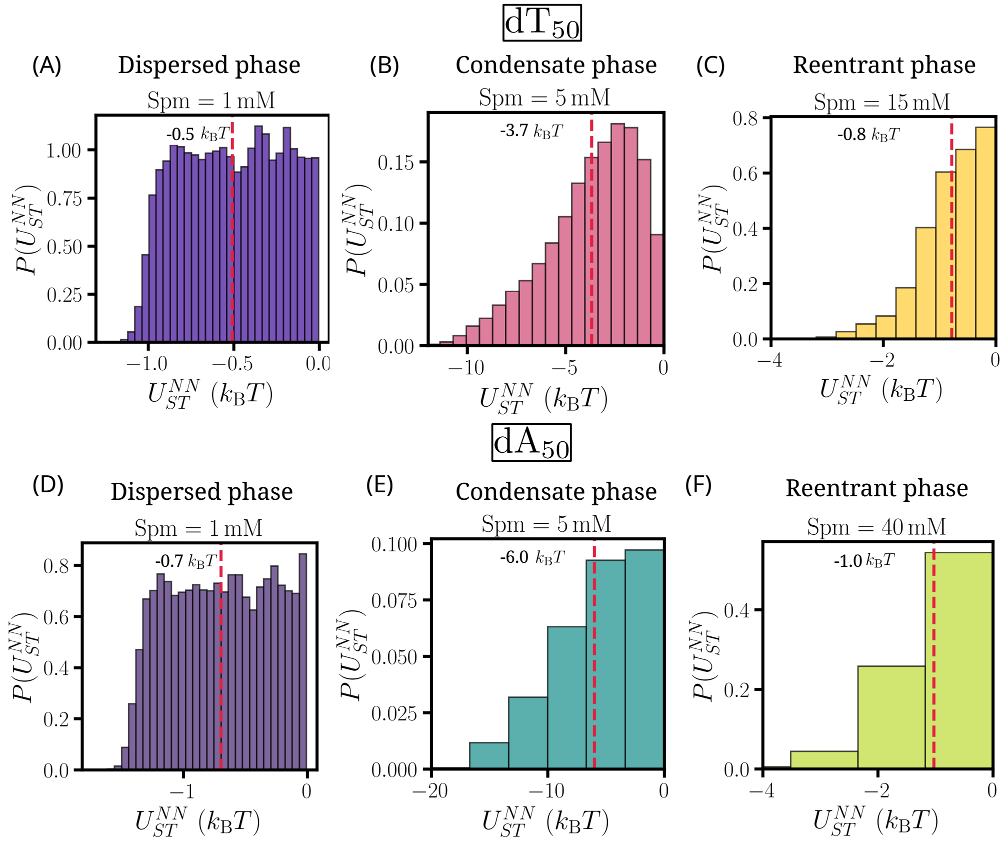


Figure S13: **Inter-strand stacking energy, $U_{ST}^{NN}$:** (A-C) Distributions of inter-strand stacking energy, $U_{ST}^{NN}$, per ssDNA chain. Different Spm concentrations correspond to (A) dispersed, (B) condensate, and (C) reentrant phases. Red dashed lines the average values. Results are shown for dT$_{50}$. (D-F) Same as (A-C), except results are shown for dA$_{50}$.

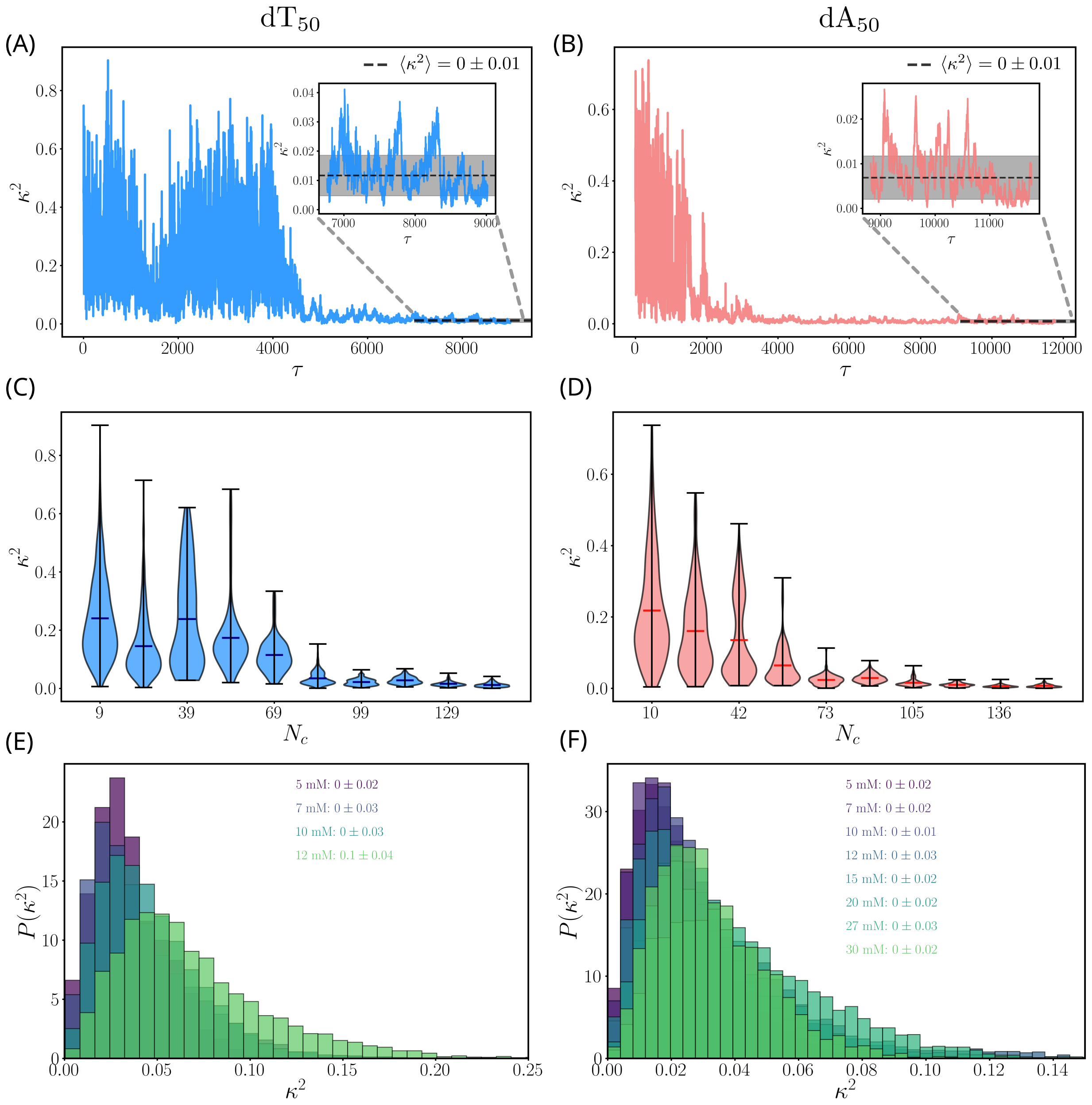


Figure S14: **Condensate shape evolution:** (A, B) Time evolution in shape anisotropy, $\kappa^2$ ( Eq. S14) in the largest cluster. $\kappa^2 = 0$ for a perfect sphere, and approaches 1 for rod-like structures. Inset highlight $\kappa^2$ after the onset of equilibrium. Dashed black lines indicate $\langle\kappa^2\rangle$ and the shaded regions show fluctuations ($\sigma_{\kappa^2} = \sqrt{\langle(\kappa^2)^2\rangle - \langle\kappa^2\rangle^2}$) in $\kappa^2$. (C, D) Distribution of $\kappa^2$ as a function of condensate size $N_c$. The results for the ssDNA sequences in [Spm$^{4+}$]=10 mM and $N$= 160. (E, F) Distributions of $\kappa^2$ as a function of Spm$^{4+}$ concentrations. Values of $\langle\kappa^2\rangle$ are shown. Results are for $N$= 64. For both dT$_{50}$ and dA$_{50}$ condensates, $\langle\kappa^2\rangle \approx 0$ at long times. As the spermine concentration increases, the distributions broaden, reflecting a progressive increase in condensate deformation.

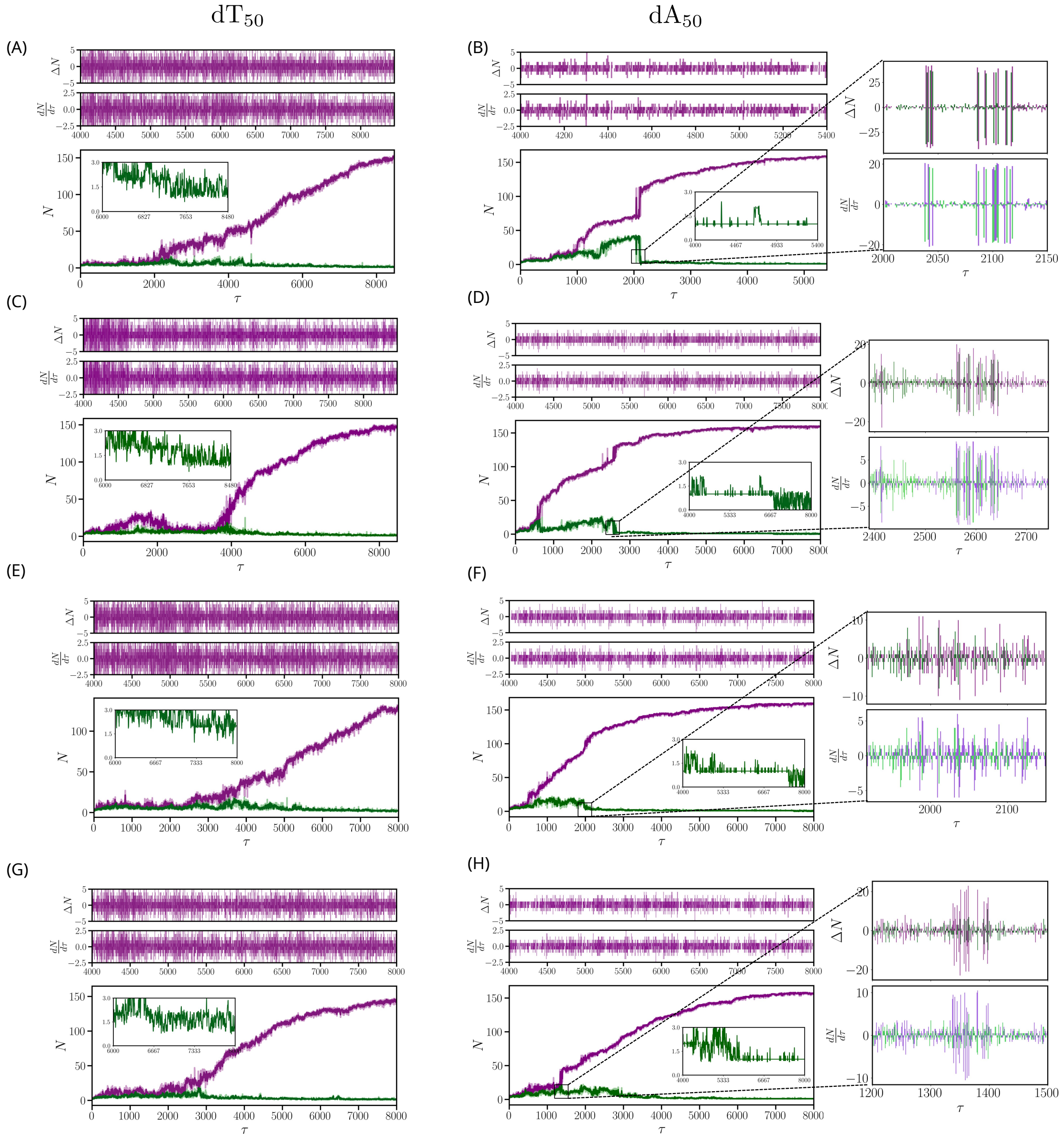


Figure S15: **Droplet formation dynamics:** (A,C,E,G) Time evolution of two largest clusters in $dT_{50}$ at 10 mM $Spm^{4+}$ from four trajectories. Purple shows that the number of chains in the condensate increases monotonically with time. Change in the number of chains ($\Delta N$) and rate of change ($\frac{\Delta N}{dt}$) are constant (see the upper panels). (B,D,F,H) Growth of the $dA_{50}$ condensate (in purple) as a function of time. In these trajectories smaller droplets fuse to form larger droplets. The fusion events are reflected in both $\Delta N$ and $\frac{\Delta N}{dt}$ (upper panel), which show an abrupt change over a narrow time interval. The number of chains in simulations is $N = 160$.

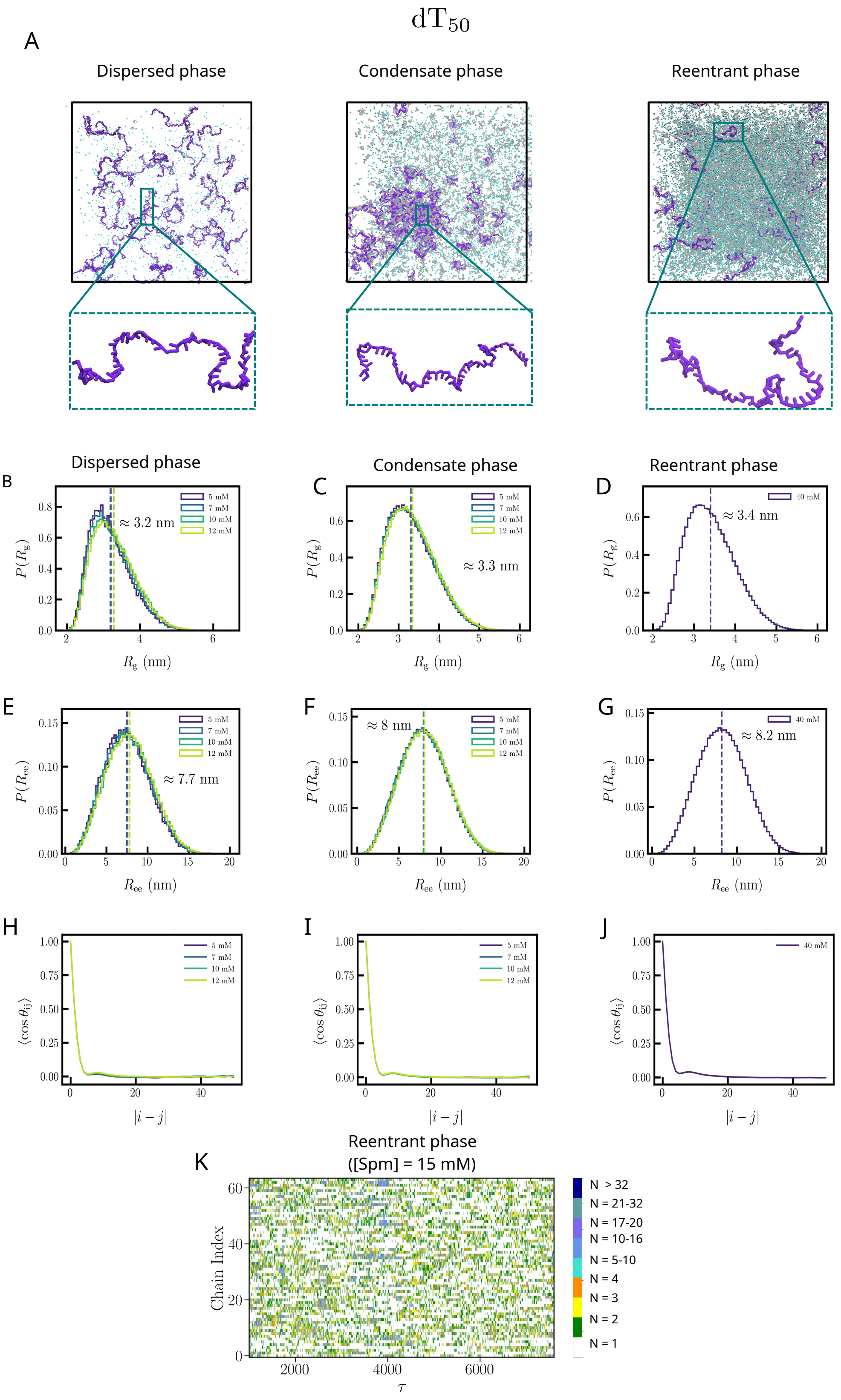
dT50
A
Dispersed phase
Condensate phase
Reentrant phase
B
C
D
≈ 3.2 nm
≈ 3.3 nm
≈ 3.4 nm
5 mM
7 mM
10 mM
12 mM
40 mM
P(Rg)
Rg (nm)
E
F
G
≈ 7.7 nm
≈ 8 nm
≈ 8.2 nm
P(Ree)
Ree (nm)
H
I
J
⟨cos θij⟩
|i − j|
K
Reentrant phase
([Spm] = 15 mM)
Chain Index
τ
N > 32
N = 21-32
N = 17-20
N = 10-16
N = 5-10
N = 4
N = 3
N = 2
N = 1

Figure S16

**Figure S16 caption:** Monomer features in the three phases: (A) Representative conformations of $dT_{50}$ chains in dispersed (1 mM Spm), condensate (10 mM Spm) and reentrant phases (15 mM Spm). $dT_{50}$ chains are shown in purple. Spm molecules are shown in silver. (B-D) Distributions of radius of gyration, $R_g$, of $dT_{50}$ monomers are shown in the dilute (B), the condensed (C) and the reentrant phases (D). Corresponding Spm concentrations are indicated in the legends. The dashed lines indicate average values of $R_g$ obtained from the distributions. $\langle R_g \rangle$ is reported. (E-G) End-to-end distance distributions, $P(R_{ee})$, of $dT_{50}$ monomers, in the dilute (E), the condensed (F) and reentrant phases (g) are plotted. The mean values of $\langle R_{ee} \rangle$, corresponding to the dashed lines, are listed. (H-J) Ensemble averaged angular correlation function, $C_n$ (Eq. S12 in the SI), as a function of $|i - j|$ for $dT_{50}$ at different [$Spm^{4+}$ ]. The panels from the left to right are for the dilute (h), droplet (I), and the reentrant (J) phases. $C_n$ for $dT_{50}$ decays rapidly indicating lack of order in the sugar-phosphate backbone. This trend is observed across the Spm concentrations and phases. The average values, shown as vertical lines, are listed in each panel. (K) Chain dynamics in the reentrant phase of $dT_{50}$. Individual chains are represented by separate lines, with their states color-coded. Higher order oligomers containing upto 17 to 20 chains (color coded blue) are observed occasionally in addition to smaller assemblies.

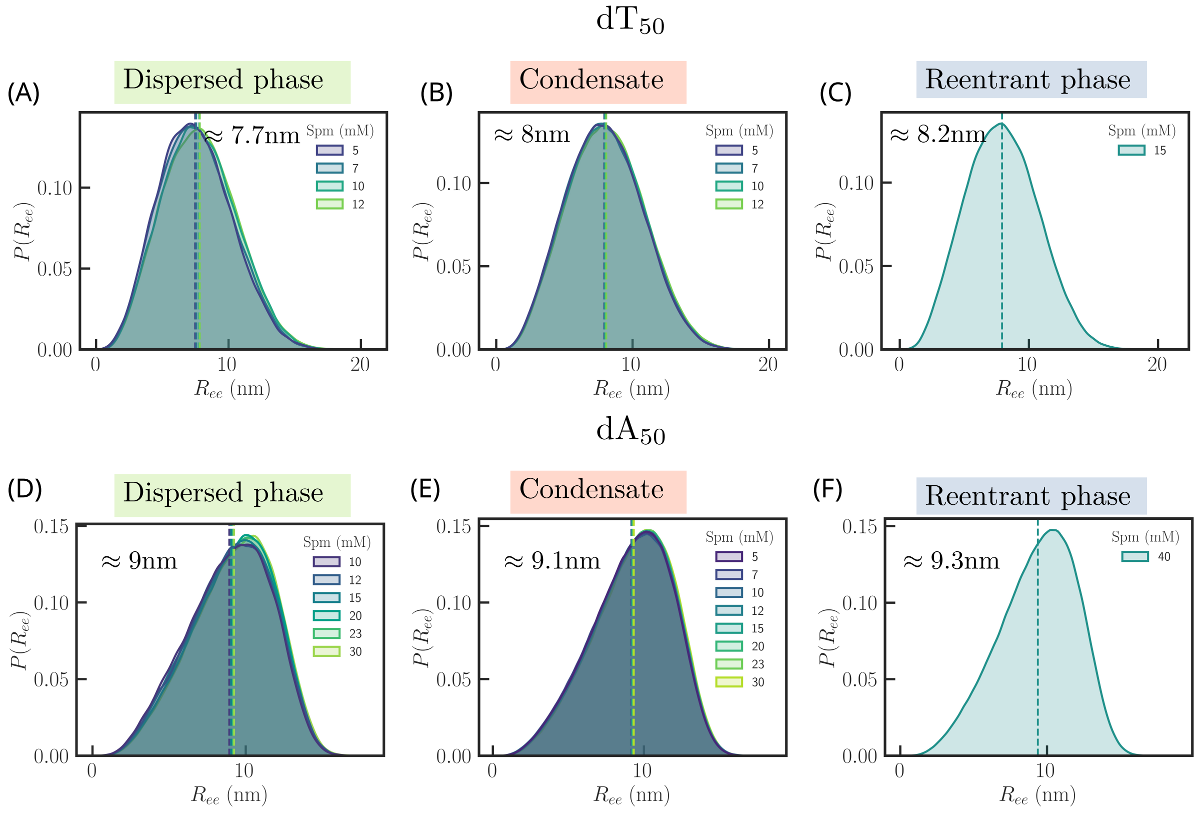


Figure S17: **Monomer end-to-end distance ($R_{ee}$) remains invariant across phases:** Distributions of $R_{ee}$ in the dispersed phase (A), condensate (B) and (C) in the reentrant phase for dT$_{50}$ chains. (D-E) Same as (A-C), except the results are for dA$_{50}$. $\langle R_{ee} \rangle$ values are reported.

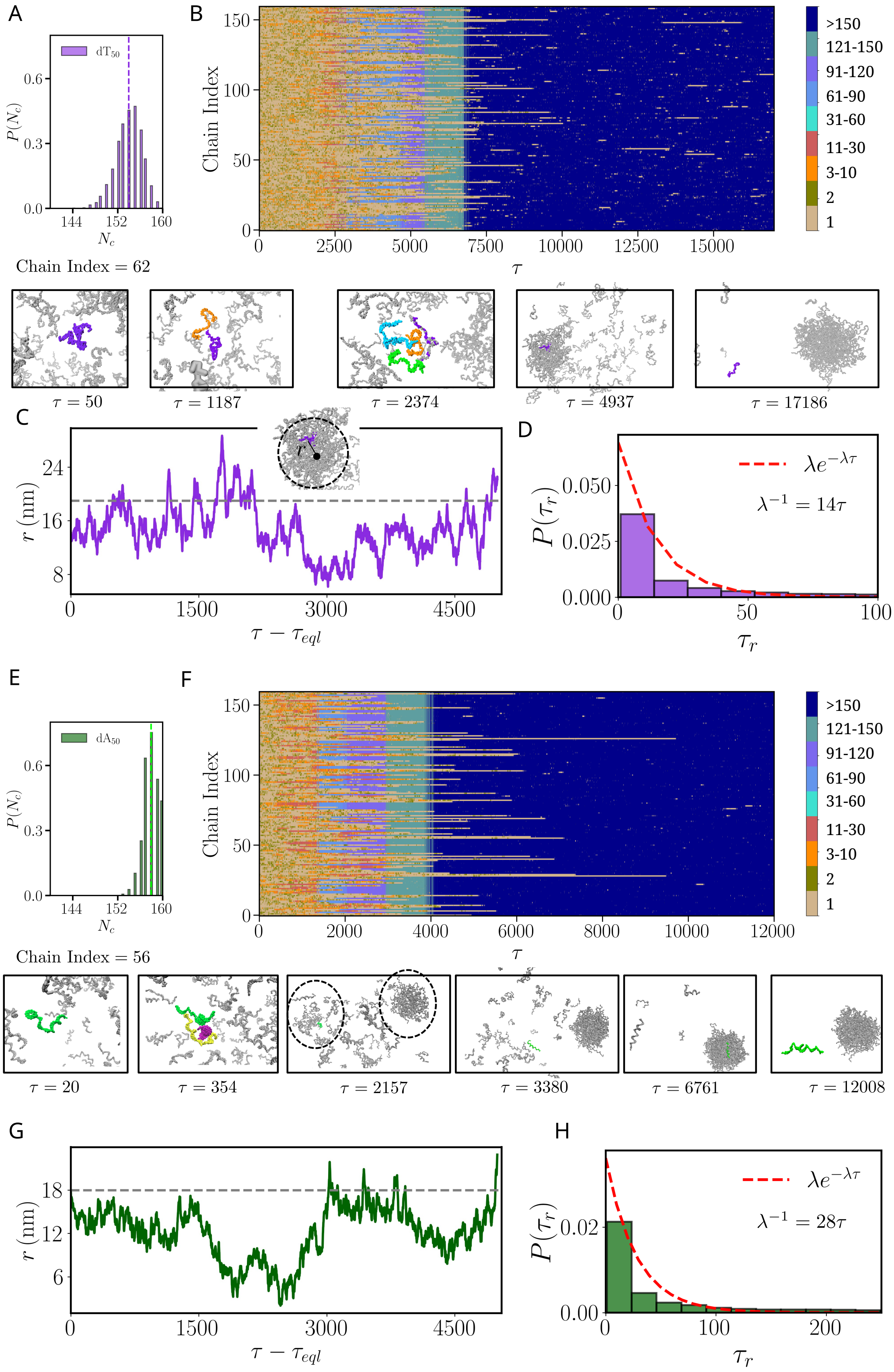




Figure S18: **Chain dynamics in condensates.**

**Figure S18 caption:** (A) Distribution ($P(N_c)$) of the number of chains $N_c$ in the condensate phase. Dashed line is the average value, $\langle N_c \rangle$. (B) Time dependent changes in individual chains during phase separation in $dT_{50}$. The chains are represented by separate lines. Colors show the oligomer sizes populated (scale on the right), determined using a clustering algorithm, during phase separation. Conformational states of a particular chain (highlighted in green) different times $\tau$ during phase separation are shown below. (C) Time evolution of distance ($r$) between the condensate center and center of mass of a labeled chain in the droplet. Dashed line is the the condensate radius. Inset shows how $r$ is calculated. (D) Distribution ($P(\tau_r)$) of the residence time ($\tau_r$) of the chains within the condensate. A chain is considered to be within the condensate if the center of mass distance ($r$) of the chain from the condensate center is less than or equal to the condensate radius. Red dashed line is an exponential fit to $P(\tau_r)$. Characteristic time ($\lambda^{-1}$) is reported. (E) Same as (A), except the results are for $dA_{50}$. (F) Same as (B), except the results are for $dA_{50}$. (G) Same as (C), except the results are shown for $dA_{50}$. (H) Same as (D), except the results are shown for $dA_{50}$. Results are shown for the largest system with 160 ssDNA chains, in 10 mM $Spm^{4+}$ .

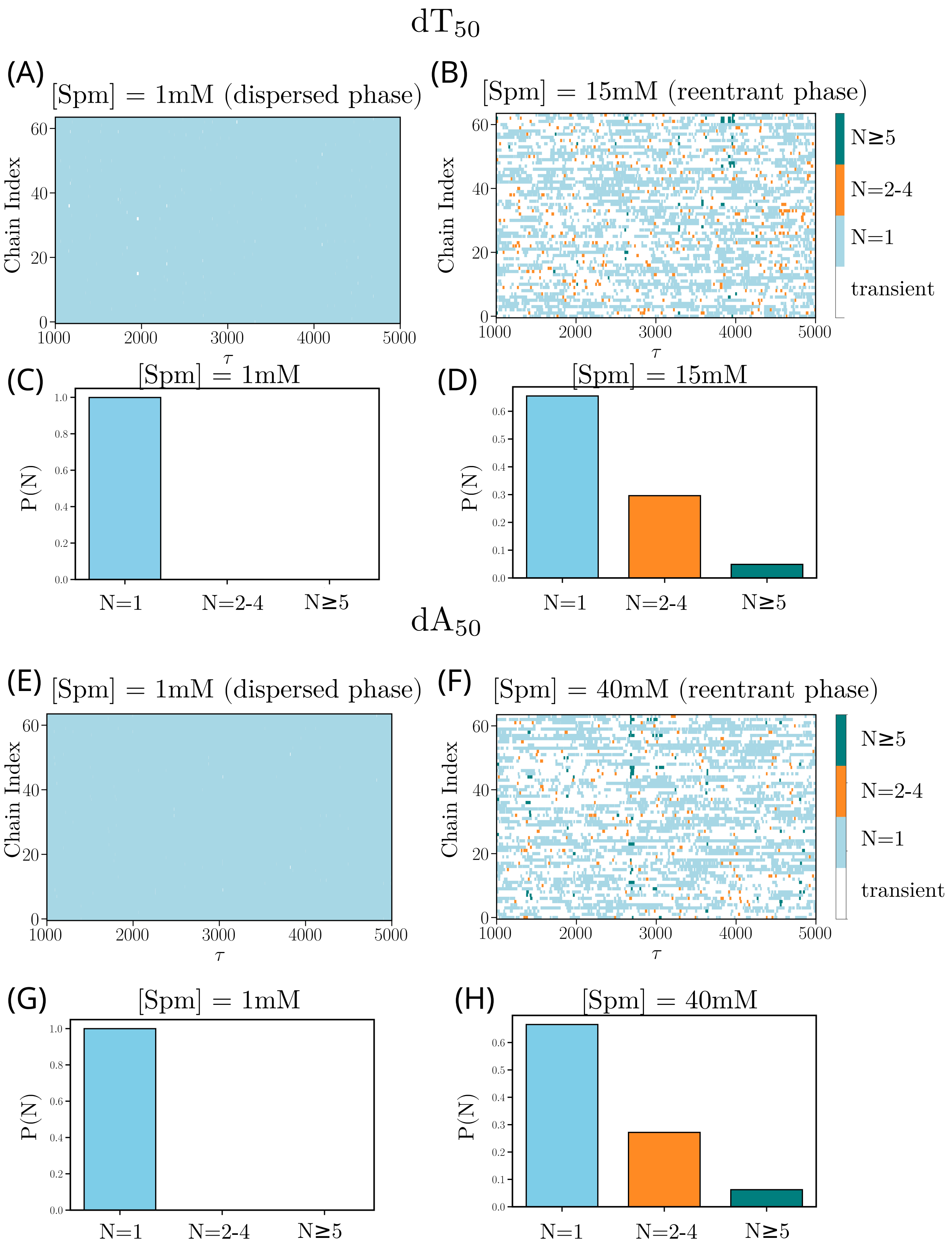
dT50
(A) [Spm] = 1mM (dispersed phase)
(B) [Spm] = 15mM (reentrant phase)
Chain Index
τ
N≥5
N=2-4
N=1
transient
(C) [Spm] = 1mM
(D) [Spm] = 15mM
P(N)
N=1
N=2-4
N≥5
dA50
(E) [Spm] = 1mM (dispersed phase)
(F) [Spm] = 40mM (reentrant phase)
(G) [Spm] = 1mM
(H) [Spm] = 40mM

Figure S19

**Comparing dispersed and reentrant phases:** (A) Chain-state evolution for all DNA chains in the dispersed phase of $dT_{50}$ in 1mM Spm. Only states that persist for at least $20\tau$ consecutive steps are color-coded. Shorter excursions are shown in white. Sustained monomers ($N = 1$), dimer–tetramers ($N = 2 - 4$), and oligomers ($N \geq 5$) are highlighted. (B) Same as (A), except results are shown for 15mM Spm. (C) Chain–state occupancies pooled over all chains ($dT_{50}$) and time frames, in 1mM Spm. (D) Same as (C), except the results are shown for 15mM Spm. (E) Same as (A), except the results are shown for $dA_{50}$. (F) Same as (E), except results are shown for 40mM Spm. (G) Same as (C), except the results are for $dA_{50}$. (H) Same as (G), except the results are for 40mM Spm.

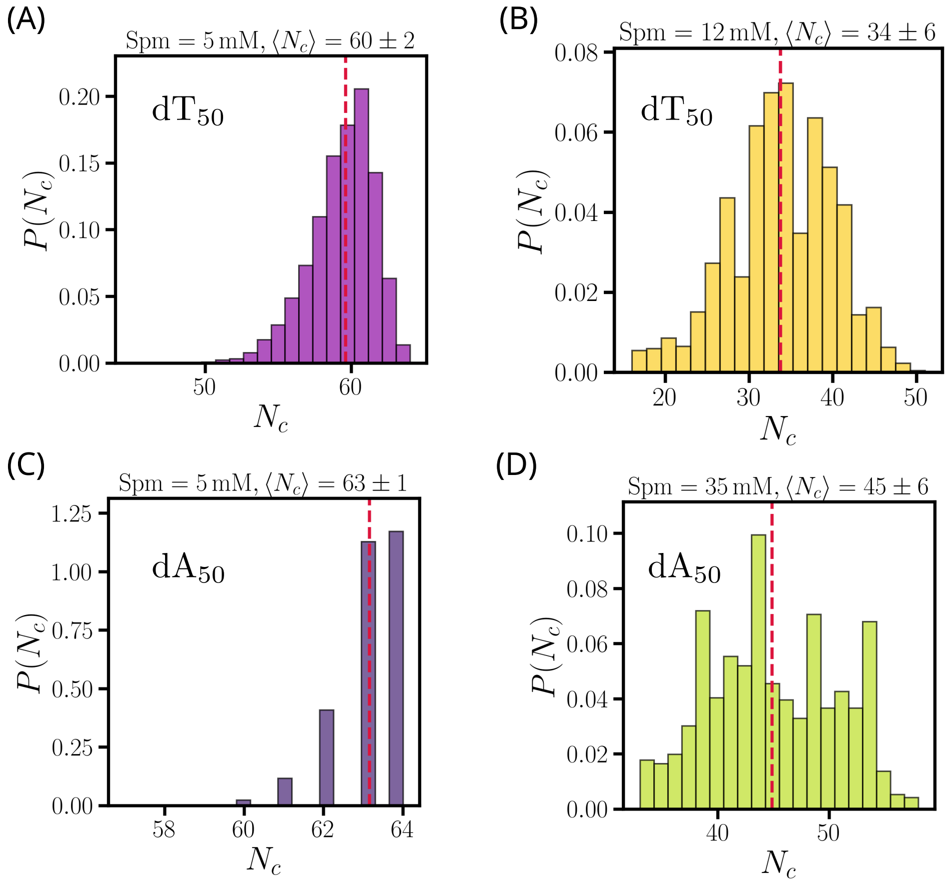


Figure S20: **Equilibrium aspects of condensed phases:** (A-D) Distributions of number of chains, $P(N_c)$ in the condensates at different $Spm^{4+}$ concentrations.

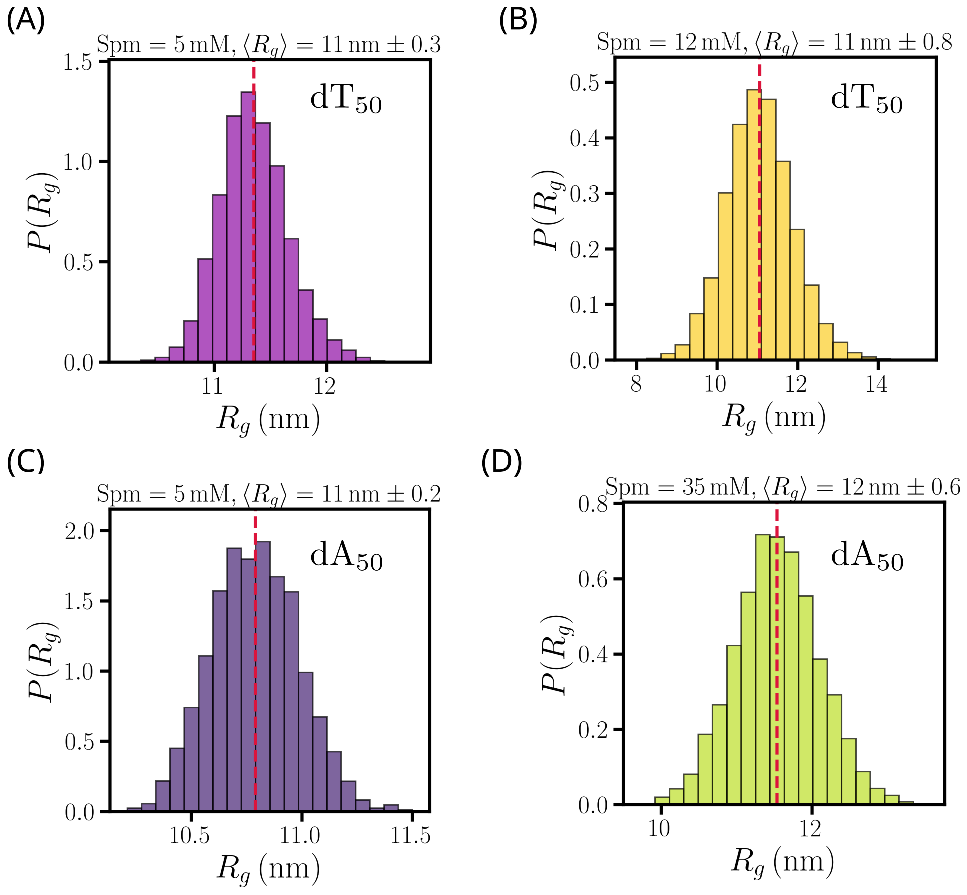


Figure S21: **Equilibrium characteristics of condensed phases:** (A-D) Distributions of radius of gyration of the condensates, $P(R_g)$, in different $Spm^{4+}$ concentrations.

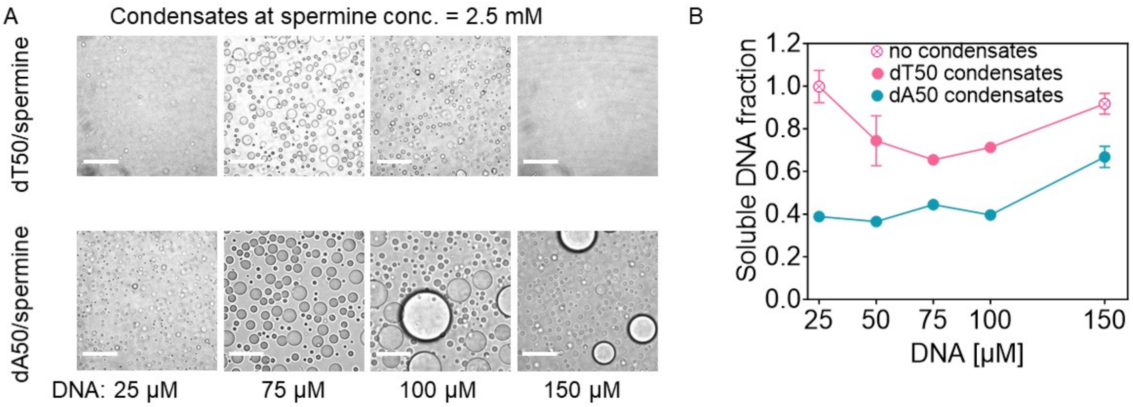


Figure S22: **ssDNA/spermine complex coacervates at varying DNA concentrations:** (A) Representative micrographs showing ssDNA/spermine complex coacervates and (B) soluble DNA fraction, at indicated DNA concentrations and 2.5 mM spermine. The complex coacervates were formed at 10 mM Tris pH 7.0 and at room temperature. Each data point in B represents mean $\pm$ SD of n = 9 measurements from 3 independent experiments. The scale bars represent 30 $\mu$m.

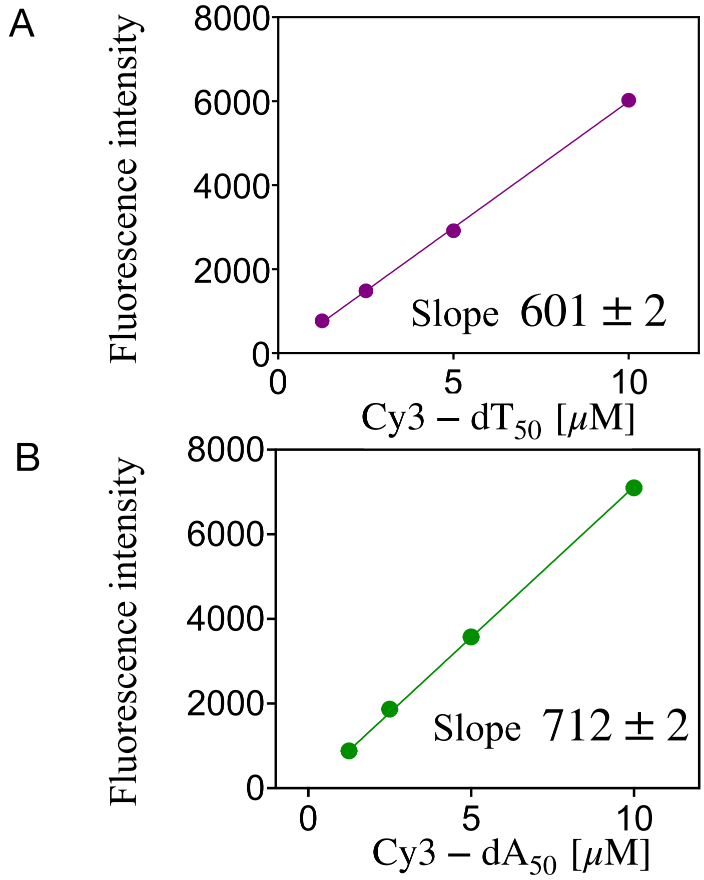


Figure S23: **Calibration curve of Cy3-**$dT_{50}$ **and Cy3-**$dA_{50}$**:** Plot showing fluorescence intensity of Cy3-$dT_{50}$ and Cy3-$dA_{50}$ scaled linearly with increasing DNA concentration.

**Supplementary Video 1. Condensate growth mechanism in** $dT_{50}$**.** Time-resolved simulation trajectory showing phase separation of $dT_{50}$ in 10 mM $Spm^{4+}$ ($N = 160$ chains, $T = 298$ K). The $dT_{50}$ condensate grows by sequential addition of individual chains to the growing droplet. This behavior is reflected in the nearly linear increase in cluster size, $N(t)$, and the constant growth rate, $dN(t)/dt$, shown in Figure 5C of the main text. Video is available at `https://zenodo.org/records/22212158`.

**Supplementary Video 2. Condensate growth mechanism in** $dA_{50}$**.** Phase separation in $dA_{50}$ in 10 mM $Spm^{4+}$ ($N = 160$ chains, $T = 298$ K). At short times, multiple small oligomers in $dA_{50}$. Subsequently they coalesce into a single, larger condensate, a growth mechanism reminiscent of Ostwald ripening. The abrupt, step-like increase in cluster size associated with a major fusion event is quantified in Figure 5D of the main text. The newly fused condensate is initially non-spherical but relaxes to a spherical droplet at longer times to minimize the interfacial free energy. Video is available at `https://zenodo.org/records/22212158`.

Table S1: Condensate ($\rho_c$) and dilute-phase ($\rho_d$) DNA concentrations (mM) for $dT_{50}$ and $dA_{50}$ at different spermine concentrations. Values are mean ± standard deviation.

| | $dT_{50}$ | | $dA_{50}$ | |
|---|---|---|---|---|
| Spm (mM) | $\rho_c$ (mM) | $\rho_d$ (mM) | $\rho_c$ (mM) | $\rho_d$ (mM) |
| 4.375 | $15.96 \pm 1.55$ | $0.04 \pm 0.02$ | $20.55 \pm 0.95$ | $0.01 \pm 0.01$ |
| 5 | $17.08 \pm 1.18$ | $0.02 \pm 0.01$ | $20.71 \pm 0.96$ | $0.00 \pm 0.00$ |
| 7 | $15.86 \pm 1.22$ | $0.05 \pm 0.02$ | $20.00 \pm 1.03$ | $0.01 \pm 0.01$ |
| 10 | $13.89 \pm 1.90$ | $0.09 \pm 0.03$ | $19.11 \pm 1.08$ | $0.02 \pm 0.01$ |
| 12 | $11.71 \pm 1.58$ | $0.15 \pm 0.02$ | $18.55 \pm 1.11$ | $0.02 \pm 0.01$ |
| 15 | | | $17.76 \pm 1.13$ | $0.02 \pm 0.01$ |
| 20 | | | $16.52 \pm 1.19$ | $0.03 \pm 0.01$ |
| 27 | | | $14.77 \pm 1.37$ | $0.06 \pm 0.02$ |
| 30 | | | $13.77 \pm 1.53$ | $0.08 \pm 0.03$ |
| 35 | | | $12.48 \pm 1.69$ | $0.11 \pm 0.04$ |

Table S2: Parameters for excluded volume and electrostatic interactions for DNA beads and ions. $R_i$: radius, $\epsilon_i$: interaction strength for volume exclusion, $q_i$: charge of $i^{th}$ species. P represents phosphate groups, S represents sugar groups, A, G, T, C represent nucleobase groups, and $Na^+$, $Cl^-$ are monovalent ions.

| | $R_i$ nm | $\epsilon_i$ kcal/mol | $q_i$ e |
|---|---|---|---|
| P | 0.21 | 0.2 | -1 |
| S | 0.29 | 0.2 | 0 |
| A | 0.28 | 0.2 | 0 |
| G | 0.3 | 0.2 | 0 |
| T | 0.27 | 0.2 | 0 |
| C | 0.27 | 0.2 | 0 |
| $Na^+$ | 0.19 | 0.00277 | 1 |
| $Cl^-$ | 0.19 | 0.3 | -1 |
| $NH^+$ | 0.163 | 0.17 | +1 |
| $(CH_2)_3$ | 0.293 | 0.17 | 0 |
| $(CH_2)_4$ | 0.308 | 0.17 | 0 |

Table S3: Parameters for non-neighboring stacking interactions (see Eq. S1).

| stacked bases | $d_{0,0}$ nm | $d_{1,0}$ nm | $d_{2,0}$ nm | $\theta_{1,0}$ radians | $\theta_{2,0}$ radians | $\psi_{0,syn}$ radians | $\psi_{0,anti}$ radians |
|---|---|---|---|---|---|---|---|
| $\frac{T}{T}$ | 4.018 | 6.7 | 6.7 | 1.679 | 1.679 | 0.58 | 2.683 |
| $\frac{A}{A}$ | 3.74 | 6.5 | 6.5 | 1.679 | 1.679 | 0.575 | 2.683 |

$k_d^{NN} = 5.0$ Å$^{-2}$, $k_\theta^{NN} = 5.0$ rad$^{-2}$, $k_\psi^{NN} = 5.0$ rad$^{-4}$;

Table S4: Parameters to compute neighboring stacking energy (Eq. 19 of main text),[31] $U^0_{ST} = -h + k_\mathrm{B}(T - T_m)s$.

| $\frac{x}{w}$ | $h$ (kcal/mol) | $s$ | $T_m$ (K) |
|---|---|---|---|
| $\frac{A}{A}$ | 5.69 | 0.94 | 322.0 |
| $\frac{A}{T}$; $\frac{T}{A}$ | 4.95; 5.03 | 0.87; 0.65 | 293.0 |
| $\frac{A}{G}$; $\frac{G}{A}$ | 5.43; 5.42 | 1.06; 0.98 | 333.6 |
| $\frac{A}{C}$; $\frac{C}{A}$ | 4.98; 5.03 | 0.92; 0.79 | 293.0 |
| $\frac{G}{G}$ | 5.66 | −0.29 | 353.9 |
| $\frac{G}{C}$; $\frac{C}{G}$ | 5.22; 5.13 | 2.33; 2.41 | 331.9 |
| $\frac{G}{T}$; $\frac{T}{G}$ | 5.28; 5.13 | 1.69; 1.73 | 332.6 |
| $\frac{C}{C}$ | 4.15 | 0.98 | 288.3 |
| $\frac{C}{T}$; $\frac{T}{C}$ | 4.13; 4.18 | 0.71; 0.94 | 288.3 |
| $\frac{T}{T}$ | 4.17 | 0.89 | 288.3 |

$k_d = 1.45$ Å$^{-2}$, $k_\psi = 3.0$ rad$^{-2}$;

### References


(1) Jhunjhunwala, A.; Ali, Z.; Bhattacharya, S.; Halder, A.; Mitra, A.; Sharma, P. On the nature of nucleobase stacking in RNA: a comprehensive survey of its structural variability and a systematic classification of associated interactions. *J. Chem. Inf. Model.* **2021**, *61*, 1470–1480.

(2) Mitra, R.; Cohen, A. S.; Sagendorf, J. M.; Berman, H. M.; Rohs, R. DNAproDB: an updated database for the automated and interactive analysis of protein–DNA complexes. *Nucleic Acids Res.* **2025**, *53*, D396–D402.

(3) Mondal, B.; Chakraborty, D.; Hori, N.; Nguyen, H. T.; Thirumalai, D. Competition between Stacking and Divalent Cation-Mediated Electrostatic Interactions Determines the Conformations of Short DNA Sequences. *J. Chem. Theory Comput.* **2024**, *20*, 2934–2946.

(4) Eastman, P.; Swails, J.; Chodera, J. D.; McGibbon, R. T.; Zhao, Y.; Beauchamp, K. A.; Wang, L. P.; Simmonett, A. C.; Harrigan, M. P.; Stern, C. D.; Wiewiora, R. P.; Brooks, B. R.; Pande, V. S. OpenMM 7: Rapid development of high performance algorithms for molecular dynamics. *PLOS Biol.* **2017**, *13*, e1005659.

(5) Darden, T.; York, D.; Pedersen, L. Particle mesh Ewald: An Nlog(N) method for Ewald sums in large systems. *J. Chem. Phys.* **1993**, *98*, 10089–10092.

(6) Honeycutt, J. D.; Thirumalai, D. The nature of folded states of globular proteins. *Biopolymers* **1992**, *32*, 695–709.

(7) Hockney, R. W.; Eastwood, J. W. *Computer simulation using particles*; CRC Press, 2021.

(8) Thirumalai, D.; Ha, B.-Y. In *Theoretical and Mathematical Models in Polymer Research*; Grosberg, A., Ed.; Academic Press: San Diego, 1998; Chapter I. Statistical

Mechanics of Semiflexible Chains: A Mean Field Variational Approach, pp 1–35, edited by A. Grosberg.

(9) Hyeon, C.; Thirumalai, D. Kinetics of interior loop formation in semiflexible chains. *J. Chem. Phys.* **2006**, *124*, 104905.

(10) Klimov, D. K.; Betancourt, M. R.; Thirumalai, D. Virtual atom representation of hydrogen bonds in minimal off-lattice models of alpha helices: Effect on stability, cooperativity and kinetics. *Folding and Design* **1998**, *3*, 481–496.

(11) Moradi, M.; Babin, V.; Roland, C.; Darden, T. A.; Sagui, C. Conformations and free energy landscapes of polyproline peptides. *Proc. Natl. Acad. Sci. USA* **2009**, *106*, 20746–20751.

(12) Theodorou, D. N.; Suter, U. W. Shape of unperturbed linear polymers: polypropylene. *Macromolecules* **1985**, *18*, 1206–1214.

(13) Hyeon, C.; Dima, R. I.; Thirumalai, D. Size, shape, and flexibility of RNA structures. *J. Chem. Phys.* **2006**, *125*.

(14) Nguyen, H. T.; Hori, N.; Thirumalai, D. Condensates in RNA repeat sequences are heterogeneously organized and exhibit reptation dynamics. *Nature Chem.* **2022**, *14*, 775–785.

(15) Muthukumar, M. Theory of counter-ion condensation on flexible polyelectrolytes: Adsorption mechanism. *The Journal of chemical physics* **2004**, *120*, 9343–9350.

(16) Ha, B.-Y.; Thirumalai, D. Bending rigidity of stiff polyelectrolyte chains: A single chain and a bundle of multichains. *Macromolecules* **2003**, *36*, 9658–9666.

(17) Hori, N.; Denesyuk, N. A.; Thirumalai, D. Ion condensation onto ribozyme is site specific and fold dependent. *Biophysical Journal* **2019**, *116*, 2400–2410.

(18) Majumder, S.; Coupe, S.; Fakhri, N.; Jain, A. Sequence-encoded intermolecular base pairing modulates fluidity in DNA and RNA condensates. *Nat. Commun.* **2025**, *16*, 1–12.

(19) Wang, Z.; Lou, J.; Zhang, H. Essence determines phenomenon: Assaying the material properties of biological condensates. *J. Biol. Chem.* **2022**, *298*, 101782.

(20) Baggett, D. W.; Medyukhina, A.; Tripathi, S.; Shirnekhi, H. K.; Wu, H.; Pounds, S. B.; Khairy, K.; Kriwacki, R. An image analysis pipeline for quantifying the features of fluorescently-labeled biomolecular condensates in cells. *Front. bioinform.* **2022**, *2*, 897238.

(21) McCall, P. M.; Kim, K.; Shevchenko, A.; Ruer-Gruß, M.; Peychl, J.; Guck, J.; Shevchenko, A.; Hyman, A. A.; Brugués, J. A label-free method for measuring the composition of multicomponent biomolecular condensates. *Nature Chem.* **2025**, *17*, 1891–1902.

(22) Nguyen, B.; Ciuba, M. A.; Kozlov, A. G.; Levitus, M.; Lohman, T. M. Protein environment and DNA orientation affect protein-induced Cy3 fluorescence enhancement. *Biophys. J.* **2019**, *117*, 66–73.

(23) Morten, M. J.; Lopez, S. G.; Steinmark, I. E.; Rafferty, A.; Magennis, S. W. Stacking-induced fluorescence increase reveals allosteric interactions through DNA. *Nucleic Acids Res.* **2018**, *46*, 11618–11626.

(24) Widom, B. Some topics in the theory of fluids. *J. Chem. Phys.* **1963**, *39*, 2808–2812.

(25) Shing, K.; Gubbins, K. The chemical potential from computer simulation: Test particle method with umbrella sampling. *Mol. Phys.* **1981**, *43*, 717–721.

(26) Qin, S.; Zhou, H.-X. Fast method for computing chemical potentials and liquid–liquid phase equilibria of macromolecular solutions. *J. Phys. Chem. B* **2016**, *120*, 8164–8174.

(27) Kumar, S. K.; Szleifer, I.; Panagiotopoulos, A. Z. Determination of the chemical potentials of polymeric systems from Monte Carlo simulations. *Phys. Rev. Lett.* **1991**, *66*, 2935.

(28) de Pablo, J. J.; Laso, M.; Suter, U. W. Estimation of the chemical potential of chain molecules by simulation. *J. Chem. Phys.* **1992**, *96*, 6157–6162.

(29) Bakhshandeh, A.; Levin, Y. Widom insertion method in simulations with Ewald summation. *J. Chem. Phys.* **2022**, *156*.

(30) Meisburger, S. P.; Sutton, J. L.; Chen, H.; Pabit, S. A.; Kirmizialtin, S.; Elber, R.; Pollack, L. Polyelectrolyte properties of single stranded DNA measured using SAXS and single-molecule FRET: Beyond the wormlike chain model. *Biopolymers* **2013**, *99*, 1032–1045.

(31) Chakraborty, D.; Hori, N.; Thirumalai, D. Sequence-Dependent Three Interaction Site Model for Single- and Double-Stranded DNA. *J. Chem. Theory Comput.* **2018**, *14*, 3763–3779.
